# Anti-amyloid immunotherapy drives APOE4 specific increases in glial reactivity, perivascular immune activation, and ARIA-like events

**DOI:** 10.64898/2026.06.26.734793

**Authors:** Akhil V Pallerla, Chloe C Lucido, Kai Saito, Georgia L Nolt, Jose Arbones-Mainar, Jessica L Funnell, Diksha Satish, Lily M Smith, Isaiah O Stephens, Danielle Goulding, Steven M MacLean, Samantha M Olmsted, Darcy Adreon, Gabriela Hernandez, Lesley R Golden, Scott Persohn, Shannon L Macauley, Paul R Territo, Josh M Morganti, Lance A Johnson

## Abstract

Anti-amyloid antibodies represent the first disease modifying therapeutics for Alzheimer’s disease (AD). Adoption of these novel treatments has been slowed by the occurrence of amyloid related imaging abnormalities (ARIA) – treatment-associated edema (ARIA-E) or microhemorrhages (ARIA-H) that disproportionately affect carriers of the E4 allele of apolipoprotein E (*APOE)*. With E4 carriers comprising nearly 70% of the AD population, there is a critical need to understand the unique vulnerability of E4 carriers to these events. To address this gap, we utilized the EFAD mouse model – which expresses human APOE isoforms on the 5xFAD background of amyloidosis – to directly compare the effects of anti-amyloid therapy across *APOE* genotypes. 9-month-old E2, E3, and E4FAD mice received weekly injections of chimeric Aducanumab (chAdu) or IgG control for 12 weeks, to assess APOE isoform-specific effects on amyloid dynamics, ARIA-H-like microhemorrhages, and underlying cellular and transcriptomic responses. E4FAD mice demonstrated plaque reductions with accompanying increases in microhemorrhages (measured on both MRI and histology), and increases in microglial and astrocyte reactivity – especially in the perivascular compartment. Additionally, vascular branching analysis and parallel single cell and spatial transcriptomics revealed a loss of vascular plasticity and increased inflammatory and immune signaling in the neurovascular units of E4FAD mice. Together, these findings suggest the cerebrovasculature of E4s is uniquely susceptible to antibody mediated vascular damage and provide immunological targets for the assessment or mitigation of ARIA risk in this highest need population

## INTRODUCTION

Alzheimer’s disease (AD) affects over 55 million individuals worldwide, with this number set to increase exponentially in the coming decades (*1*). Until recently, treatment for AD was limited to symptomatic management; the FDA approval of anti-amyloid (anti-Aβ) antibody treatments, however, now provides the first disease-modifying therapeutics for AD. These antibodies produce robust Aβ clearance with a modest – but clinically meaningful – slowing of cognitive decline (*2–4*). Adoption of these therapies has been tempered by the risk of amyloid-related imaging abnormalities (ARIA) – findings of vasogenic edema (ARIA-E) or microhemorrhage and hemosiderin deposition (ARIA-H) detected on MRI following treatment (*5*).

ARIA risk is strongly modulated by apolipoprotein E (*APOE*) genotype. *APOE* exists in three common allelic variants in the human population: E2, E3, and E4. Among these, E2 is protective against AD, E3 is considered a neutral risk allele, and E4 drives a 2-15-fold increase in AD risk depending on allelic dose (*6–8*). Critically, nearly 70% of all AD patients carry at least one E4 allele, and ARIA rates in E4 homozygotes are nearly triple those of non-carriers (*2–4, 6–8*). The need to understand ARIA risk is emphasized by the most extreme outcomes: three published ARIA-related deaths in clinical trials, all of which occurred in homozygous APOE4 carriers (*9–11*). The over-representation of E4 carriers in the AD population, coupled with the higher rate and severity of ARIA in these individuals highlights the critical need to understand the underlying mechanisms of ARIA. In addition to the over-representation of E4 carriers in the AD population, they are also disproportionately affected by cerebral amyloid angiopathy (CAA), vascular deposits of Aβ, another key modulator of ARIA risk. CAA alone has been shown to drive ARIA-like inflammatory events, making it important to dissect the relative contributions of elevated levels of CAA and other E4 mediated vascular or immune dysfunction(*5, 12*).

Preclinical studies of anti-amyloid antibodies have largely been conducted in models of amyloidosis that express murine *Apoe* (*13–20*), which shares only ∼70% sequence homology with human *APOE* and has been shown to drive a more aggressive amyloid phenotype than human E4 (*21–24*). This is a critical limitation, as murine ApoE cannot recapitulate the isoform-specific biology that underlies differential ARIA risk in humans. While a small number of studies have examined anti-amyloid therapy in the context of human APOE isoforms these studies primarily focused on treatment efficacy and quantification of ARIA-like events without in-depth exploration of potential mechanistic drivers at the cell/tissue levels (*25–27*). This presents a key gap in understanding, both in terms of the role that the different human APOE isoforms play in treatment efficacy, and the role that E4 plays in ARIA pathogenesis.

To address this gap, we leveraged the EFAD mouse model, in which human APOE isoforms are expressed on the amyloidogenic 5xFAD background, to systematically characterize isoform-specific responses to anti-amyloid therapy (*22, 23*). Nine-month-old E2FAD, E3FAD, and E4FAD mice were treated for 12 weeks with a low dose of chimeric Aducanumab (chAdu), a validated preclinical surrogate for clinical anti-amyloid antibodies (*26, 28*). We demonstrate that chAdu-driven plaque clearance in E4FAD mice is accompanied by increased ARIA-H-like events detected by susceptibility-weighted MRI and Prussian Blue staining, E4-specific increases in peri-plaque and perivascular astro- and microgliosis, and isoform-specific differences in vascular remodeling responses. To further characterize the cellular mechanisms underlying these findings, single-cell RNA sequencing and Xenium spatial transcriptomics were performed on a subset of E4FAD mice, revealing treatment-induced alterations in microglial subpopulations and spatial reorganization of the perivascular niche. Together, these findings provide mechanistic insight into the role of *APOE4* in driving ARIA after anti-amyloid immunotherapy, laying the groundwork for targeted strategies to improve treatment safety in E4 carriers.

## RESULTS

### Aducanumab reduces parenchymal and vascular Aβ in E4FAD mice

To validate the use of the EFAD mouse model as a translationally relevant system for studying anti-amyloid antibodies and therapies, we employed a treatment paradigm validated by MODEL-AD, where E2FAD, E3FAD and E4FAD mice were treated for 12 weeks with 1.56 mg/kg/week of chAdu or IgG isotype control (Fig 1A)(*29*). Whole brain quantification of AmyloGlo staining area showed that, as previously reported, there are genotype specific differences in plaque burden mimicking human pathology in EFAD mice (E4>E3>E2) (*23, 24, 30–34*). After treatment with chAdu, there was a modest, but significant, reduction in plaque burden in E4FAD mice only. We also noted *APOE* genotype differences in plaque size (E4>E3, E2), although all groups were unaffected by chAdu treatment (Fig 1B-D).

**Figure 1.**
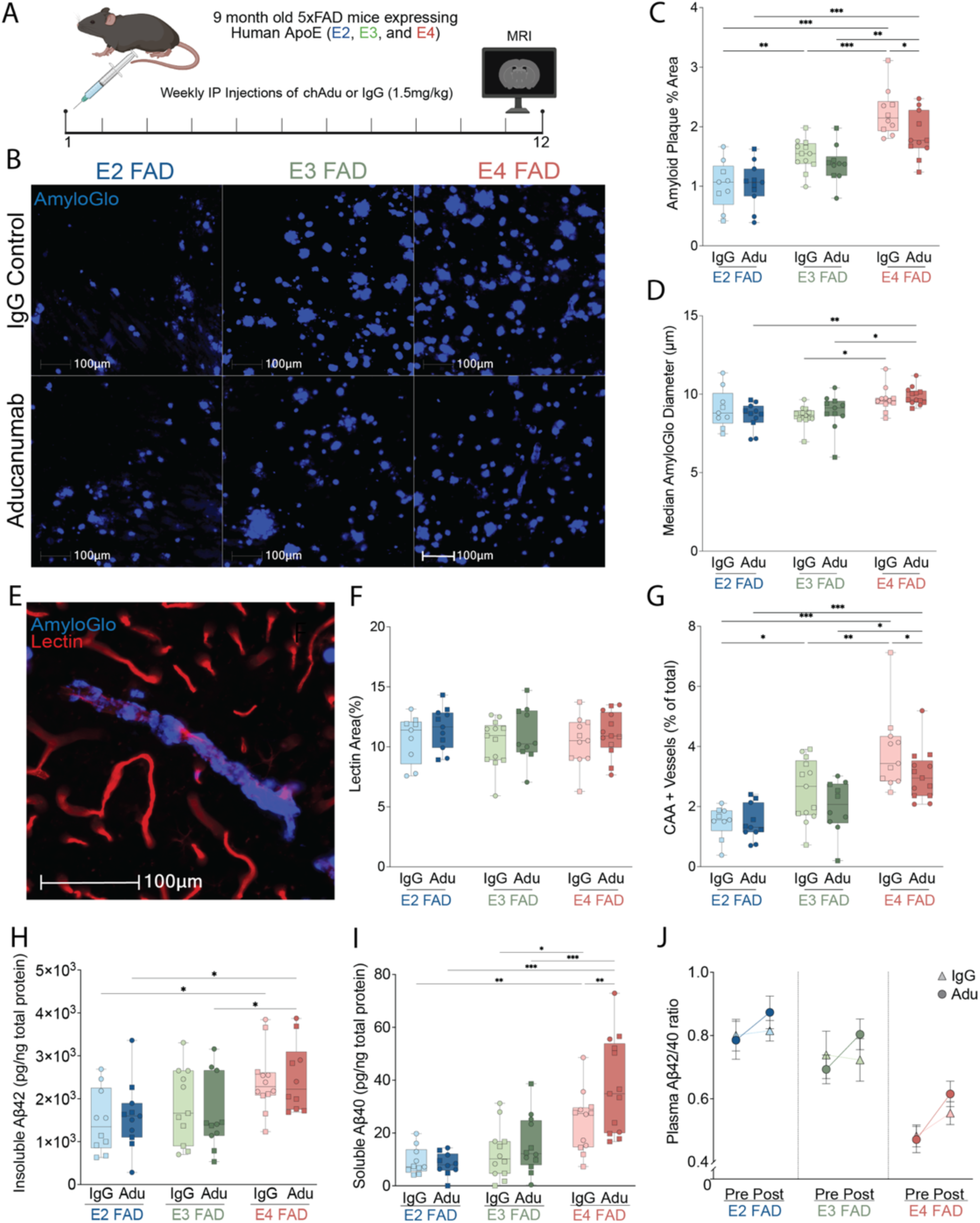
Anti-amyloid antibody treatment reduced parenchymal and vascular Aβ in E4FAD mice. (A) Experimental design and treatment paradigm. EFAD mice of each genotype (n=5 minimum per sex/genotype/treatment) were treated with a chimeric Aducanumab weekly for 12 weeks. MRI scan was conducted one day prior to sacrifice and tissue collection. (B) Representative images of AmyloGlo staining used to quantify plaque burden across animals. (C) Histological quantification of amyloid plaque burden across genotypes. Males are denoted by a square and females with circles. (D) Quantification of median plaque diameter in µm. (E) Representative image of AmyloGlo staining overlapped with isolectin vascular labeling. (F) Quantification of total area coverage of isolectin positive vessels (G) Quantification of colocalization of amyloglo and isolectin vascular staining as percent of isolectin. (H-J) ELISA quantification of brain insoluble Aβ42, soluble Aβ40, and plasma Aβ42/40 ratio, respectively. 2-way ANOVA with Tukey’s post-hoc comparisons(p-value * <.05, ** <.005 *** <.001), specific n’s and p-values used in each analysis are included in supplementary data file 1.

As elevated levels of CAA in E4 carriers have been identified as a potential driver of ARIA, we quantified CAA burden through colocalization of AmyloGlo with isolectin, a vascular marker (*5*). While there were no differences in total vascular area by genotype or treatment, we observed genotype specific differences in CAA (E4>E3>E2) and a modest, but significant, reduction in E4FAD mice only following treatment (Fig 1E-G). This reduction in CAA occurs regardless of region of interest (Supp. Fig. 1B-D).

To further assess amyloid dynamics, we measured Aβ40 and Aβ42 in brain tissue and plasma by ELISA. In the brain, soluble Aβ42 did not differ across genotypes or treatments, but insoluble Aβ42 showed a significant genotype effect, with E4FAD mice having higher levels than other genotypes (Fig 1H, Supp Fig 2A). Aβ40 fractions showed significant genotype effects without a significant overall treatment response, although chAdu-treated E4FAD mice had significantly higher soluble Aβ40 than IgG-treated E4FAD mice (Fig 1I, Supp Fig 2B). In plasma, both Aβ40 and Aβ42 changed significantly over time – with Aβ40 declining in E4FAD mice and Aβ42 declining in E2FAD mice – though no significant time-by-treatment interaction was detected (Supp Fig 2C-F). The plasma Aβ42/40 ratio showed significant effects of both time and treatment group: chAdu-treated E4FAD mice showed significant increases in this ratio while IgG-treated E4FAD mice did not. (Fig 1J, Supp Fig 2G). Taken together, while individual Aβ fractions reflect primarily genotype-driven differences, the plasma Aβ42/40 ratio shows the clearest treatment-associated differences particularly in E4FAD mice –suggesting a possible genotype-dependent relationship between chAdu treatment and amyloid dynamics.

**Figure 2:**
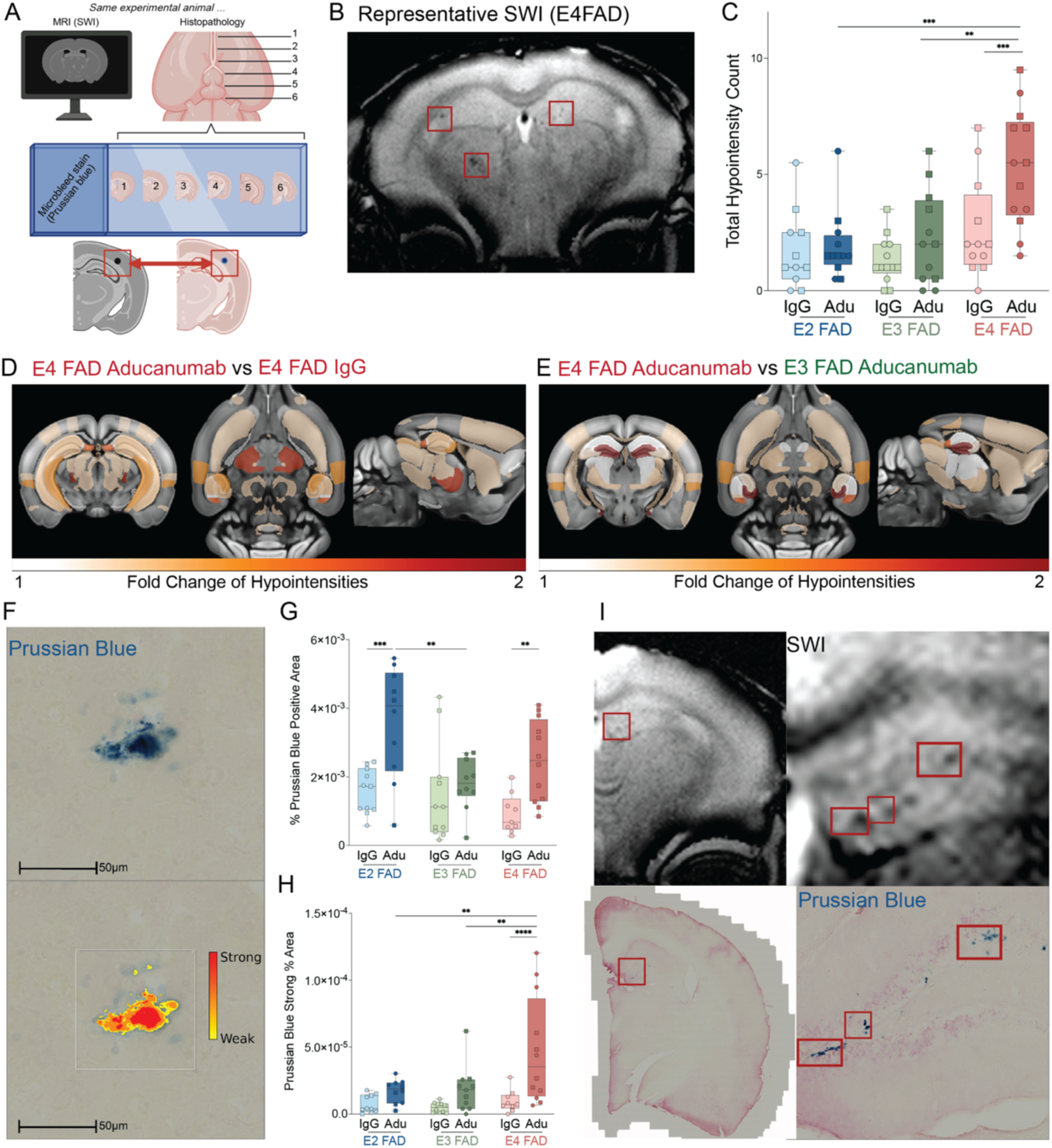
Aducanumab treatment increases microbleed occurrence in E4FADs, seen with MRI and Prussian Blue staining. (A) Experimental design of MRI and Prussian Blue quantification, as well as cross-measure alignment validation. (B) Representative SWI MRI with bleeds indicated by red boxes. (C) Quantification of hypointense regions representing bleeds. (D-E) AtlasModeler representation of bleed changes between Aducanumab and IgG treated E4FAD mice and Aducanumab treated E3 and E4FAD mice. (F) Representative Prussian Blue bleed (top) with quantification algorithm (bottom). Breakdown of bleed intensities are included in supplementary figure 2. (G) Quantification of Prussian Blue positive area per animal. (H) Quantification of strong Prussian Blue area (deep red). (I) Representative MRI (top) and Prussian Blue (bottom) overlap with zoomed in regions. 2-way ANOVA with Tukey’s post-hoc comparisons (p-value * <.05, ** <.005 *** <.001), specific n’s and p-values used in each analysis are included in supplementary data file 2.

### Aducanumab drives increased ARIA-H like events in E4FAD mice

We next wanted to understand the role of *APOE* on incidence of ARIA-like events in EFAD mice. Because ARIA is an MRI-based phenomenon, all mice underwent MRI scanning one day prior to tissue collection and subsequent histopathological confirmation (Fig 2A). Susceptibility-weighted (SWI) and T2 imaging sequences were registered to the Allen Common Coordinate Framework reference atlas, and were then used to assess the incidence and location of ‘ARIA-H like’ events, with resultant values compared across genotypes and treatments in each region. We observed a significant increase in MRI hypointensities in E4FAD mice after treatment (Fig 2B), and significantly more hypointensities in E4FAD mice than E2 and E3 mice after chAdu treatment (Fig 2C), a finding that mimics clinical ARIA incidence (*2–4*).

To determine if there was regional bias in hypointensity occurrence, we used atlas-aligned SWI images to compare regions that were significantly altered across genotype and/or treatment. From this, we generated heatmaps of where bleeds increased the most in E4FAD mice treated with chAdu (compared to IgG; Fig 2D or compared to E3FAD; Fig 2E). When compared to E3FAD mice, there was a 1.7x increase in bleeds in the dentate gyrus (DG) and 1.4x increase in bleeds in the subiculum of chAdu treated E4FAD mice. We also noted changes in the subiculum of E4 chAdu treated mice when comparing with IgG treated mice (fold change 1.4), as well as smaller changes in CA1 and CA3, suggesting hippocampal susceptibility to treatment-associated vascular dysfunction.

For histological confirmation of ARIA-H like events, six evenly spaced anterior to posterior sections were stained with Prussian Blue to label microhemorrhages (Fig 2A). Interestingly, quantification of Prussian Blue bleeds showed significant increases in microbleed occurrence in both E2FAD mice and E4FAD mice after treatment with chAdu (Fig 2F-G). In addition to quantifying total microbleed area, we were further able to bin and quantify areas of weak to strong signal intensity (Fig 2H, Supp. Fig 3A-B). When comparing the relative distribution of weak, moderate, and strong signal intensity bleeds we noted similarities in distribution between the “strong” bleeds and SWI quantification, suggesting that only the strongest signal intensity bleeds will appear on MRI. In fact, we were able to spatially align the location of these “strong” signal microhemorrhages to their corresponding SWI hypointensities (Fig 2I).

**Figure 3:**
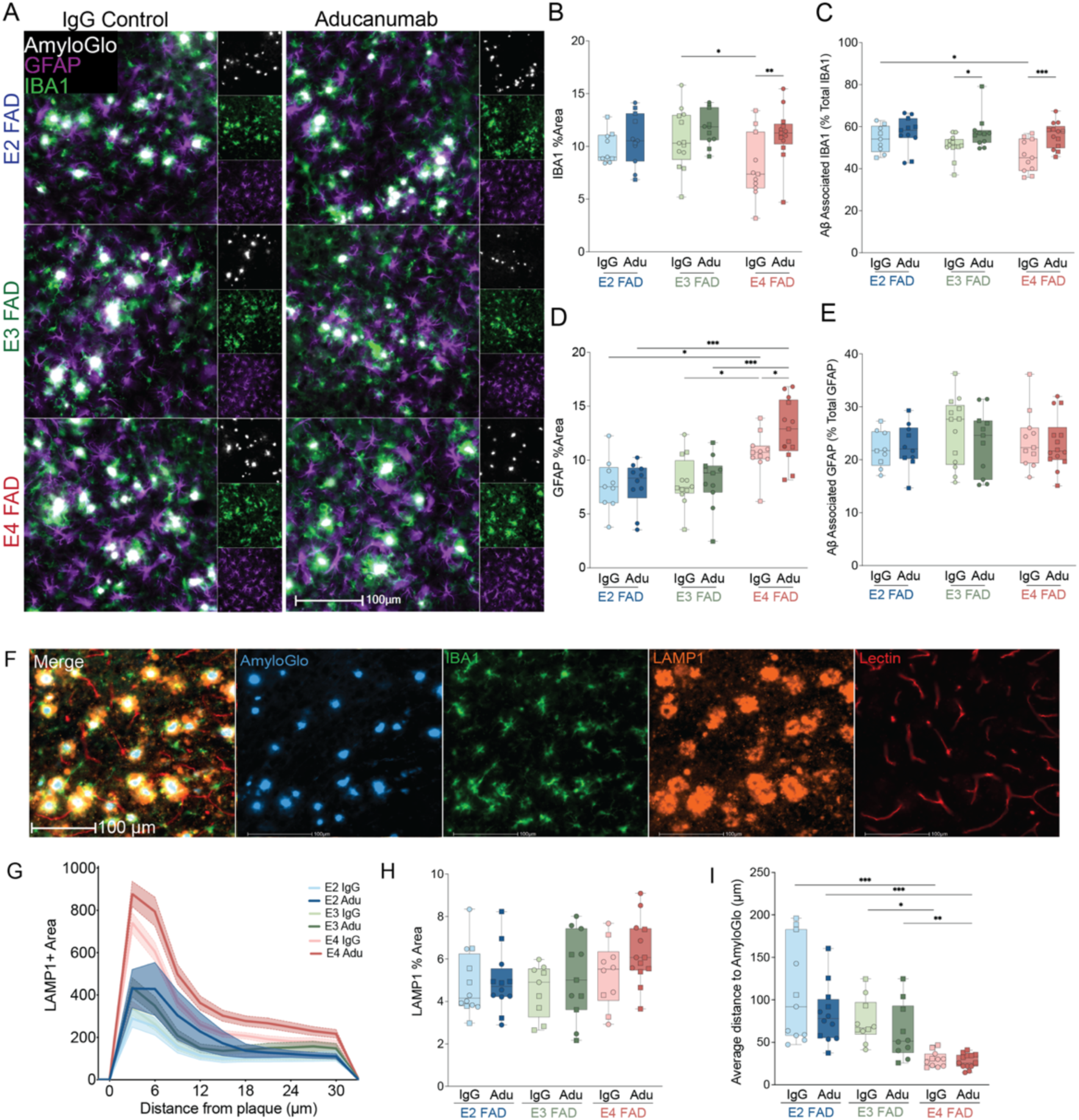
Aducanumab treatment drives E4-specific increases in overall and peri-plaque gliosis. (A) Representative images of IBA1 (green), GFAP (purple), and AmyloGlo (white). (B-C) Quantification of overall IBA1+ area across genotypes and peri-plaque IBA1 (as %Total IBA1). (D-E) Quantification of total GFAP+ area and peri-plaque GFAP (as % of total GFAP). (F) Representative images of AmyloGlo (blue), IBA1 (green), LAMP1 (orange), and Lectin (red). (G) Histogram of proximity between LAMP1 and amyloid plaques. (H) Total LAMP1% area coverage. (I) Average distance between LAMP1+ area and plaque. (2-way ANOVA with Tukey’s post-hoc comparisons(p-value * <.05, ** <.005 *** <.001), specific n’s and p-values for each analysis are included in supplementary data file 3

### Aducanumab treatment induces widespread micro-and astrogliosis in E4FAD mice

To elucidate mechanisms underlying ARIA-like events in E4FAD mice, we next stained serial sections for various markers of glial reactivity. chAdu treatment drove increases in overall IBA1 area in E4FAD mice only, as well as increases in peri-plaque microgliosis in both E3 and E4FAD mice (Fig 3A-C). There was also E4 specific increase in overall GFAP positive area, although there were no changes in plaque associated GFAP (Fig 3D-E). These changes mirror those seen in recent reports examining the effects of anti-amyloid therapy on the CNS and may indicate a potentially maladaptive immune response mechanism in E4s that may be driving ARIA events during antibody-mediated plaque clearance(*26, 28*).

As there is noted plaque associated dystrophy in E4 carriers, we also stained for LAMP1 in conjunction with IBA1, to determine if the increase in peri-plaque microglia is associated with a change in dystrophic neurites(*35*). While there were no significant changes in overall LAMP1 area, there was a genotype specific decrease in average distance from LAMP1 to AmyloGlo, with E4FAD mice having significantly lower distance to plaques than both E2FAD mice and E3FAD mice. In addition, LAMP1 was more closely colocalized to amyloid plaques, with a significant increase in LAMP1 positivity within 10 microns of plaque (Fig 3F). This finding aligns with other studies suggesting that there is a limited treatment effect on neuritic dystrophy, although there are significant genotypic drivers of plaque-associated neuritic dystrophy in E4FAD mice especially (*28, 35, 36*).

### Aducanumab treatment drives activated microglial states in E4FAD mice

To further evaluate whether there are changes in microglial states after chAdu treatment in E4FAD mice, we next stained for markers of homeostatic and activated microglia (P2ry12 and CD68 respectively). There were no changes in P2ry12 positive homeostatic microglia across genotypes or treatment, though we observed an increase in CD68+ microglia in E4FAD mice after treatment (Fig 4A-CD). There were no peri-plaque changes in CD68 in chAdu treated E4FAD mice, which may suggest plaque independent microglial activation (Fig 4D). This data appears to suggest that in the context of chAdu treatment in E4FAD mice, microglia and CNS macrophages are becoming more activated to drive immune mediated phagocytosis of Aβ throughout the brain.

**Figure 4:**
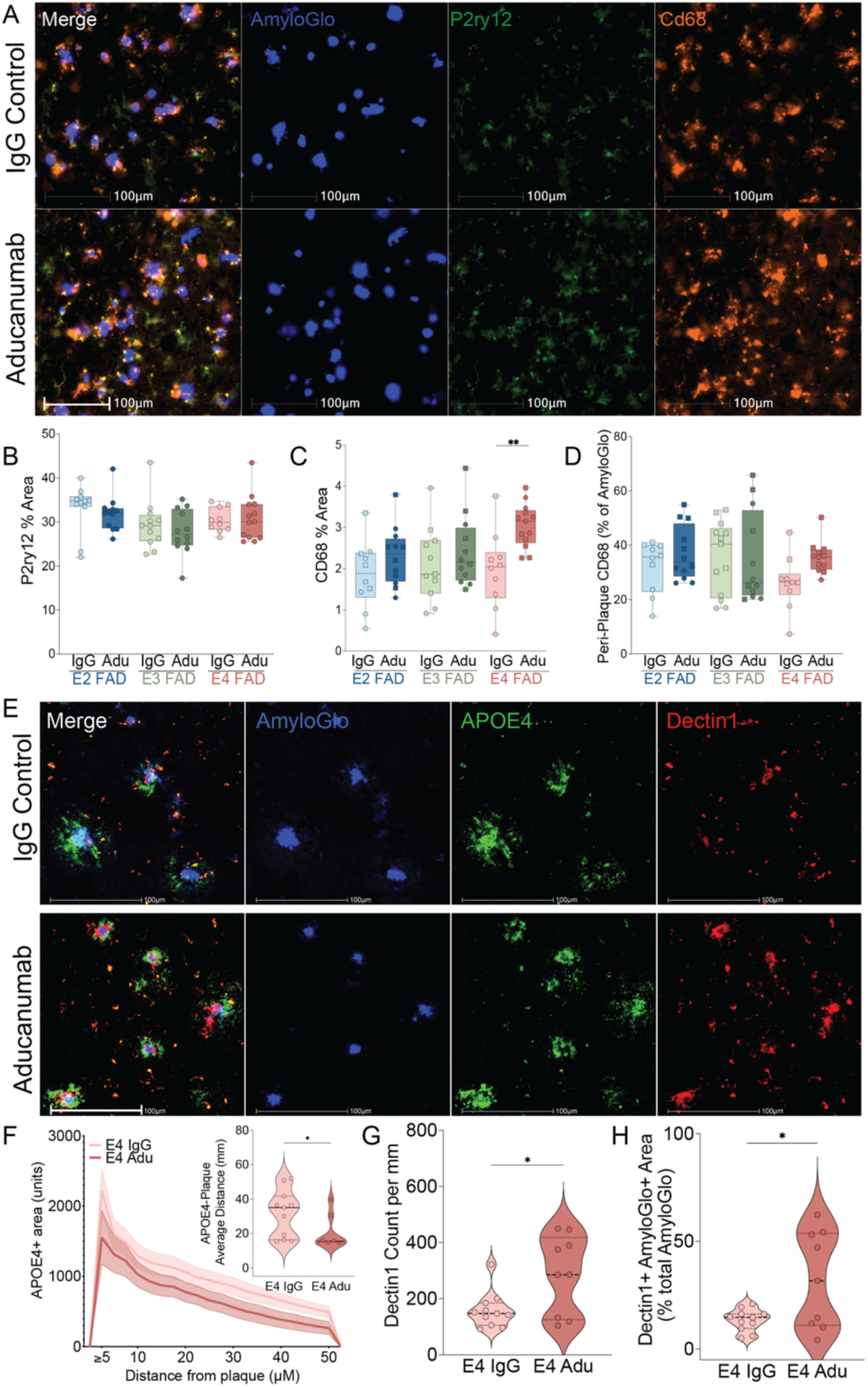
Aducanumab drives microglial shifts towards proinflammatory phenotypes in E4FAD mice. (A) Representative images of microglial staining with P2ry12 for homeostatic microglia (green) and CD68 for activated microglia (orange) in E4FAD. (B) Quantification of P2ry12 positive stain area. (C) Quantification of CD68 positive area. There is an increase in CD68 area after chAdu treatment in E4FAD mice. (D) Colocalization of CD68 and AmyloGlo. (E) Representative image of AmyloGlo (blue), ApoE4 (green), and Dectin1 (red). (F) Proximity histogram of APOE4 to plaque with inset analysis of average distance between E4 and plaque. (G-H) Quantification of total Dectin1 count and colocalization of Dectin1 and plaque (as % total AmyloGlo). (B-D) 2-way ANOVA with Tukey’s post-hoc comparisons. (F-H) 2-tailed T-test with Welch’s correction (p-value * <.05, ** <.005 *** <.001), specific n’s and p-values for each analysis are included in supplementary data file 3.

To corroborate this, we stained for two noted disease associated microglial (DAM) markers in E4FAD mice, APOE4 protein itself and Dectin1, the protein associated with DAM gene *Clec7a* (Fig 4E). While there was no change in the overall burden of APOE4, there was a reduction in the average distance of APOE4 protein to plaques after chAdu treatment (Fig 4 F). After performing plaque colocalization quantification, as well as plaque proximity assays, chAdu treated E4FAD mice, showed an overall increase in Dectin1 expression, but did not show significant increases in peri-plaque Dectin1 (Fig 4G-H). This may suggest that E4 has a more diffuse cellular response to chAdu treatment, particularly with activated and disease-associated microglia.

### Aducanumab alters vascular structure and perivascular gliosis in E4FAD mice

As described earlier, total vascular area did not differ significantly across genotypes or treatment groups (Fig 1F). However, chAdu treatment increased vascular branching area and total branch length in E2FAD and E3FAD mice but not E4FAD mice, normalized to tissue area (Fig 5A-C). This may suggest that E2 and E3 FAD mice have greater vascular plasticity than E4FAD mice, allowing for more effective vascular remodeling after injury and/or treatment..

**Figure 5.**
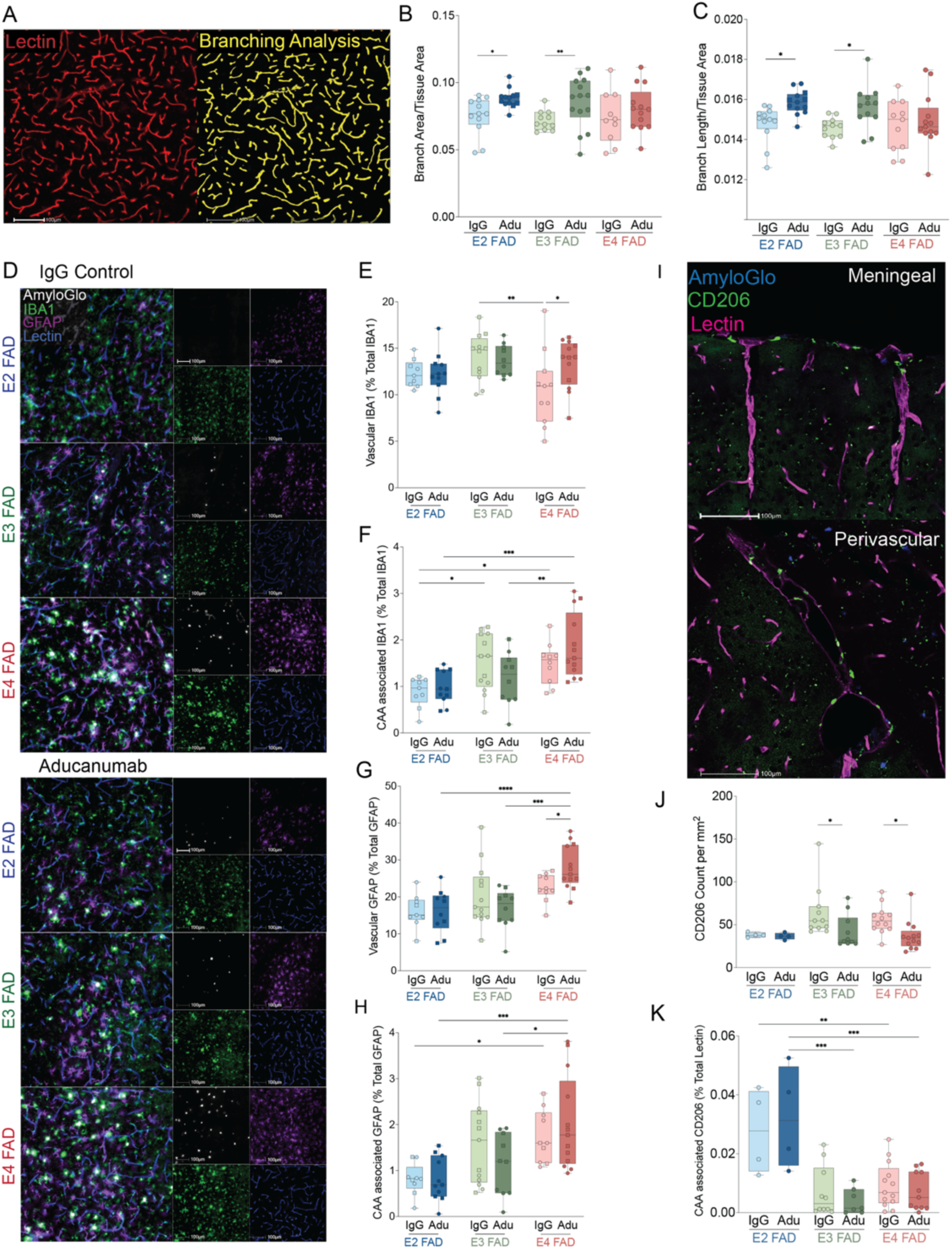
Aducanumab drives E4-specific increases in perivascular gliosis. (A) Vascular image analysis depicting isolectin vessels and analysis markup. (B) Total branching area normalized to tissue area. (C) Total branch length normalized to tissue area. (D) Representative images of perivascular gliosis withIBA1 shown in green, GFAP in purple, AmyloGlo in white, and isolectin in blue. (E) Quantification of IBA1 and isolectin colocalization. (F) Colocalization of IBA1 with CAA+ vessels. (G) Quantification of perivascular GFAP. (H) Colocalization of GFAP and CAA+ vessels. (I) Representative images of staining for border associated macrophages withAmyloGlo staining in blue, CD206 in green, and isolectin in pink. (J) Quantification of CD206 counts normalized to tissue area. (K) Colocalization of CD206 with CAA+ vessels. 2-way ANOVA with Tukey’s post-hoc comparisons (p-value * <.05, ** <.005 *** <.001), specific n’s and p-values for each analysis are in supplementary data file 3.

Perivascular microgliosis and astrogliosis, assessed by IBA1 and GFAP immunoreactivity surrounding isolectin-labeled vessels, were selectively increased in chAdu-treated E4FAD mice but not in E2FAD or E3FAD mice (Fig 5D-H). Neither IBA1 nor GFAP immunoreactivity around CAA deposits differed significantly across treatment groups in any genotype (Fig 5D-H). To assess border-associated macrophage (BAM) responses, we quantified CD206 immunoreactivity across genotypes and treatment groups. CD206 expression was decreased in E3FAD and E4FAD mice following chAdu treatment, with no significant differences observed around CAA deposits (Fig 5I-K). Together, these data demonstrate that chAdu treatment drives E4-specific perivascular glial activation and isoform-dependent differences in vascular remodeling, alongside BAM responses shared across E3 and E4 genotypes. Similar to the activated microglial findings described above, this increased gliosis may be plaque/CAA-independent, suggesting a more widespread inflammatory vascular phenotype in E4FAD mice.

### Aducanumab drives spatially localized and cell-specific responses in E4FAD mice

To characterize cellular and genomic drivers of E4-associated ARIA, we em ployed scRNAseq and Xenium spatial transcriptomics. Due primarily to the increased risk of AD in females, as well as the more robust and consistent pathology in female E4FADs, transcriptomics were performed on brain tissue from 6 female E4FAD mice (n=3 per treatment group)(*22, 37*). The middle third of each brain was hemisected, with one hemisphere processed for scRNAseq and the contralateral hemisphere sectioned at 10μm for Xenium (Fig 6A). While scRNAseq data did not reveal any changes in cellular composition after chAdu treatment, there were cell-specific changes in differential gene expression (Fig 6B-D). Xenium data corroborated these findings, as there were no significant differences between treatments or regions (Figure 6E-G, Supp Fig 5A-C). Again, while there were only modest gene expression changes, microglia and astrocytes showed regional alterations in gene expression (Fig 6H).

**Figure 6:**
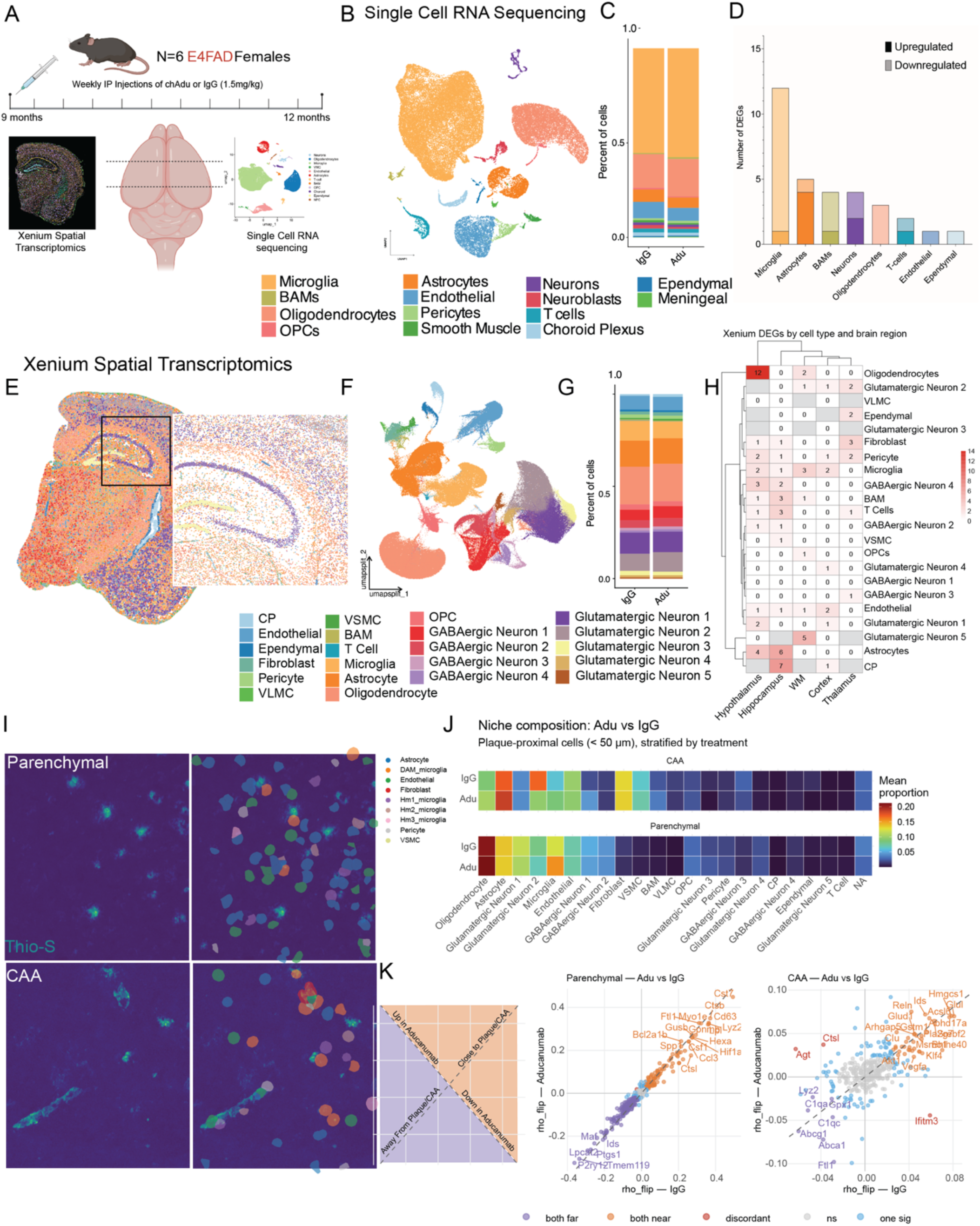
Aducanumab drives transcriptomic changes in immune and vascular cells in E4FADsFAD mice. (A) Parallel experimental design for single cell RNAseq (scRNAseq) and Xenium spatial transcriptomics (ST). (B) Single Cell RNAseq UMAP of detected cell types. (C) Cell proportions in scRNAseq dataset aggregated by treatment group. (D) Differential gene expression analysis of SCRNAseq data reveals DEGs in multiple cell types, primarily microglia. (E) Representative brain section of Xenium spatial transcriptomics. (F) UMAP of cells detected by Xenium ST. (G) Proportions of cell types by treatment group in Xenium ST. (H) Differential gene expression analysis of Xenium ST stratified by cell type and region. (I-J) Overlay of Xenium ST cell segmentation on Thioflavin S positive parenchymal plaques and CAA. (K) Differential Spearman analysis scatter plot comparing DEGs by treatment group and plaque proximity.

Thioflavin-S co-staining of Xenium sections enables spatial analysis of transcriptomic changes by plaque proximity (Fig 6I, Supp Fig 5D). chAdu treatment altered cellular composition in both parenchymal plaque and CAA niches, predominantly affecting astrocytes and microglia (Fig 6J, Supp. Fig 5E), with the CAA niche showing the most pronounced changes, including increased expression of *Reelin (Reln)* and *Clusterin (Clu)* in cells proximal to CAA deposits (Fig 6K). Together, these data identify the CAA niche and hippocampus as spatial loci of chAdu-driven transcriptomic response in E4FAD mice.

### Aducanumab alters the regional and plaque niche composition of microglia and astrocytes in E4FAD mice

We then independently subclustered microglia and astrocytes and annotated by functional state based on relative gene expression profiles in Xenium (Supp. Fig 6) and scRNAseq data(Supp. Fig. 7C-D). Microglial subclustering of single cell RNAseq data revealed 8 unique clusters (Fig 7A). Proportionality analysis of scRNAseq data revealed a significant decrease in Homeostatic 1 (Hm1) microglia and a trending increase in Transitional DAM microglia in chAdu-treated E4FAD mice (Fig 7B-C). Microglial state annotations were cross-validated against the Fumagalli et al. framework using module scoring, and overlaps between our scRNAseq-defined states and published microglial states were calculated (Fig 7D) (*38*). Xenium spatial analysis of microglia identified 10 unique subclusters (Fig 7E-G), with regional shifts in microglial state composition, with significant increases in interferon-responsive microglia (IRM) in the hippocampus and activated response microglia (ARM) in the cortex following chAdu treatment (Fig 7H). In addition to microglia, we subclustered astrocytes from our Xenium data, revealing 8 unique clusters that aligned with existing literature on astrocyte states in aging and disease (Fig 7I-K)(*39–42*). We found that there were also regional differences in astrocyte composition, with significant increases in interferon responsive reactive astrocytes (IRRAs) in the cortex, and decreases in lipid biosynthesis and metabolic reactive astrocytes in the thalamus (Fig 7L).

**Figure 7:**
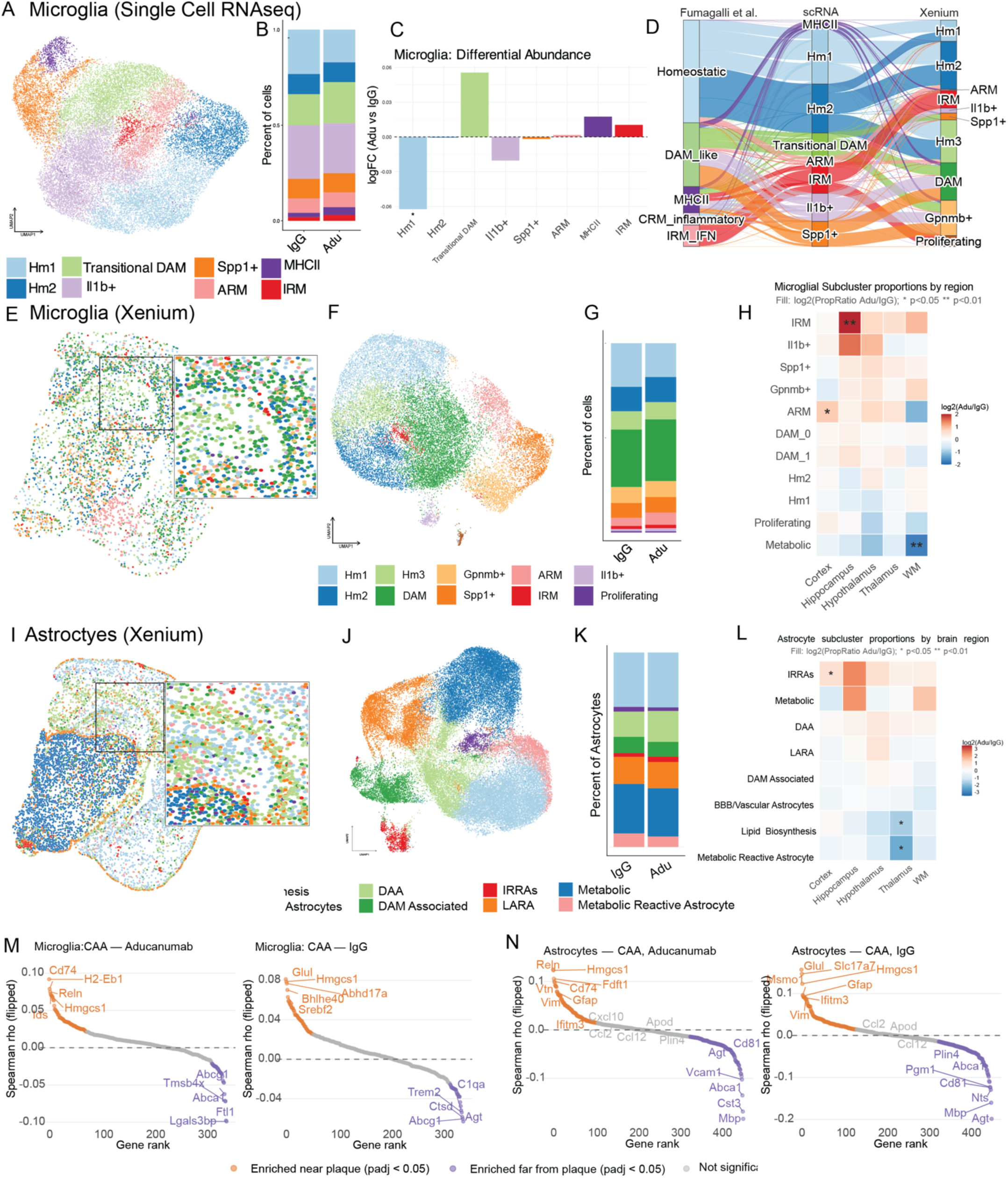
Microglia and Astrocytes show spatially distinct responses to Aducanumab treatment in E4FAD mice. (A) UMAP of SCRNAseq microglial subsets. (B) Microglial cell proportions in SCRNAseq dataset aggregated by treatment group. (C) Differential abundance of microglial subtypes seen in SCRNAseq. (D) Sankey plot showing microglial states defined in Fumagalli et al. 2025, SCRNAseq microglia, and Xenium Microglia. (E) Representative brain section of Xenium ST microglia. (F) UMAP of microglial subsets detected by Xenium ST. (G) Proportions of microglial subsets by treatment group in Xenium ST. (H) Proportionality analysis of microglial subtypes by brain region. (I) Representative brain section of Xenium ST astrocytes with zoom inset. (J) Astrocytes UMAP from Xenium ST. (K) Astrocyte subcluster proportions by brain region in Xenium ST. (L) Differential abundance of astrocyte subclusters by brain region. (M-N) Spearman rank plot of genes altered near CAA in aducanumab and IgG treated xenium brains in microglia and astrocytes respectively.

Within the CAA niche, chAdu-treated mice showed increased microglial expression of the AD-associated gene *Reelin* and MHCII genes *Cd74* and *H2-Eb1,* relative to IgG controls (Fig 7M). Astrocytes demonstrated similar CAA-proximal changes, with upregulations of *Cd74* and *Reln*, along with *Vitronectin (Vtn)*, suggesting greater proximity to the neurovascular unit and increased capacity for vascular remodeling in chAdu treated astrocytes (Fig 7N).

### Aducanumab drives neurovascular domain and network changes in E4FAD mice

Spatially distinct transcriptomic domains were identified across cortical and hippocampal regions using BANKSY, a clustering algorithm reflecting both transcriptomic identity and local tissue context (*43*) (Fig 8A). Differential neighborhood enrichment analysis via Squidpy, stratified by proximity to parenchymal plaques or CAA, identified increased co-enrichment of vascular-related pathways in chAdu-treated mice centered on BAM and astrocyte nodes (Fig 8B). Spatial domains enriched for vascular cell types (domains 5 and 6) and T cells (domain 18) were identified and subjected to neighborhood analysis (Fig 8C). Delta Z-score analysis between chAdu and IgG groups revealed increased spatial co-localization of ARM microglia, BAMs, and astrocytes with endothelial cells and vascular mural cells in chAdu-treated animals, alongside decreased fibroblast and VLMC neighborhood enrichment in perivascular domains (Fig 8D-E).

**Figure 8:**
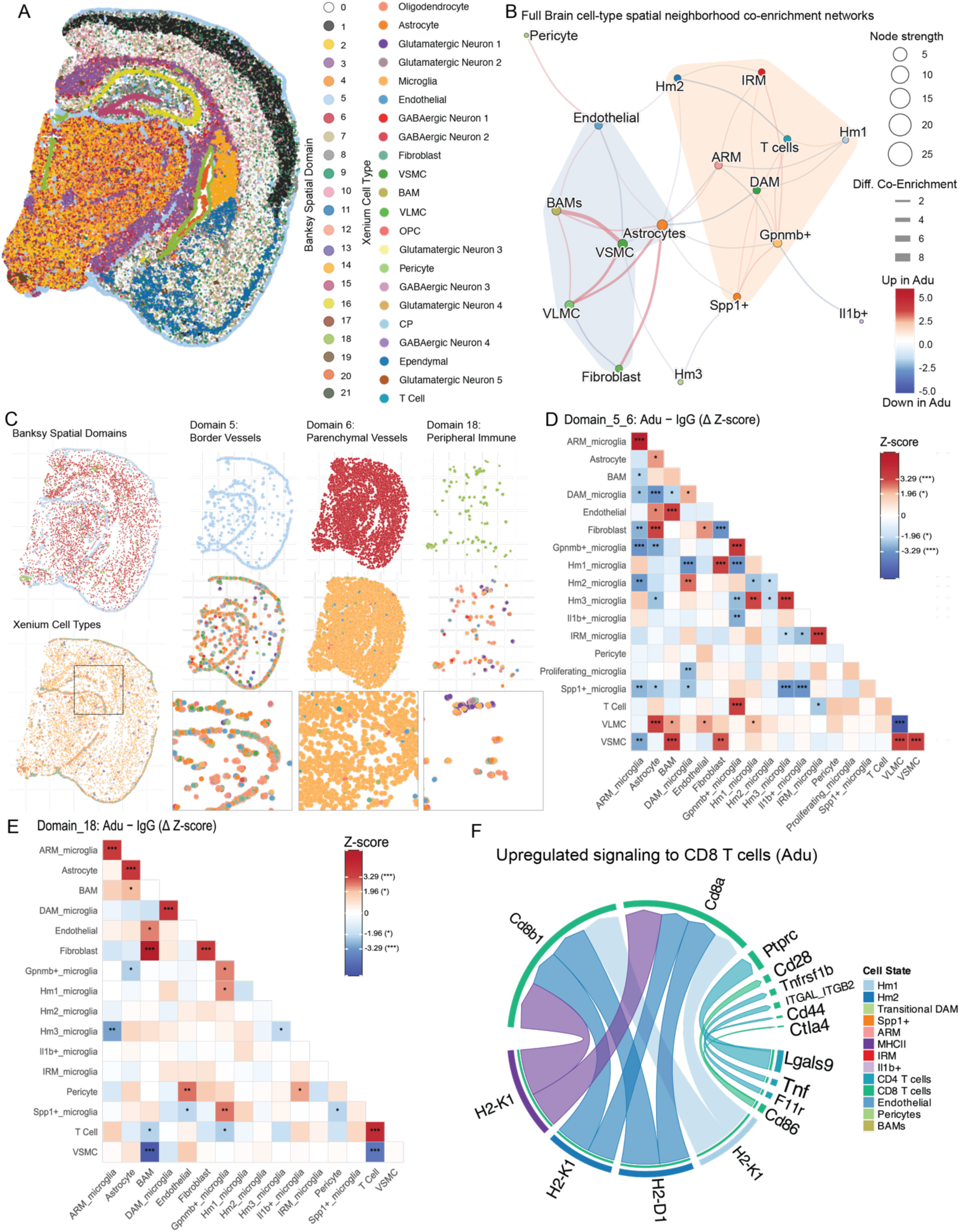
Aducanumab drives spatial network changes in E4FAD mice. (A) Spatial domains generated using the BANKSY package in R. (B) Spatial network co-enrichment plot with pseudo-neighborhoods generated by ggforce. (C) Image plot representation of selected domains 5, 6, and 18 corresponding to vascular and immune compartments with cell type compositions. On left are the selected domains colored by domain identity (top) and cell type (bottom). On right are each of the domains represented individually with domain identity (top), cell type composition (middle), and zoomed cell type composition (bottom). (D-E) Heatmaps of spatial co-enrichment z-scores for selected domains described in C, stratified by treatment. (F) CD8 signaling pathways upregulated in Aducanumab treated mice.

Differential CellChat analysis of immune-vascular cell communications in scRNAseq data identified increased BAM-to-CD4 T cell MHCII signaling and decreased BAM-to-CD8 T cell MHCI signaling in chAdu-treated mice (Supp. Fig 4B-C). Multiple microglial subclusters and vascular cells showed increased ligand-receptor signaling directed toward CD8 T cells in chAdu-treated relative to IgG-treated animals (Fig 8F). Together, these data demonstrate that chAdu treatment reorganizes the perivascular immune niche in E4FAD mice, with convergent shifts in spatial co-enrichment and intercellular signaling centered on BAM, astrocyte, and microglial interactions with vascular cell types.

## DISCUSSION

Although anti-amyloid antibodies such as aducanumab, lecanemab, and donanemab, have now been approved for treatment of AD in some populations, fears over ARIA risk, coupled with the high cost and burden of safety monitoring of these drugs, have significantly tempered uptake of these novel therapies (*44*). ARIA describes findings of fluid effusion (ARIA-E) or microhemorrhage (ARIA-H) that occur at significantly higher rates – with worse symptoms and potential for fatal reactions – in carriers of APOE4 (*2, 3, 10, 11, 45, 46*). E4 carriers comprise nearly 70% of the AD population, making it critical to understand the relative contributions of E4 to ARIA and related complications.

Several studies have examined brain and peripheral changes following anti-amyloid antibody treatment, though preclinical work has focused primarily on ARIA mechanisms independent of *APOE*. Here, we present a multi-modal investigation into the role of *APOE* genotype in anti-Aβ antibody therapy using the humanized EFAD mouse model. We demonstrate that even subclinical doses of a chimeric aducanumab drives increased microhemorrhages in E4FAD mice relative to E2/E3FAD and IgG controls, confirmed by both MRI and Prussian Blue histology, findings that remarkably mimic human clinical observations. E4FAD mice also exhibited elevated overall, peri-plaque, and perivascular microgliosis, increased activated and DAM signatures, and heightened overall and perivascular astrocyte reactivity following treatment. Additional findings include reduced CD206 in E3/E4FAD mice and vascular structural alterations in E2/E3FAD but not E4FAD mice. Transcriptomic analysis further revealed E4FAD-specific alterations in plaque and CAA niche cellular composition, gene expression, spatial reorganization, and differential vascular and immune cellular communication. Together, these results establish a preclinical model for studying APOE’s role in anti-amyloid therapy responsiveness and identify E4-mediated perivascular and glial alterations that may underlie ARIA-like events.

We believe this work helps bridge a key gap in understanding ARIA pathogenesis: the role of a strong genetic risk factor for both AD and ARIA – APOE4. Studying anti-amyloid therapy in preclinical models expressing murine *Apoe* has several limitations, primarily that murine ApoE exists in only one isoform and only contains ∼70% homology with human APOE, making assessments of the the protein’s precise role in ARIA difficult (*21*). Furthermore, murine ApoE has been shown to produce significantly accelerated onset of pathology with variable deposition patterns favoring parenchymal plaques over CAA compared to APOE4, highlighting the need for study in human-relevant models of *APOE* and amyloidosis (*21–23, 47*). To the best of our knowledge, this is one of a limited number of existing studies to examine anti-amyloid treatment across human *APOE* genotypes(*25–27, 48*). Amyloid expressing models do not capture the full scope of AD pathology, and the 5xFAD model in particular exhibits different patterns of plaque deposition than human AD cases. Despite this limitation, however, the use of these models expressing human APOE has allowed us to answer several important questions about the interactions between anti-Aβ therapies, APOE4, and the immune system. Additionally, our findings highlight the potential maladaptive responses that E4 drives to anti-amyloid therapy and the key need to understand this underlying pathology. And while the dose used in this study is ∼85% lower than the standard clinical dose, other work has identified similar changes with subclinical dosing strategies (*28, 29*). This may have implications for current dose titration strategies; while they have shown reductions in detected ARIA rates, the underlying gliosis and vascular dysfunction may still be ongoing (*49*).

In this study, we have demonstrated that ARIA-H like events occu r in an *APOE* genotype-dependent manner. Prior studies have established that microbleeds increase in E4FAD mice after treatment with chAdu, albeit at a significantly supratherapeutic dose (40-50 mg/kg), as well as increases in microbleeds in both E2 and E4FAD mice when treated with an alternative murine anti-Aβ antibody (*25, 26, 28*). More recently published work examining the role of E3 vs E4 in anti-amyloid therapies has demonstrated very similar results with an acute dosage paradigm (*48*). Our data reproduces these findings at a subtherapeutic dose (1.56 mg/kg), as we also see an increase in microbleed occurrence in E2 and E4FAD mice after chAdu treatment measured by Prussian Blue positive area. Interestingly, we also reproduced a similar trend when analyzing MRI scans for hypointense regions, noting that there were significantly increased bleeds in E4FAD mice. Both findings mimic descriptions of clinical ARIA risk, with E4 homozygotes having significantly higher occurrences of microbleeds, at baseline and after treatment, providing additional translational validation for our model system (*2–4*). Furthermore, these findings indicate that, while E2 may be neuroprotective with regard to AD risk, there may still be significant potential for vascular injury and hemorrhage. In fact, there is some, albeit limited, evidence that suggests E2 carriers have a higher risk for CAA (*50*). While the literature supporting elevated rates of CAA is limited due to low numbers of E2 homozygotes with severe amyloid pathology, there is growing evidence that E2 induces a CAA-related vasculopathy, in which there is greater potential for vascular fragility in CAA-laden vessels, as demonstrated by a 2019 meta-analysis of the link between CAA, superficial siderosis, and E2 status (*50–54*). This E2 mediated vasculopathy may explain findings in our study – and a similar finding in Pankeiwicz et al. – of increased microhemorrhages in E2 mice (*25*). Furthermore, a recent neuropathological study in human brain tissue identified a strong link between existing pathology and CAA in E2 carriers, particularly when there is no clinical diagnosis of AD (*55*). In mouse models engineered to overproduce amyloid pathology, this could explain the greater vascular fragility and microhemorrhage occurrence that we and others have seen. Altogether, this raises significant questions about the role of E2 in CAA and vascular dysfunction, and the risk profile of E2s being treated with anti-amyloid therapies. While the population of individuals who carry an allele of E2 that require treatment with this drug is likely extremely small, there is nevertheless a small subset of individuals who carry both an E2 and an E4 allele concurrently (*6*). In these individuals, whose AD risk is closer to that of an E3/E4 than an E2/E3, this potential “double” risk of severe ARIA-related sequelae may be an important additional factor to consider when initiating anti-amyloid therapies. This also raises the question of heterozygote risk profile, a limitation of our current study which only utilizes homozygous EFAD mice, and was not powered to detect sex dependent differences in ARIA risk. E3/E4 individuals outnumber E4 homozygotes almost 2:1, and carry very different ARIA risk profiles, with risk almost half that of E4 homozygotes (*2–4, 6*). It is unclear what the underlying E4 gene-dose mechanism of ARIA is, and understanding this may provide potential therapeutic targets to limit risk in all E4 carriers.

Accompanying these changes in microbleed occurrence and plaque clearance are alterations in microglial and astrocyte reactivity after treatment. Here, we show that there are overall increases in IBA1, CD68, and Dectin1 in E4FAD mice, findings seen by several other papers at both supra and sub therapeutic dosages in both E4FAD and APPe4 mice (*25, 26*). In fact, in one published report of fatal ARIA-related reactions in an E4 homozygote, there was a notable elevation of pro-inflammatory microglial signaling, particularly with increased CD68 in the perivascular spaces (*9, 56*). Increases in microglial activation have also been shown in models expressing murine *Apoe* at both acute and chronic timepoints, with even two weekly doses of 40mg/kg chAdu being sufficient to induce widespread microglial activation (*36*). When examining the transcriptomes of mice treated with chAdu or other similar anti-amyloid antibodies, the resulting programs are generally overwhelmingly immune focused, a finding that is duplicated in our study as well as others, and generally involve antigen-presenting or polarizing functions (*14, 19, 20, 25, 26, 28, 57, 58*). Further transcriptomic characterization of microglia from both murine *Apoe*-expressing and E4FAD mice indicates an expansion of disease associated, activated microglia, sometimes to the point of immune exhaustion. This near exhausted state was recently defined as Terminally Inflammatory Microglial (TIM) and was seen in aged and chAdu treated E4FAD mice at significantly higher rates than controls (*27*). This signature may help explain immune blunting that is seen after a long washout of chAdu, as the inflammatory responses seen post treatment have faded or been silenced, though this may also be a product of using a non-chimerized antibody in a murine model (*36*). Interestingly, a, dose-dependent microglial exhaustion was seen in early intervention with chAdu in APP/SAA mice, where a chronic 10 mg/kg dose appeared to drive lowered microglial activation than a 1 mg/kg dose (*28*). This raises the question of whether intervention earlier in life may prevent ARIA through reduced inflammatory signaling, and whether this may protect against E4 driven ARIA.

Our laboratory has previously demonstrated that E4FAD microglia exhibit a heightened baseline inflammatory state that predisposes them to chronic neuroinflammation, while others have shown an apparent inability of APOE4 microglia to mount a robust DAM response to late-life amyloid pathology (*59–61*). While seemingly contradictory, these findings likely reflect two temporally distinct manifestations of the same underlying process rather than mutually exclusive phenomena. Our previous work captures a cell-intrinsic inflammatory baseline driven by APOE4, characterized by increased HIF1α expression, a disrupted TCA cycle, and an inherently pro-glycolytic state consistent with constitutive immune activation (*61*). This chronic inflammation driven by E4 may persist across the lifespan, ultimately driving microglia toward a state of functional exhaustion that prevents adequate responses to late-life AD pathology. Another recent study demonstrated that anti-amyloid antibody therapy mediates amyloid clearance by activating microglial effector functions in an Fc-dependent manner, inducing a focused transcriptional program that enhances phagocytosis, lysosomal degradation, metabolic reprogramming, and antigen presentation – a mechanism that is absent when the Fc fragment is silenced (*58*). Together, this data may suggest that, to drive plaque phagocytosis in response to chAdu treatment, there may be an FcR mediated immunometabolic reprogramming in microglia, though our data may suggest that E4 microglia may be driving additional maladaptive responses to this reprogramming, particularly in certain inflammatory niches, which may then result in ARIA-like events.

One such inflammatory niche is the peri-plaque region. It is well known that APOE, and particularly APOE4, is one of the key co-aggregating proteins along with Aβ in amyloid plaques (*62–65*). In fact, recent work has further identified the molecular basis of E4 mediated aggregation, suggesting that the initial interaction of APOE and Aβ may occur in the phagolysosomes of microglia, and that increased co-aggregation in E4 microglia may explain the more aggressive plaque pathology seen in E4 carriers (*66*). Other studies have also suggested that APOE is a key facilitator of microglial phagocytosis of amyloid plaques, and that specific APOE isoform differences may appear at the level of clearance of plaques, rather than at the level of aggregation (*67, 68*). Putting these findings into the context of our data, where increases in peri-plaque E4, along with increased microglial IBA1, Dectin1, and CD68, may point to a role of chAdu in pushing increased plaque phagocytosis by E4 microglia. Whether or not this is beneficial or effective is unclear, but studies described above would suggest that this is likely a maladaptive effect of treatment. The combination of elevated baseline inflammatory state, immunological exhaustion, and phagolysosomal overloading may produce a toxic combination that is driving brain-wide inflammation that centers on the perivascular spaces.

Histological and spatial transcriptomic analysis of the CAA and vascular niches after chAdu treatment seem to corroborate this premise. Studies examining the neurovascular coupling in mice after anti-amyloid therapy have demonstrated that there is an alteration in vascular integrity, likely driven by clearance of vascular amyloid deposits or CAA (*19, 20*). Furthermore, there is a noted detrimental effect of E4 on neurovascular coupling even in the absence of AD pathology (*69, 70*), suggesting a potential amyloid-independent role of E4 on the cerebrovasculature, an effect which may be further exacerbated by CAA deposits. Our data appears to support this, as we observed restorations of vascular complexity after treatment in E2 and E3FAD mice, but not in E4FAD mice. This post-treatment increase in vascular branching could reflect greater vascular plasticity in E2 and E3FAD mice compared to E4FAD mice. This lack of neovascularization in E4FADs may also be explained by neurovascular uncoupling seen in both E4 knock in and 5xFAD mice (*71, 72*). Activation of HIF1a, as described earlier in E4FADs, mixed with pre-existing vascular-metabolic uncoupling may exacerbate the vascular injury caused in E4FADs, producing more ARIA-like events (*73, 74*). The clearance of CAA deposits from the brain during treatment with antibodies such as chAdu could further expose underlying vascular and smooth muscle damage that E4 carriers struggle to respond to due to immune and metabolic dysfunction, thereby leading to vascular injury, inflammation, and findings of ARIA – regardless of the overall burden of CAA.

Of course, it is difficult – if not impossible – to disentangle the interplay of E4 and CAA in a pathology dependent context, as E4 inherently leads to increased plaque deposition in both the parenchyma and vasculature(*8*). In fact, early literature during the inception of amyloid targeting as a promising therapeutic strategy seemed to suggest that the administration of passive anti-Aβ antibodies may inherently increase CAA through vascular clearance mechanisms (*14*). It is also important to consider that in the clinical context (and in many preclinical studies including our own), these antibodies are injected or infused into the periphery, rather than directly into the CNS. In the current study, antibodies were injected to the intraperitoneal space; therefore, similar to intravenous injections in the clinic, the first Aβ that these drugs interact with is presumably CAA – and immune cells in the neurovascular unit are potentially the first cells to have an FcR mediated response to the antibodies.

This logic has fueled increased study into border-associated macrophages, especially in the contexts of neurovascular dysfunction and ARIA. BAMs have been shown to influence vascular integrity and drive BBB leakage through ROS production, and more specifically E4 derived from BAMs appears to be highly responsible for this vascular dysfunction (*69, 70*). Our work suggests that treatment with chAdu, adds to the pre-existing E4-dependent vascular dysfunction by increasing vascular inflammation and engagement. This is likely occurring through increased recruitment of BAMs and microglia to the perivascular spaces, and increased reactivity of these cells and other neurovascular cells (astrocytes, pericytes, endothelium, etc.) – potentially independent of CAA. In fact, recent papers from Taylor et al. show a similar increase in BAM activation at the perivascular spaces which may be driving these phenomena, though these studies utilized models of murine *Apoe* with a preference for CAA deposition (*19, 20*). The downregulation of CD206 that we note after treatment may be indicative of a similar shift in our E3 and E4 FAD mice, though the occurrence of increased perivascular gliosis is only seen in E4FAD mice. Another recently published study supports this, noting in both murine *Apoe* and human *APOE4* expressing mice, that there appears to be vascular fragility either induced or exposed by anti-amyloid therapy – in this case suggesting that complement signaling is the driving factor(*75*). While we do not see complement pathway signaling as top hits in our E4FAD transcriptomic data, we do find similar increases in glial reactivity in the perivascular niche, suggesting that complement signaling may be part of a larger immune-vascular inflammatory “burst” that occurs during treatment with these anti-amyloid antibodies (*75*).

This increase in perivascular signaling may also be setting up an additional factor to consider – the adaptive immune response to chronic antibody therapy. Several recent studies have demonstrated that in diseased states, microglia (and even astrocytes) produce chemotactic signals to drive homing of antigen specific T-lymphocytes into the CNS (*40, 76–80*). In fact, a recent collaborative study between our group and a regional medical center demonstrated peripheral expansions of T-effector memory (TEM) and T-effector memory expressing CD45RA (TEMRA) subsets in patients who developed ARIA (*81*). These subsets have been shown in several AD related studies to drive proinflammatory milieu in the brain, albeit primarily in response to Tau pathology rather than amyloid (*79, 80*). Our data suggests that, even in the absence of tau pathology, chAdu treatment drives increased MHCI-CD8 interactions, along with increases in MHCII-high antigen presenting microglia and astrocytes, particularly Cxcl10 high astrocytes. These factors together support a role of T-lymphocytes in driving or responding to ARIA. The temporal dynamics of this response, however, remain to be resolved. As a response to these concerns surrounding the neurovascular interface and elevated immune signaling, there has been a push towards therapeutics that bypass this BBB interaction, for example through the addition of a transferrin receptor binding fragment (*82*). These drugs appear to have significantly greater brain penetrance, and avoid producing ARIA through vascular binding, although their clinical efficacy has yet to be fully established(*83*).

In this study, we provide an in-depth histological and transcriptomic characterization of the E4-driven response to anti-amyloid immunotherapies, offering novel insight into mechanistic alterations that may drive ARIA in E4 carriers. Our findings converge on the neurovascular unit as the center of E4-mediated immune and glial dysfunction, positioning immune-neurovascular interactions as tractable targets to reduce ARIA risk in the highest-need population.

## MATERIALS AND METHODS

### Animals and Study Design

EFAD mice expressing 5 mutations found in familial AD and crossed to express human alleles of *APOE* (E2 FAD, E3 FAD, and E4 FAD)(*22, 23*). E2FAD, E3FAD, and E4FAD male and female mice were matched into cohorts at 9 months of age and randomized to receive one of two treatments. Sexes were balanced as best as possible to have even distribution across genotypes and treatment groups, however, based on final numbers the study was not powered to analyze sex effects. An n=5 minimum per sex, genotype, and treatment were selected to achieve a total study sample size of n = 72 across all groups. Additionally, a separate group of n = 6 E4FAD mice were age matched to be used for transcriptomics analyses described below, with 3 in each treatment group. Baseline and endpoint plasma was collected from all animals via retro-orbital bleed. Investigators were not blinded to animal ID during treatment to allow for appropriate health monitoring, but were blinded during experimental testing and analysis.

### Chimeric Aducanumab Antibody Synthesis and Treatment

Chimeric aducanumab (chAdu) and IgG isotype control were synthesized by LakePharma (Michigan, USA, now Curia Biologics) and purified as previously described (*84*). Antibody was shipped to UKY under a material transfer agreement and was stored at −80°C until use after being aliquoted into necessary volumes for weekly cohort injections based on starting weights of animals. Aliquots were thawed on ice and diluted using sterile 0.9% isotonic saline for injections as described below.

Mice were randomly selected to receive one of two treatments: chimeric aducanumab (chAdu) or IgG isotype control. Mice received weekly intraperitoneal injections of the selected treatment. chAdu and IgG were diluted to a subclinical dose of 1.56 mg/kg in 0.1 mL of sterile isotonic saline prior to injection. Body weight was tracked throughout the injection course. Treatment was well tolerated, with minimal loss of animals throughout the study.

### MRI Imaging and Image Analysis

All MRI data were acquired on a Bruker BioSpin AV4 Neo Clinscan 74/30 (7.06 T) scanner at the University of Kentucky using ParaVision 360.3.2. A volume resonator (RF RES 300 1H) and a 4-element phased-array surface coil (RF ARR 300 1H) were used for radiofrequency transmission and signal reception, respectively. T2-weighted structural images were acquired using a TurboRARE sequence with the following parameters: field of view = 20 × 20 mm², matrix = 256 × 256, slice thickness = 0.8 mm, 13 slices, repetition time = 2,800 ms, effective echo time = 33 ms, echo train length = 8, and 2 averages. Total acquisition time was approximately 3 minutes. Susceptibility-weighted images were acquired using a 2D gradient-echo FLASH sequence (15te_SWI_FLASH) with the following parameters: field of view = 20 × 20 mm², matrix = 290 × 290, slice thickness = 0.6 mm, 17 slices, repetition time = 564 ms, echo time = 15 ms, flip angle = 30°, and 3 averages. Total acquisition time was approximately 8 minutes. All images were obtained the day prior to final injection and tissue collection. After scanning, mice were allowed to recover and acclimate for 12 hours prior to sacrifice.

Images were imported into MIM Encore (Version 7.3.2) analysis software (MIM Software Inc., Cleveland, OH; 2024) and manually registered to the Allen Common Coordinate Framework (ACCF) as has been previously described (*85, 86*). The registered image was manually examined by two independent viewers to provide count and Allen region of hypointensities. Regional quantifications were imported into proprietary AtlasModeler software for heatmap generation. Counts were then collapsed to more broad regional quantification (Hippocampus, Thalamus, Cortex) for count analysis.

### Isolectin Vascular Labeling

Prior to sacrifice, mice were placed under isoflurane anesthesia for 3 minutes, before receiving a retro-orbital injection of fluorescent isolectin (Vector Labs DL-1178-1). Isolectin was allowed to circulate for 15 minutes before sacrifice.

### Brain Tissue processing

At the completion of the study, animals were euthanized with a 0.1mL injection of sodium pentobarbital. Blood was collected through retro-orbital puncture for plasma isolation. Mice were then transcardially perfused with 0.9% saline for 2 minutes. Brains were collected, with the right hemibrain drop fixed in 4% paraformaldehyde for 24 hours with subsequent transfer to sucrose solution. The left hemibrain was flash frozen in liquid nitrogen for long term storage. Fixed tissue was transferred to 30% Sucrose after 24 hours, and then cryosectioned into 30 μm sections for histological measurements.

### Single Cell RNA Sequencing and Xenium Prep

Animals in the transcriptomics cohort (n=6) were treated as described above. At the conclusion of 12 weeks, animals were sacrificed without isolectin injection or MRI scan to avoid fluorescent overlap or isoflurane effects respectively. Mice were perfused with 1X Dulbecco’s Phosphate Buffered Saline (DPBS) for 2 minutes before brain dissection. The right hemibrain was dissected for single cell sequencing and processed with the Miltenyi Adult Brain Dissociation Kit (Miltenyi Biotec #130-107-677). Enzyme mix was thawed and prepared the morning of, and placed in C-tubes. Brains were cut into small chunks and placed in the corresponding tube for gentleMACs dissociation. After dissociation, debris was removed and the resultant single cell suspension was run on 10x Genomics Chromium X 3’ v4. cDNA libraries were sent to Novogene for sequencing on Illumina NovaSeq X-plus. The left hemibrain was taken for Xenium spatial transcriptomics. Xenium hemibrains were sectioned into 10 μm sections, and 6 hemibrains were mounted onto one Xenium slide cassette. Slides were incubated with fluorophore tagged oligonucleotide sequences using a custom 480 gene panel. Slides were run in the Xenium analyzer over two days. After running on Xenium, slides were stained with DAPI nuclear stain and Thioflavin-S for co-registration of transcriptomic data to amyloid pathology.

### Immunohistochemistry

30 µm serial coronal sections were selected from specific regions of interest (hippocampus, thalamus, and cortex) for all fluorescent immunohistochemistry, with three averaged replicates for each animal per stain. All staining was done free-floating unless otherwise stated. Sections were washed three times for 10 minutes each in 1X Phosphate Buffered Saline, incubated in 1X AmyloGlo solution diluted in slightly acidic saline (pH 6.7-6.8) and washed once in slightly acidic saline, followed by incubation in blocking buffer followed by primary antibody cocktail overnight (Table 1). Sections were then washed and incubated with secondary antibody cocktail for two hours at RT before another series of PBS washes and tissue mounting onto slides. Prolong Diamond Antifade mounting media (Thermo Scientific P36970) was used before coverslipping, and slides were allowed to dry overnight before sealing with clear nail polish.

**Table 1:** Primary antibodies and stains used in immunohistochemistry.

| Antibody | Host | Dilution | Manufacturer | Catalog # |
| --- | --- | --- | --- | --- |
| IBA1 | Rabbit | 1:500 | Abcam | Ab178846 |
| GFAP | Rat | 1:500 | Thermo-Fisher | 13-0300 |
| CD206 | Rat | 1:400 | Bio-Rad | MCA2235 |
| mDectin/Clec7a | Rat | 1:200 | InvivoGen | Mabg-mdect |
| ApoE4 | Rabbit | 1:500 | Cell Signaling Technologies | 39327S |
| CD68 | Rat | 1:250 | Bio-Rad | MCA1957 |
| LAMP1 | Rat | 1:400 | DSHB | 1D4B-s |
| Isolectin | N/A | 100 uL | Vector Labs | DL-1178-1 |
| AmyloGlo | N/A | 1x | Biosensis | TR-300-AG |
| Thioflavin-S | N/A | 0.5mM | Thermo Fisher | T1892 |

### Prussian Blue

Six evenly spaced sections (approximately 1.44 mm apart) were selected for Prussian Blue staining to verify microbleed occurrence in mice. Sections were mounted on slides and stained using glass coplin jars. The staining sequence was as follows: Rinsing of slides in MilliQ filtered H2O, 30 minutes of staining in Perl’s Prussian Blue solution (Abcam Iron Stain Kit ab150674), 3 H2O washes, 5 minute incubation in nuclear fast red solution, 4 sequential washes in H2O, differentiation in 70% EtOH, 3 washes in H2O, followed by a graded ethanol dehydration. Slides were coverslipped with Cytoseal60 (Fisher Scientific 23-244256) and allowed to dry overnight before imaging.

### Amyloid Beta ELISA

Aβ40/42 was quantified using the Meso Scale Discovery V-PLEX Aβ peptide panel 1 (6E10) kit (K15200E). Plasma samples were diluted 1:1 and the soluble and insoluble brain fractions were diluted 1:300 and 1:5000, respectively. Samples were normalized to total protein using a BCA protein assay kit (Thermo Scientific 23225). Brain amyloid data were analyzed by fitting to a standard curve (4PL), normalized to total protein in the sample, and analyzed as described below for between group variances. Plasma concentrations were also fitted to a standard curve and quantified by concentration of amyloid in solution. Plasma data was analyzed using a linear mixed model to account for effects of time, treatment, and APOE genotype.

### scRNAseq data processing

Single cell data analysis was conducted in R (v4.5.0) using Seurat (v5.3.0) and supporting packages (*87–94*). Cells with >30% mitochondrial RNA were removed. Doublet and low quality cell processing was done using median absolute deviation per sample on nFeatureRNA counts and scDblFinder (v1.24.0) (*95*). Normalization and scaling were done with SCTransform using 2000 features, with subsequent reciprocal PCA integration (rpca). Clustering was performed at a resolution of 0.8 for all cells and a resolution of 0.3 for microglial subset. UMAP dimensionality reduction was computed using 20 dimensions. Clusters were annotated based on canonical marker genes for astrocytes (*Aldoc, Aqp4, Gja1*), microglia (*P2ry12, Tmem119, Aif1*), oligodendrocytes (*Mog, Opalin*), OPCs (*Pdgfra, Opcml*), choroid plexus (*Ttr, Igfbp2*), ependymal cells (*Ccdc153, Dnah11*), vascular mural cells (*Bgn, Vtn*), endothelial cells (*Cldn5, Cdh5, Flt1*), neuroprogenitor cells (*Dcx*), neurons (*Snap25, Rbfox3*), T cells (*Cd3d, Cd3e*), and border-associated macrophages (*Mrc1, Pf4*). Any clusters expressing multiple cell-specific signatures were excluded from analysis. Microglia were subsetted and classified based on functional state and published gene signatures (*38, 59, 60*). Cell-cell signaling was evaluated using CellChat (v1.6.1) with the CellChatDB.mouse database (*93, 94*). To capture APOE-mediated interactions, we manually incorporated ligand *Apoe* and its receptors (*CHRNA4, LDLR, LRP1, LRP2, LRP5, LRP8, SCARB1, SORL1, VLDLR*)(*96*). Analyses were conducted both on the integrated dataset and the datasets split by treatment (chAdu vs. IgG), after which results were merged for comparative evaluation. Gene set enrichment analysis was conducted on microglia and astrocytes using the Escape package and MSigDB Hallmark, CP-Reactome, and GO-Biological Processes gene sets (*97–104*).

### Xenium Spatial Transcriptomics Data Processing

Xenium spatial transcriptomics data was obtained using Xenium analyzer and our custom 480 gene panel including cell type marker genes, immune function, vascular, and metabolic genes for in depth spatial transcript characterization. Data was run through Xenium Ranger with a default 5µm nuclear expansion. Cortex, hippocampus, thalamus, white matter tracts (WM), and hypothalamus were annotated in Xenium Explorer (v4.0.0) and exported along with cell IDs and coordinates. Cells were purified using RCTD/SPLIT pipeline for decontamination followed by removal of low count cells (cells with nCount < 5) (*105, 106*). Processing and annotation of cell types was done as described above, with 30 PCs and 480 features, clustering resolution of 0.3. Image coregistration and visualization was done as previously described for analysis of transcripts and plaque proximity gene correlations (*107*). Differential abundance was calculated using propeller, contained within speckle v1.10.0 (*108*). Neighborhood enrichment analysis was done as previously described, and visualized in network plots using the ggforce package for pseudo neighborhood generation. Banksy v1.6.0 algorithm was used to generate spatial domains, with lambda = 0.8 to favor spatial characteristics(*43*). Domains were then identified based on their spatial distribution, cell-type composition, and gene expression. Three domains of interest were selected based on findings.

### Imaging and Image Analysis

Unless otherwise noted, slides were imaged using a Zeiss Axioscan Z7 slide scanner. CD206, Dectin, and ApoE slides were imaged using the Nikon W1-SoRA spinning disk confocal microscope to allow for greater resolution of vascular interfaces. All fluorescent images were taken at 40x zoom. Images were quantified in Indica Labs HALO software.

### Statistics

All statistics were conducted in GraphPad Prism 10. Sexes were pooled and data analyzed using Two-way ANOVA with Tukey’s post-hoc correction unless otherwise noted with significance at p < 0.05 (p-value * <.05, ** <.005 *** <.001). Outlier analyses were conducted where appropriate using ROUT >1%, outliers were assessed by a blinded lab member and removed where appropriate.

## Acknowledgments

We would like to thank the University of Kentucky Light Microscopy core, and manager Dr. Xu Fu for their invaluable support. We would also like to thank the University of Kentucky MRI and Spectroscopy core, particularly Dr. David Powell and Stephen Dundon for their help with sequence generation. We would also like to thank Drs. Jamie Sturgill and Richard King for invaluable guidance during this project.

## Author contributions

LAJ, JMM, PRT and AVP conceived the experiment, designed the study, and secured funding. AVP, LAJ, JMM, JLF, CCL, KS and JAM analyzed the data. PRT and SP provided chAdu antibody and assisted with MRI hypointensity analysis. Animal work was done by AVP, GH, GLN, and SMM. The majority of experiments were performed by AVP with guidance or assistance from GLN, IOS, LMS, JLF, DG, DS, SMO, and DA. Transcriptomics analyses were performed by CCL, KS, AVP, JAM and JMM. Technical expertise and guidance regarding data analysis and interpretation was given by SLM, LRG, and PRT. Manuscript was written by AVP and LAJ with input from all co-authors.

## Data and materials availability

All data needed to evaluate the conclusions in the paper are present in the paper and/or the Supplementary Materials. The single-cell RNA-sequencing and Xenium spatial transcriptomics datasets generated in this study have been deposited in the NCBI Gene Expression Omnibus (GEO). Analysis code is available on GitHub. Additional data and materials are available from the corresponding authors upon reasonable request.

## List of Supplementary Materials

Fig. S1 Amyloid Burden by region

Fig. S2 Amyloid MSD Quantification and Analyses

Fig. S3 Prussian Blue and MRI additional quantifications

Fig. S4 CellChat and MultiNicheNet

Fig. S5 Xenium brain characterization

Fig. S6 Xenium microglia and astrocyte characterization

Fig. S7 scRNAseq cell type annotations and GSVA

Fig. S8 Banksy Domains and Network plots

Supplementary Data 1 – Amyloid Statistical Testing

Supplementary Data 2 – Prussian Blue/MRI statistical testing

Supplementary Data 3 – Gliosis Statistical Testing

## List of Abbreviations

AD: Alzheimer’s Disease
ARIA: Amyloid Related Imaging
Abnormalities APOE: Apolipoprotein E
CAA: Cerebral Amyloid Angiopathy
chAdu: Chimeric Aducanumab
MRI: Magnetic Resonance Imaging
SWI: Susceptibility Weighted Imaging
ACCF: Allen Common Coordinate Framework
IBA1: Ionizing Binding Adapter 1
GFAP: Glial Fibrillary Acidic Protein
CNS: Central Nervous System
LAMP1: Lysosomal Associated Membrane Protein 1
P2RY12: Purinergic Receptor P2Y12
BAM: Border Associated Macrophages
scRNAseq: Single Cell RNA Sequencing
ST: Spatial Transcriptomics
Hm: Homeostatic Microglia
DAM: Disease Associated Microglia
ARM: Activated Response Microglia
IRM: Interferon Response Microglia
IRRA: Interferon Responsive Reactive Astrocyte
MHC: Major Histocompatibility Complex

---

**Supplementary Figure 1:**
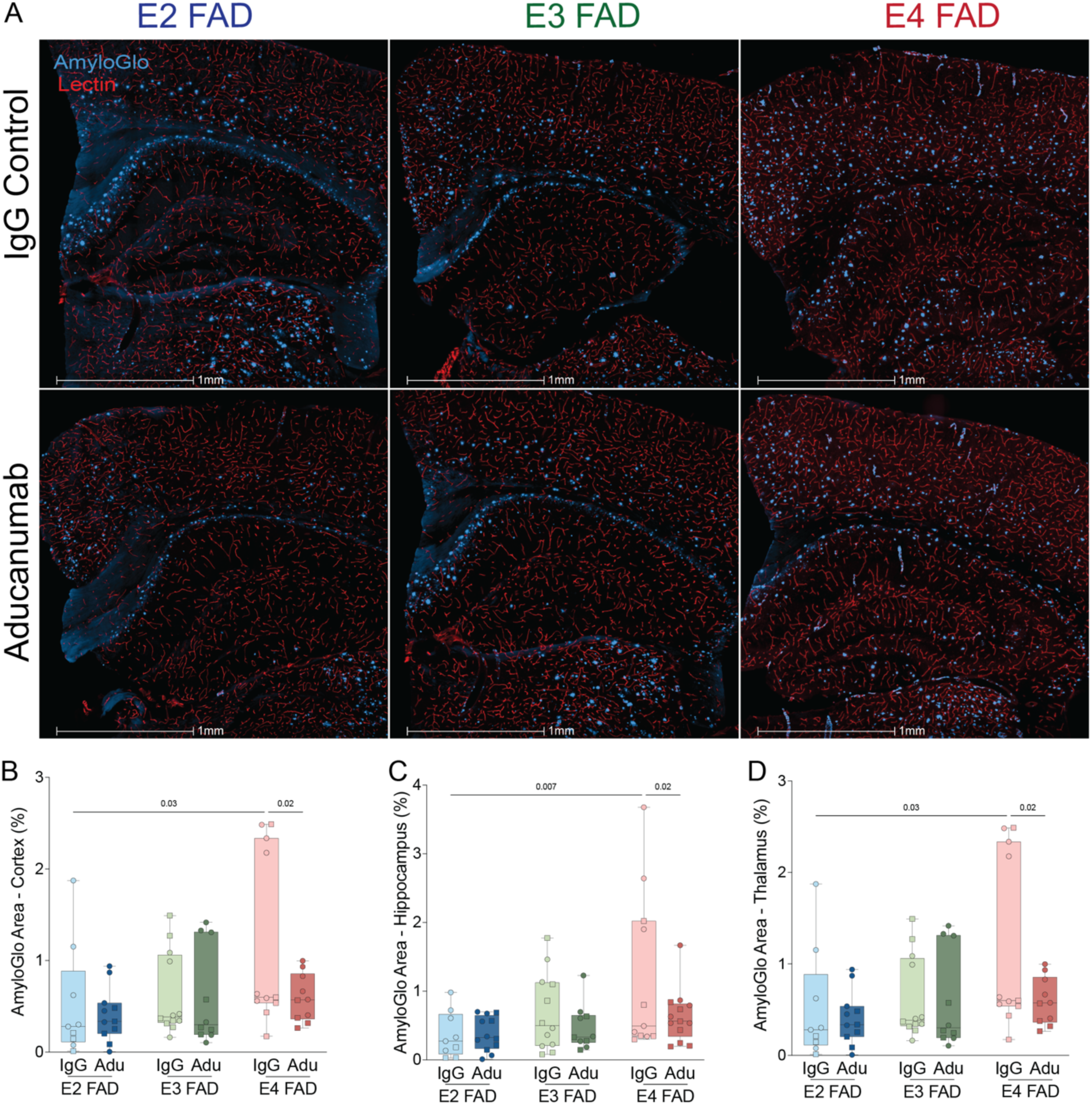
Regional quantification of amyloid burden. A) Representative image of amyloid plaque staining (AmyloGlo, blue) and vessels (Isolectin, red). B-D) Quantification of amyloid burden in cortex, hippocampus, and thalamus, respectively as percent area coverage.

**Supplementary Figure 2:**
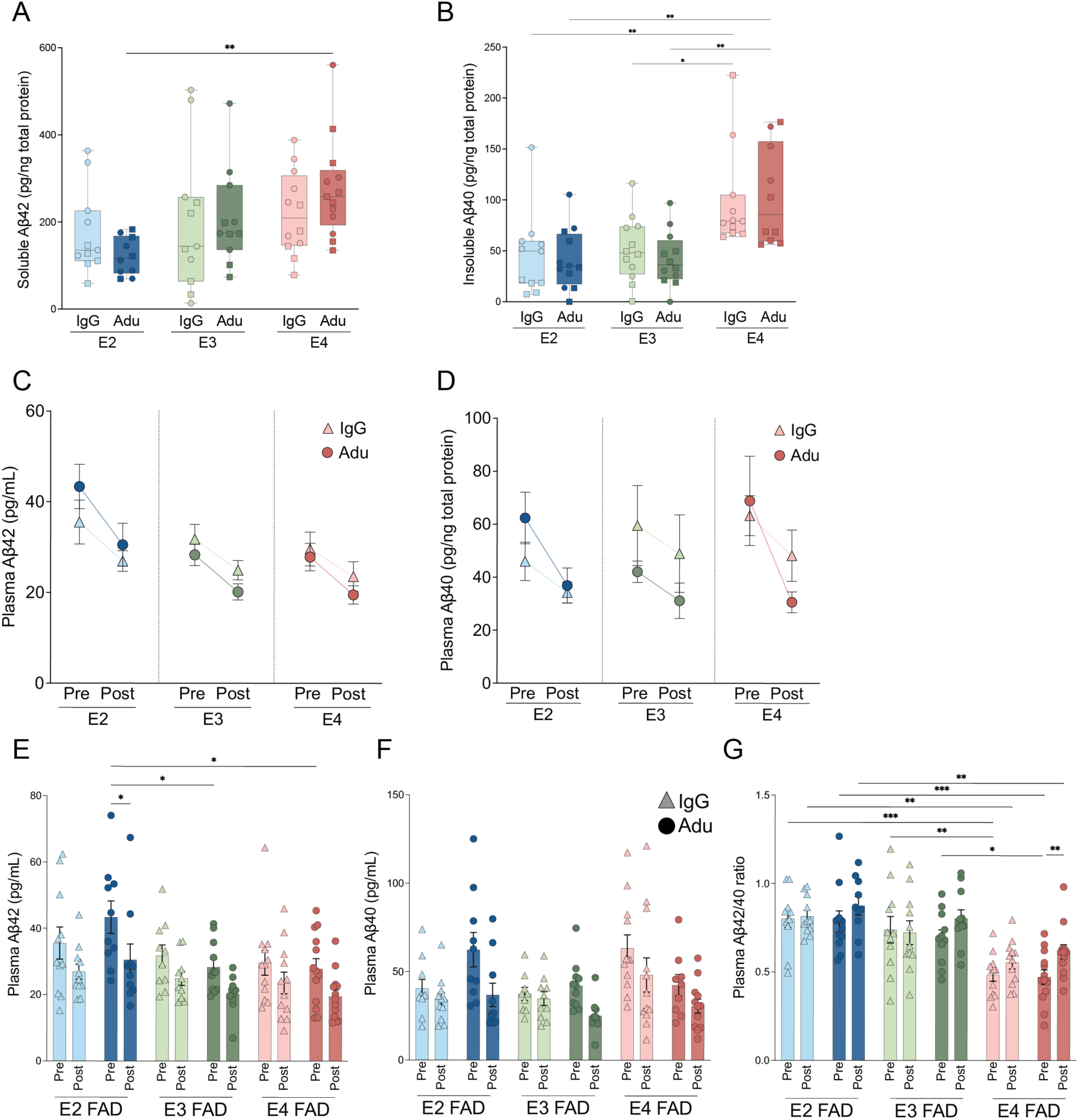
ELISA quantification of Aβ40 and 42 in plasma and brain tissue. A-B) Quantification of Aβ42 and Aβ40 in insoluble brain fraction. C-D) Quantification of plasma Aβ42 and plasma Aβ40 over treatment course. E-G) Analysis breakdown of pre-post treatment plasma Aβ42, Aβ40, Aβ42/40 ratio quantification, respectively.

**Supplementary Figure 3:**
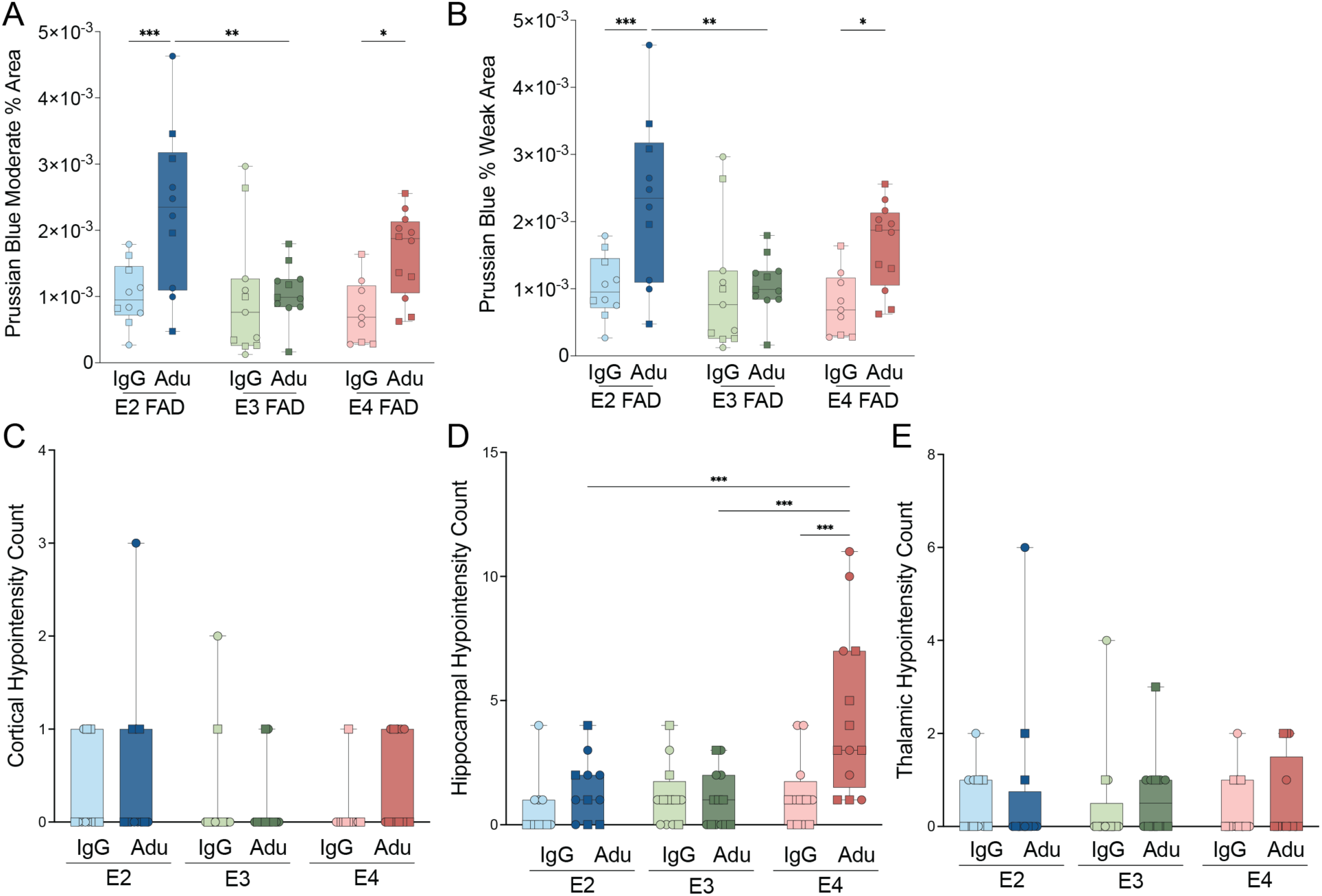
Additional quantifications of Prussian Blue and regional hypointensity occurrence on MRI. A-B) Quantification of Prussian blue moderate area (A) and weak area (B). C-E) Hypointensity occurrence in cortex, hippocampus, and thalamus, respectively.

**Supplementary Figure 4:**
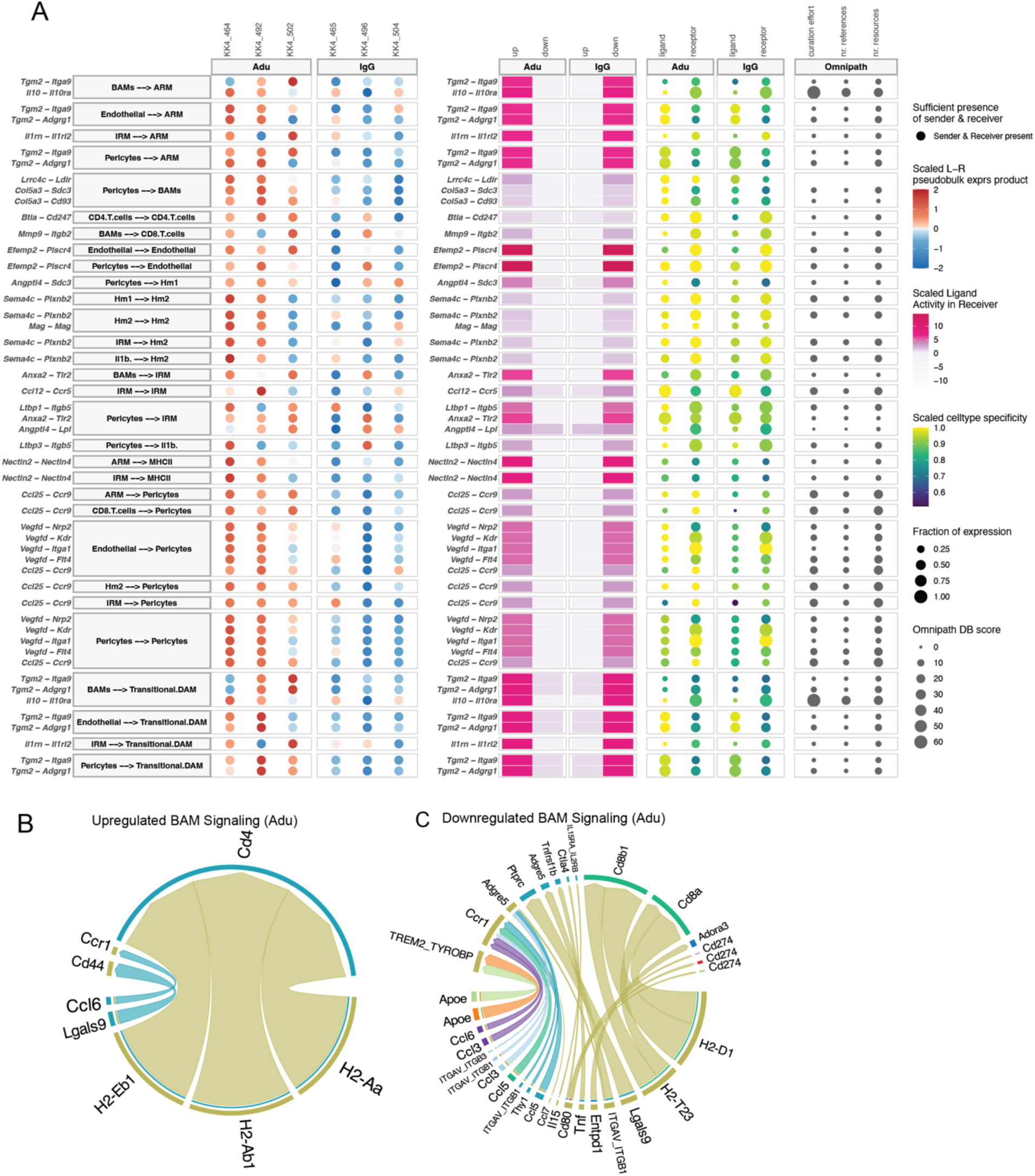
MultiNicheNet and CellChat signaling between vascular and immune cells. A) MultiNicheNet diagram depicting signaling pathways up/downregulated in single cell RNA sequencing samples with pathway and celltypes on left, regulation status in the middle and cellular specificity on right. B-C) Up and downregulated signaling after chAdu treatment in border associated macrophages.

**Supplementary Figure 5:**
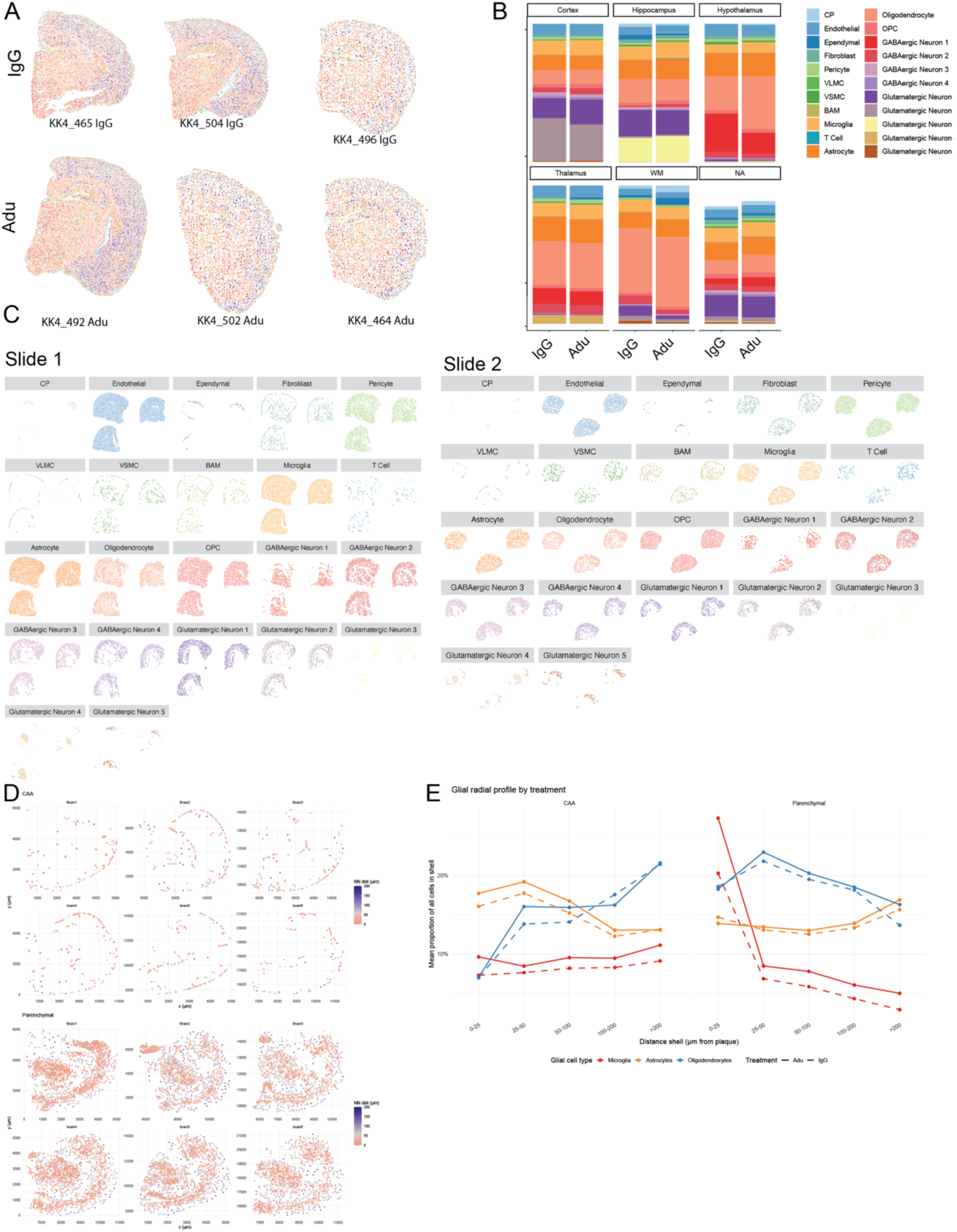
Xenium data characterization. A) Representative images of 6 brains used for Xenium ST. B) Cell type proportions split by treatment and region. C) Cell type spatial distribution breakdown. D) Spatial distribution of plaques in each brain sample. E) Composition of peri-plaque and peri-caa niche in terms of microglia, astrocytes, and oligodendrocytes by distance.

**Supplementary Figure 6:**
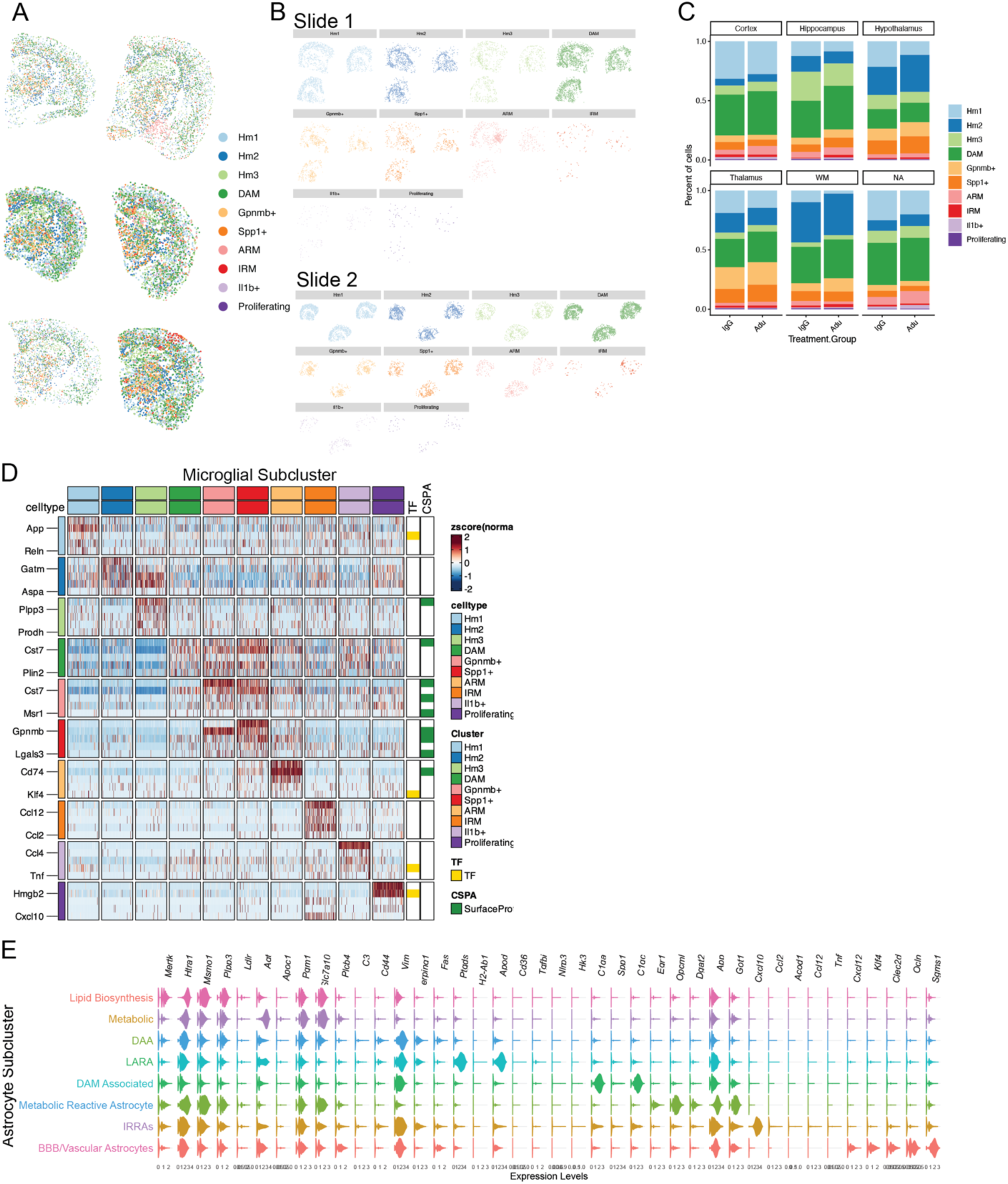
Breakdown of Astrocytes and microglia in Xenium spatial transcriptomics and cellular annotations. A) Representative image of Microglial breakdown. B) Microglial subcluster spatial breakdown. C) Relative proportions of microglial subclusters by region and treatment group. D) expression of canonical microglial state markers by subcluster. E) Expression of canonical astrocyte state markers by subcluster.

**Supplementary Figure 7:**
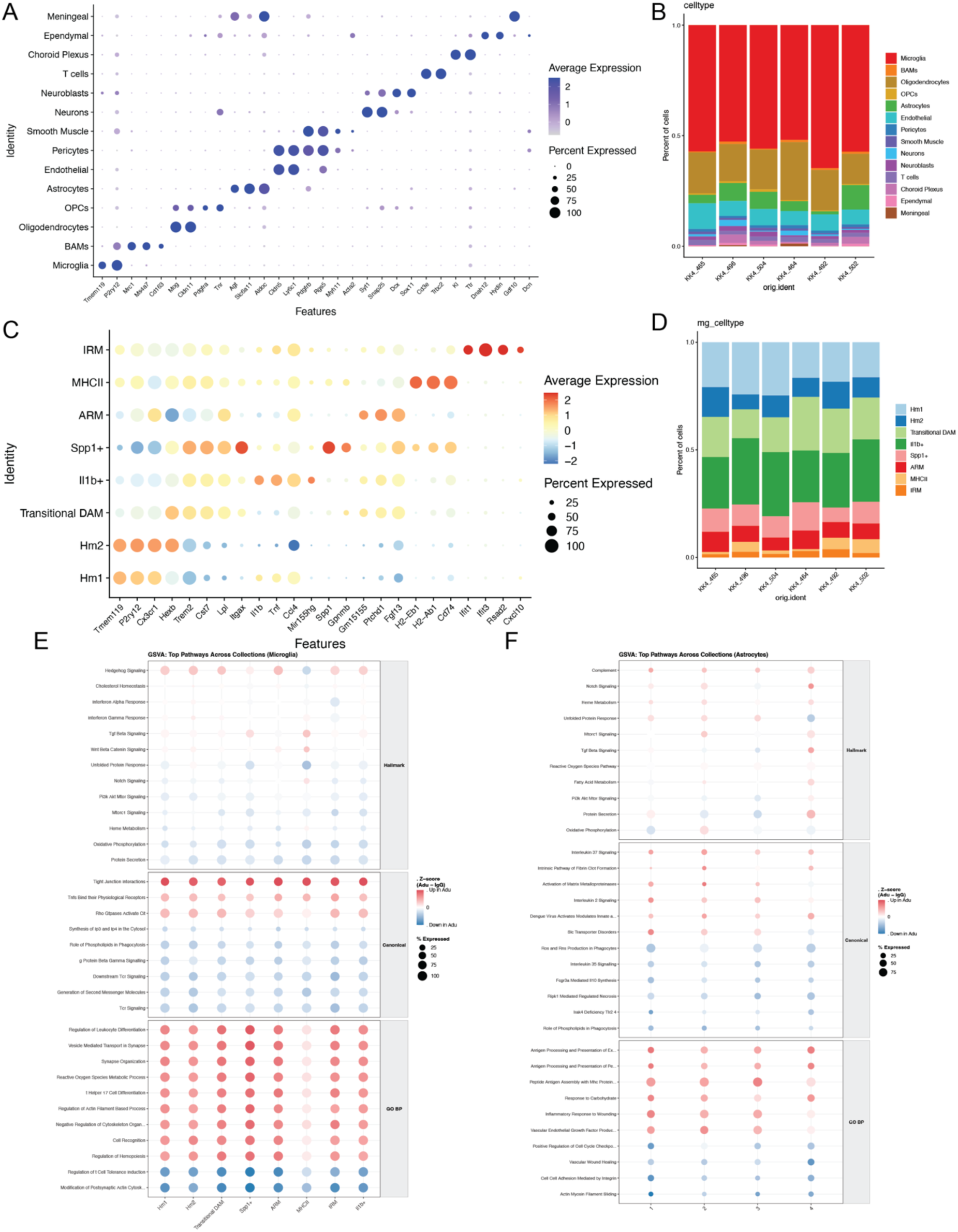
Cell type annotations in scRNAseq data and Gene set variation analysis. A) Cell type x canonical marker matrix dotplot. B) Distribution of celltypes in scRNAseq data. C) Microglial subcluster x canonical marker matrix dotplot. D) Proportions of microglial subclusters in scRNAseq data. E) Gene set variation analysis across 3 pathway sets in microglia (E) and astrocytes (F).

**Supplementary Figure 8:**
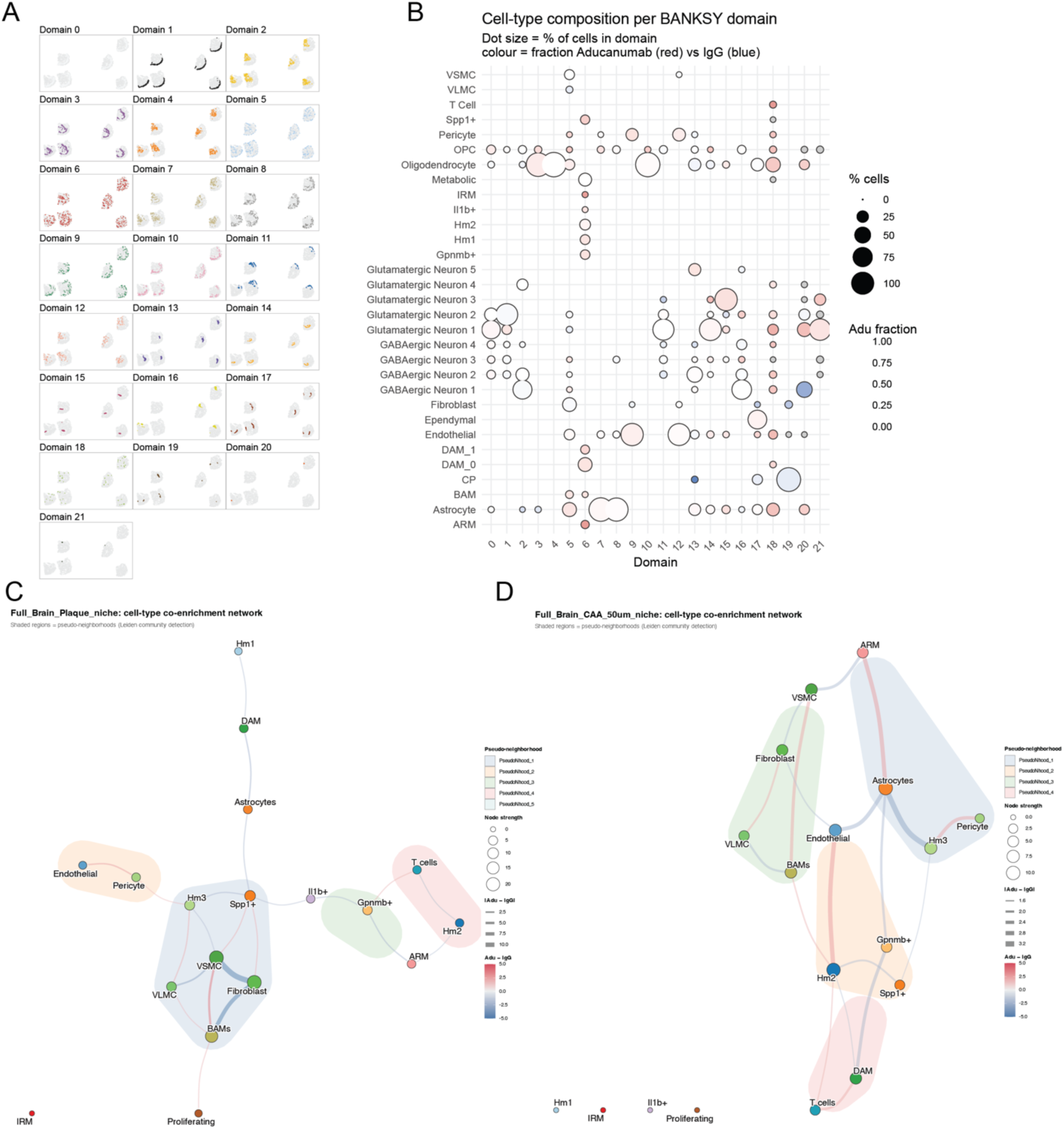
BANKSY Domain compositions and Networks. A) Spatial distribution of BANKSY domains. B) Cell type composition of BANKSY domains. C-D) Spatial neighborhood network plot of parenchymal plaque spatial neighborhood (C) and CAA spatial neighborhood (D).

## References and Notes

1. 2024 Alzheimer’s disease facts and figures. Alzheimer’s Dement. 20, 3708–3821 (2024).

2. S. Salloway, S. Chalkias, F. Barkhof, P. Burkett, J. Barakos, D. Purcell, J. Suhy, F. Forrestal, Y. Tian, K. Umans, G. Wang, P. Singhal, S. B. Haeberlein, K. Smirnakis, Amyloid-Related Imaging Abnormalities in 2 Phase 3 Studies Evaluating Aducanumab in Patients With Early Alzheimer Disease. JAMA Neurol. 79, 13–21 (2022).

3. C. H. v. Dyck, C. J. Swanson, P. Aisen, R. J. Bateman, C. Chen, M. Gee, M. Kanekiyo, D. Li, L. Reyderman, S. Cohen, L. Froelich, S. Katayama, M. Sabbagh, B. Vellas, D. Watson, S. Dhadda, M. Irizarry, L. D. Kramer, T. Iwatsubo, Lecanemab in Early Alzheimer’s Disease. N. Engl. J. Med. 388, 9–21 (2022).

4. J. R. Sims, J. A. Zimmer, C. D. Evans, M. Lu, P. Ardayfio, J. Sparks, A. M. Wessels, S. Shcherbinin, H. Wang, E. S. M. Nery, E. C. Collins, P. Solomon, S. Salloway, L. G. Apostolova, O. Hansson, C. Ritchie, D. A. Brooks, M. Mintun, D. M. Skovronsky, T.-A. Investigators, R. Abreu, P. Agarwal, P. Aggarwal, M. Agronin, A. Allen, D. Altamirano, G. Alva, J. Andersen, A. Anderson, D. Anderson, J. Arnold, T. Asada, Y. Aso, V. Atit, R. Ayala, M. Badruddoja, H. Badzio-jagiello, M. Bajacek, D. Barton, D. Bear, S. Benjamin, R. Bergeron, P. Bhatia, S. Black, A. Block, M. Bolouri, W. Bond, J. Bouthillier, S. Brangman, B. Brew, S. Brisbin, T. Brisken, A. Brodtmann, M. Brody, J. Brosch, C. Brown, P. Brownstone, S. Bukowczan, J. Burns, A. Cabrera, H. Capote, A. Carrasco, J. C. Yepez, E. Chavez, H. Chertkow, U. Chyrchel-paszkiewicz, A. Ciabarra, E. Clemmons, D. Cohen, R. Cohen, I. Cohen, M. Concha, B. Costell, D. Crimmins, Y. Cruz-pagan, A. Cueli, R. Cupelo, M. Czarnecki, D. Darby, P. l. j. Dautzenberg, P. D. Deyn, J. D. L. Gandara, K. Deck, D. Dibenedetto, M. Dibuono, E. Dinnerstein, A. Dirican, S. Dixit, J. Dobryniewski, R. Drake, P. Drysdale, R. Duara, J. Duffy, A. Ellenbogen, V. Faradji, M. Feinberg, R. Feldman, S. Fishman, S. Flitman, C. Forchetti, I. Fraga, A. Frank, B. Frishberg, H. Fujigasaki, H. Fukase, I. Fumero, K. Furihata, C. Galloway, R. Gandhi, K. George, M. Germain, D. Gitelman, N. Goetsch, D. Goldfarb, M. Goldstein, L. Goldstick, Y. G. Rojas, I. Goodman, D. Greeley, C. Griffin, E. Grigsby, D. Grosz, K. Hafner, D. Hart, S. Henein, B. Herskowitz, S. Higashi, Y. Higashi, G. Ho, J. Hodgson, M. Hohenberg, L. Hollenbeck, R. Holub, T. Hori, J. Hort, J. Ilkowski, K. J. Ingram, M. Isaac, M. Ishikawa, L. Janu, M. Johnston, W. Julio, W. Justiz, T. Kaga, T. Kakigi, M. Kalafer, M. Kamijo, J. Kaplan, M. Karathanos, S. Katayama, S. Kaul, A. Keegan, D. Kerwin, U. Khan, A. Khan, N. Kimura, G. Kirk, G. Klodowska, H. Kowa, C. Kutz, J. Kwentus, R. Lai, A. Lall, M. Lawrence, E. Lee, R. Leon, G. Linker, P. Lisewski, J. Liss, C. Liu, S. Losk, E. Lukaszyk, J. Lynch, S. Macfarlane, J. Macsweeney, N. Mannering, O. Markovic, D. Marks, J. Masdeu, Y. Matsui, K. Matsuishi, P. McAllister, B. McConnehey, A. McElveen, L. McGill, A. Mecca, M. Mega, J. Mensah, A. Mickielewicz, A. Minaeian, B. Mocherla, C. Murphy, P. Murphy, H. Nagashima, A. Nair, M. Nair, J. Nardandrea, M. Nash, Z. Nasreddine, Y. Nishida, J. Norton, L. Nunez, J. Ochiai, T. Ohkubo, Y. Okamura, E. Okorie, E. Olivera, J. O’Mahony, O. Omidvar, D. Ortiz-Cruz, A. Osowa, M. Papka, A. Parker, P. Patel, A. Patel, M. Patel, C. Patry, E. Peckham, M. Pfeffer, A. Pietras, M. Plopper, A. Porsteinsson, R. P. Robitaille, N. Prins, O. Puente, M. Ratajczak, M. Rhee, A. Ritter, R. Rodriguez, L. R. Ables, J. Rojas, J. Ross, P. Royer, J. Rubin, D. Russell, S. M. Rutgers, S. Rutrick, M. Sadowski, B. Safirstein, T. Sagisaka, D. Scharre, L. Schneider, C. Schreiber, M. Schrift, P. Schulz, H. Schwartz, J. Schwartzbard, J. Scott, L. Selem, P. Sethi, S. Sha, K. Sharlin, S. Sharma, T. Shiovitz, R. Shiwach, M. Sladek, B. Sloan, A. Smith, P. Solomon, E. Sorial, E. Sosa, M. Stedman, S. Steen, L. Stein, A. Stolyar, J. Stoukides, S. Sudoh, J. Sutton, J. Syed, K. Szigeti, H. Tachibana, Y. Takahashi, A. Tateno, J. D. Taylor, K. Taylor, O. Tcheremissine, A. Thebaud, S. Thein, L. Thurman, S. Toenjes, H. Toji, M. Toma, D. Tran, P. Trueba, M. Tsujimoto, R. Turner, A. Uchiyama, D. Ussorowska, S. Vaishnavi, E. Valor, J. Vandersluis, A. Vasquez, J. Velez, C. Verghese, K. Vodickova-borzova, D. Watson, D. Weidman, D. Weisman, A. White, K. Willingham, I. Winkel, P. Winner, J. Winston, A. Wolff, H. Yagi, H. Yamamoto, S. Yathiraj, Y. Yoshiyama, M. Zboch, Donanemab in Early Symptomatic Alzheimer Disease. JAMA 330, 512–527 (2023).

5. H. Hampel, A. Elhage, M. Cho, L. G. Apostolova, J. A. R. Nicoll, A. Atri, Amyloid-related imaging abnormalities (ARIA): radiological, biological and clinical characteristics. Brain, (2023).

6. E. M. Reiman, J. F. Arboleda-Velasquez, Y. T. Quiroz, M. J. Huentelman, T. G. Beach, R. J. Caselli, Y. Chen, Y. Su, A. J. Myers, J. Hardy, J. P. Vonsattel, S. G. Younkin, D. A. Bennett, P. L. D. Jager, E. B. Larson, P. K. Crane, C. D. Keene, M. I. Kamboh, J. K. Kofler, L. Duque, J. R. Gilbert, H. E. Gwirtsman, J. D. Buxbaum, D. W. Dickson, M. P. Frosch, B. F. Ghetti, K. L. Lunetta, L.-S. Wang, B. T. Hyman, W. A. Kukull, T. Foroud, J. L. Haines, R. P. Mayeux, M. A. Pericak-Vance, J. A. Schneider, J. Q. Trojanowski, L. A. Farrer, G. D. Schellenberg, G. W. Beecham, T. J. Montine, G. R. Jun, T. A. s. D. G. Consortium, E. Abner, P. M. Adams, M. S. Albert, R. L. Albin, L. G. Apostolova, S. E. Arnold, S. Asthana, C. S. Atwood, C. T. Baldwin, R. C. Barber, L. L. Barnes, S. Barral, J. T. Becker, D. Beekly, E. H. Bigio, T. D. Bird, D. Blacker, B. F. Boeve, J. D. Bowen, A. Boxer, J. R. Burke, J. M. Burns, N. J. Cairns, L. B. Cantwell, C. Cao, C. S. Carlson, C. M. Carlsson, R. M. Carney, M. M. Carrasquillo, H. C. Chui, D. H. Cribbs, E. A. Crocco, C. Cruchaga, C. DeCarli, M. Dick, R. S. Doody, R. Duara, N. Ertekin-Taner, D. A. Evans, K. M. Faber, T. J. Fairchild, K. B. Fallon, D. W. Fardo, M. R. Farlow, S. Ferris, D. R. Galasko, M. Gearing, D. H. Geschwind, V. Ghisays, A. M. Goate, N. R. Graff-Radford, R. C. Green, J. H. Growdon, H. Hakonarson, R. L. Hamilton, K. L. Hamilton-Nelson, L. E. Harrell, L. S. Honig, R. M. Huebinger, C. M. Hulette, G. P. Jarvik, L.-W. Jin, A. Karydas, M. J. Katz, J. S. K. Kauwe, J. A. Kaye, R. Kim, N. W. Kowall, J. H. Kramer, B. W. Kunkle, A. P. Kuzma, F. M. LaFerla, J. J. Lah, Y. Y. Leung, J. B. Leverenz, A. I. Levey, G. Li, A. P. Lieberman, R. B. Lipton, O. L. Lopez, C. G. Lyketsos, J. Malamon, D. C. Marson, E. R. Martin, F. Martiniuk, D. C. Mash, E. Masliah, W. C. McCormick, S. M. McCurry, A. N. McDavid, S. McDonough, A. C. McKee, M. Mesulam, B. L. Miller, C. A. Miller, J. W. Miller, J. C. Morris, S. Mukherjee, A. C. Naj, S. O’Bryant, J. M. Olichney, J. E. Parisi, H. L. Paulson, E. Peskind, R. C. Petersen, A. Pierce, W. W. Poon, H. Potter, L. Qu, J. F. Quinn, A. Raj, M. Raskind, B. Reisberg, J. S. Reisch, C. Reitz, J. M. Ringman, E. D. Roberson, E. Rogaeva, H. J. Rosen, R. N. Rosenberg, D. R. Royall, M. A. Sager, M. Sano, A. J. Saykin, L. S. Schneider, W. W. Seeley, A. G. Smith, J. A. Sonnen, S. Spina, P. S. George-Hyslop, R. A. Stern, R. H. Swerdlow, R. E. Tanzi, J. C. Troncoso, D. W. Tsuang, O. Valladares, V. M. V. Deerlin, L. J. V. Eldik, B. N. Vardarajan, H. V. Vinters, S. Weintraub, K. A. Welsh-Bohmer, K. C. Wilhelmsen, J. Williamson, T. S. Wingo, R. L. Woltjer, C. B. Wright, C.-K. Wu, C.-E. Yu, L. Yu, Y. Zhao, Exceptionally low likelihood of Alzheimer’s dementia in APOE2 homozygotes from a 5,000-person neuropathological study. Nat. Commun. 11, 667 (2020).

7. C.-C. Liu, C.-C. Liu, T. Kanekiyo, H. Xu, G. Bu, Apolipoprotein E and Alzheimer disease: risk, mechanisms and therapy. Nat. Rev. Neurol. 9, 106–118 (2013).

8. J. Raber, Y. Huang, J. W. Ashford, ApoE genotype accounts for the vast majority of AD risk and AD pathology. Neurobiol. Aging 25, 641–650 (2004).

9. E. Solopova, W. Romero-Fernandez, H. Harmsen, L. Ventura-Antunes, E. Wang, A. Shostak, J. Maldonado, M. J. Donahue, D. Schultz, T. M. Coyne, A. Charidimou, M. Schrag, Fatal iatrogenic cerebral β-amyloid-related arteritis in a woman treated with lecanemab for Alzheimer’s disease. Nature Communications 14, 8220 (2023).

10. N. J. Reish, P. Jamshidi, B. Stamm, M. E. Flanagan, E. Sugg, M. Tang, K. L. Donohue, M. McCord, C. Krumpelman, M.-M. Mesulam, R. Castellani, S. H. Y. Chou, Multiple Cerebral Hemorrhages in a Patient Receiving Lecanemab and Treated with t-PA for Stroke. N. Engl. J. Med. 388, 478–479 (2023).

11. J. N. Briard, A. Duquette, R. Cayrol, S. Lapalme-Remis, Refractory Status Epilepticus in a Patient With Aducanumab-Induced Amyloid-Related Imaging Abnormalities. Neurology 103, e209582 (2024).

12. L. Antolini, J. C. DiFrancesco, M. Zedde, G. Basso, A. Arighi, A. Shima, A. Cagnin, M. Caulo, R. Carare, A. Charidimou, M. Cirillo, V. D. Lazzaro, C. Ferrarese, A. Giossi, D. Inzitari, M. Marcon, R. Marconi, M. Ihara, R. Nitrini, B. Orlandi, A. Padovani, R. Pascarella, F. Perini, G. Perini, M. Sessa, E. Scarpini, F. Tagliavini, R. Valenti, J. F. Vázquez-Costa, A. Villarejo-Galende, Y. Hagiwara, N. Ziliotto, F. Piazza, Spontaneous ARIA-like Events in Cerebral Amyloid Angiopathy–Related Inflammation: A Multicenter Prospective Longitudinal Cohort Study. Neurology 97, 10.1212/WNL.0000000000012778 (2021).

13. E. M. Weekman, T. L. Sudduth, C. N. Caverly, T. J. Kopper, O. W. Phillips, D. K. Powell, D. M. Wilcock, Reduced Efficacy of Anti-Aβ Immunotherapy in a Mouse Model of Amyloid Deposition and Vascular Cognitive Impairment Comorbidity. J. Neurosci. 36, 9896–9907 (2016).

14. D. M. Wilcock, A. Rojiani, A. Rosenthal, S. Subbarao, M. J. Freeman, M. N. Gordon, D. Morgan, Passive immunotherapy against Aβ in aged APP-transgenic mice reverses cognitive deficits and depletes parenchymal amyloid deposits in spite of increased vascular amyloid and microhemorrhage. J. Neuroinflammation 1, 24 (2004).

15. D. M. Wilcock, A. Rojiani, A. Rosenthal, G. Levkowitz, S. Subbarao, J. Alamed, D. Wilson, N. Wilson, M. J. Freeman, M. N. Gordon, D. Morgan, Passive Amyloid Immunotherapy Clears Amyloid and Transiently Activates Microglia in a Transgenic Mouse Model of Amyloid Deposition. J. Neurosci. 24, 6144–6151 (2004).

16. S. D. Mesquita, Z. Papadopoulos, T. Dykstra, L. Brase, F. G. Farias, M. Wall, H. Jiang, C. D. Kodira, K. A. d. Lima, J. Herz, A. Louveau, D. H. Goldman, A. F. Salvador, S. Onengut-Gumuscu, E. Farber, N. Dabhi, T. Kennedy, M. G. Milam, W. Baker, I. Smirnov, S. S. Rich, D. I. A. Network, B. A. Benitez, C. M. Karch, R. J. Perrin, M. Farlow, J. P. Chhatwal, D. M. Holtzman, C. Cruchaga, O. Harari, J. Kipnis, Meningeal lymphatics affect microglia responses and anti-Aβ immunotherapy. Nature 593, 255–260 (2021).

17. P. Pikus, R. S. Turner, G. W. Rebeck, Mouse models of Anti-Aβ immunotherapies. Mol. Neurodegener. 20, 57 (2025).

18. W. Zago, S. Schroeter, T. Guido, K. Khan, P. Seubert, T. Yednock, D. Schenk, K. M. Gregg, D. Games, F. Bard, G. G. Kinney, Vascular alterations in PDAPP mice after anti-Aβ immunotherapy: Implications for amyloid-related imaging abnormalities. Alzheimer’s Dement. 9, S105–S115 (2013).

19. X. Taylor, I. M. Clark, G. J. Fitzgerald, H. Oluoch, J. T. Hole, R. B. DeMattos, Y. Wang, F. Pan, Amyloid-β (Aβ) immunotherapy induced microhemorrhages are associated with activated perivascular macrophages and peripheral monocyte recruitment in Alzheimer’s disease mice. Mol. Neurodegener. 18, 59 (2023).

20. X. Taylor, H. N. Noristani, G. J. Fitzgerald, H. Oluoch, N. Babb, T. McGathey, L. Carter, J. T. Hole, P. N. Lacor, R. B. DeMattos, Y. Wang, Amyloid-β (Aβ) immunotherapy induced microhemorrhages are linked to vascular inflammation and cerebrovascular damage in a mouse model of Alzheimer’s disease. Mol. Neurodegener. 19, 77 (2024).

21. F. Liao, T. J. Zhang, H. Jiang, K. B. Lefton, G. O. Robinson, R. Vassar, P. M. Sullivan, D. M. Holtzman, Murine versus human apolipoprotein E4: differential facilitation of and co-localization in cerebral amyloid angiopathy and amyloid plaques in APP transgenic mouse models. Acta Neuropathol. Commun. 3, 70 (2015).

22. D. Balu, A. C. Valencia-Olvera, Z. Islam, C. Mielczarek, A. Hansen, T. M. P. Ramos, J. York, M. J. LaDu, L. M. Tai, APOE genotype and sex modulate Alzheimer’s disease pathology in aged EFAD transgenic mice. Front. Aging Neurosci. 15, 1279343 (2023).

23. L. M. Tai, D. Balu, E. Avila-Munoz, L. Abdullah, R. Thomas, N. Collins, A. C. Valencia-Olvera, M. J. LaDu, EFAD transgenic mice as a human APOE relevant preclinical model of Alzheimerŉs disease[S]. J. Lipid Res. 58, 1733–1755 (2017).

24. K. L. Youmans, L. M. Tai, E. Nwabuisi-Heath, L. Jungbauer, T. Kanekiyo, M. Gan, J. Kim, W. A. Eimer, S. Estus, G. W. Rebeck, E. J. Weeber, G. Bu, C. Yu, M. J. LaDu, APOE4-specific Changes in Aβ Accumulation in a New Transgenic Mouse Model of Alzheimer Disease*. J. Biol. Chem. 287, 41774–41786 (2012).

25. J. E. Pankiewicz, J. Baquero-Buitrago, S. Sanchez, J. Lopez-Contreras, J. Kim, P. M. Sullivan, D. M. Holtzman, M. J. Sadowski, APOE Genotype Differentially Modulates Effects of Anti-Aβ, Passive Immunization in APP Transgenic Mice. Mol. Neurodegener. 12, 12 (2017).

26. M. Xiong, H. Jiang, J. R. Serrano, E. R. Gonzales, C. Wang, M. Gratuze, R. Hoyle, N. Bien-Ly, A. P. Silverman, P. M. Sullivan, R. J. Watts, J. D. Ulrich, G. J. Zipfel, D. M. Holtzman, APOE immunotherapy reduces cerebral amyloid angiopathy and amyloid plaques while improving cerebrovascular function. Sci. Transl. Med. 13, (2021).

27. A. Millet, J. H. Ledo, S. F. Tavazoie, An exhausted-like microglial population accumulates in aged and APOE4 genotype Alzheimer’s brains. Immunity, (2023).

28 . L. d. Weerd, S. Hummel, S. A. Müller, I. Paris, T. Sandmann, M. Eichholtz, R. Gröger, A. L. Englert, S. Wagner, C. Ha, S. S. Davis, V. Warkins, D. Xia, B. Nuscher, A. Berghofer, M. Reich, A. F. Feiten, K. Schlepckow, M. Willem, S. F. Lichtenthaler, J. W. Lewcock, K. M. Monroe, M. Brendel, C. Haass, Early intervention anti-Aβ immunotherapy attenuates microglial activation without inducing exhaustion at residual plaques. Mol. Neurodegener. 20, 92 (2025).

29. P. R. Territo, S. K. Quinney, A. R. Masters, K. A. Haynes, Z. A. Cope, G. Little, S. P. Williams, J. A. Meyer, J. Peters, L. Figueiredo, S. C. Persohn, A. A. Bedwell, K. Eldridge, R. Speedy, N. T. Seyfried, K. D. Onos, M. Sasner, G. R. Howell, G. W. Carter, A. L. Oblak, B. T. Lamb, S. J. S. Rizzo, Pharmacodynamics Assessment of Aducanumab in 5XFAD mice: A MODEL-AD PTC Study. Alzheimer’s Dement. 18, (2022).

30. E. H. Corder, A. M. Saunders, W. J. Strittmatter, D. E. Schmechel, P. C. Gaskell, G. W. Small, A. D. Roses, J. L. Haines, M. A. Pericak-Vance, Gene Dose of Apolipoprotein E Type 4 Allele and the Risk of Alzheimer’s Disease in Late Onset Families. Science 261, 921–923 (1993).

31. S. Knudtzon, B. E. Kirsebom, L. Pålhaugen, F. Bettella, B. Gísladóttir, S. Bahrami, L. Athanasiu, A. Rongve, A. Nakling, I. S. Almdahl, L. Kalheim, J. A. Jarholm, G. R. Grøntvedt, R. E. Skogseth, D. Aarsland, K. Waterloo, O. A. Andreassen, T. Fladby, K. Nordengen, Genotype–phenotype interaction in Alzheimer’s disease immune activation. Alzheimer’s Dement. 22, e70978 (2026).

32. J. C. Morris, C. M. Roe, C. Xiong, A. M. Fagan, A. M. Goate, D. M. Holtzman, M. A. Mintun, APOE predicts amyloid-beta but not tau Alzheimer pathology in cognitively normal aging. Ann. Neurol. 67, 122–131 (2010).

33. E. M. Reiman, K. Chen, X. Liu, D. Bandy, M. Yu, W. Lee, N. Ayutyanont, J. Keppler, S. A. Reeder, J. B. S. Langbaum, G. E. Alexander, W. E. Klunk, C. A. Mathis, J. C. Price, H. J. Aizenstein, S. T. DeKosky, R. J. Caselli, Fibrillar amyloid-β burden in cognitively normal people at 3 levels of genetic risk for Alzheimer’s disease. Proc. Natl. Acad. Sci. 106, 6820–6825 (2009).

34. D. Balu, A. J. Karstens, E. Loukenas, J. M. Weng, J. M. York, A. C. Valencia-Olvera, M. J. LaDu, The role of APOE in transgenic mouse models of AD. Neurosci. Lett. 707, 134285 (2019).

35. D. M. Holtzman, K. R. Bales, T. Tenkova, A. M. Fagan, M. Parsadanian, L. J. Sartorius, B. Mackey, J. Olney, D. McKeel, D. Wozniak, S. M. Paul, Apolipoprotein E isoform-dependent amyloid deposition and neuritic degeneration in a mouse model of Alzheimer’s disease. Proc. Natl. Acad. Sci. 97, 2892–2897 (2000).

36. M. P. Cadiz, K. A. Gibson, K. T. Todd, D. G. Nascari, N. Massa, M. T. Lilley, K. C. Olney, M. M. Al-Amin, H. Jiang, D. M. Holtzman, J. D. Fryer, Aducanumab anti-amyloid immunotherapy induces sustained microglial and immune alterations. J. Exp. Med. 221, e20231363 (2024).

37. G. B. D. D. F. Collaborators, E. Nichols, J. D. Steinmetz, S. E. Vollset, K. Fukutaki, J. Chalek, F. Abd-Allah, A. Abdoli, A. Abualhasan, E. Abu-Gharbieh, T. T. Akram, H. A. Hamad, F. Alahdab, F. M. Alanezi, V. Alipour, S. Almustanyir, H. Amu, I. Ansari, J. Arabloo, T. Ashraf, T. Astell-Burt, G. Ayano, J. L. Ayuso-Mateos, A. A. Baig, A. Barnett, A. Barrow, B. T. Baune, Y. Béjot, W. M. M. Bezabhe, Y. M. Bezabih, A. S. Bhagavathula, S. Bhaskar, K. Bhattacharyya, A. Bijani, A. Biswas, S. R. Bolla, A. Boloor, C. Brayne, H. Brenner, K. Burkart, R. A. Burns, L. A. Cámera, C. Cao, F. Carvalho, L. F. S. Castro-de-Araujo, F. Catalá-López, E. Cerin, P. P. Chavan, N. Cherbuin, D.-T. Chu, V. M. Costa, R. A. S. Couto, O. Dadras, X. Dai, L. Dandona, R. Dandona, V. D. l. Cruz-Góngora, D. Dhamnetiya, D. D. d. Silva, D. Diaz, A. Douiri, D. Edvardsson, M. Ekholuenetale, I. E. Sayed, S. I. El-Jaafary, K. Eskandari, S. Eskandarieh, S. Esmaeilnejad, J. Fares, A. Faro, U. Farooque, V. L. Feigin, X. Feng, S.-M. Fereshtehnejad, E. Fernandes, P. Ferrara, I. Filip, H. Fillit, F. Fischer, S. Gaidhane, L. Galluzzo, A. Ghashghaee, N. Ghith, A. Gialluisi, S. A. Gilani, I.-R. Glavan, E. V. Gnedovskaya, M. Golechha, R. Gupta, V. B. Gupta, V. K. Gupta, M. R. Haider, B. J. Hall, S. Hamidi, A. Hanif, G. J. Hankey, S. Haque, R. K. Hartono, A. I. Hasaballah, M. T. Hasan, A. Hassan, S. I. Hay, K. Hayat, M. I. Hegazy, G. Heidari, R. Heidari-Soureshjani, C. Herteliu, M. Househ, R. Hussain, B.-F. Hwang, L. Iacoviello, I. Iavicoli, O. S. Ilesanmi, I. M. Ilic, M. D. Ilic, S. S. N. Irvani, H. Iso, M. Iwagami, R. Jabbarinejad, L. Jacob, V. Jain, S. K. Jayapal, R. Jayawardena, R. P. Jha, J. B. Jonas, N. Joseph, R. Kalani, A. Kandel, H. Kandel, A. Karch, A. S. Kasa, G. M. Kassie, P. Keshavarz, M. A. B. Khan, M. N. Khatib, T. A. M. Khoja, J. Khubchandani, M. S. Kim, Y. J. Kim, A. Kisa, S. Kisa, M. Kivimäki, W. J. Koroshetz, A. Koyanagi, G. A. Kumar, M. Kumar, H. M. Lak, M. Leonardi, B. Li, S. S. Lim, X. Liu, Y. Liu, G. Logroscino, S. Lorkowski, G. Lucchetti, R. L. Saute, F. G. Magnani, A. A. Malik, J. Massano, M. M. Mehndiratta, R. G. Menezes, A. Meretoja, B. Mohajer, N. M. Ibrahim, Y. Mohammad, A. Mohammed, A. H. Mokdad, S. Mondello, M. A. A. Moni, M. Moniruzzaman, T. B. Mossie, G. Nagel, M. Naveed, V. C. Nayak, S. N. Kandel, T. H. Nguyen, B. Oancea, N. Otstavnov, S. S. Otstavnov, M. O. Owolabi, S. Panda-Jonas, F. P. Kan, M. Pasovic, U. K. Patel, M. Pathak, M. F. P. Peres, A. Perianayagam, C. B. Peterson, M. R. Phillips, M. Pinheiro, M. A. Piradov, C. D. Pond, M. H. Potashman, F. H. Pottoo, S. I. Prada, A. Radfar, A. Raggi, F. Rahim, M. Rahman, P. Ram, P. Ranasinghe, D. L. Rawaf, S. Rawaf, N. Rezaei, A. Rezapour, S. R. Robinson, M. Romoli, G. Roshandel, R. Sahathevan, A. Sahebkar, M. A. Sahraian, B. Sathian, D. Sattin, M. Sawhney, M. Saylan, S. Schiavolin, A. Seylani, F. Sha, M. A. Shaikh, K. S. Shaji, M. Shannawaz, J. K. Shetty, M. Shigematsu, J. I. Shin, R. Shiri, D. A. S. Silva, J. P. Silva, R. Silva, J. A. Singh, V. Y. Skryabin, A. A. Skryabina, A. E. Smith, S. Soshnikov, E. E. Spurlock, D. J. Stein, J. Sun, R. Tabarés-Seisdedos, B. Thakur, B. Timalsina, M. R. Tovani-Palone, B. X. Tran, G. W. Tsegaye, S. V. Tahbaz, P. R. Valdez, N. Venketasubramanian, V. Vlassov, G. T. Vu, L. G. Vu, Y.-P. Wang, A. Wimo, A. S. Winkler, L. Yadav, S. H. Y. Jabbari, K. Yamagishi, L. Yang, Y. Yano, N. Yonemoto, C. Yu, I. Yunusa, S. Zadey, M. S. Zastrozhin, A. Zastrozhina, Z.-J. Zhang, C. J. L. Murray, T. Vos, Estimation of the global prevalence of dementia in 2019 and forecasted prevalence in 2050: an analysis for the Global Burden of Disease Study 2019. Lancet Public Heal. 7, e105–e125 (2022).

38. L. Fumagalli, A. N. Mohebiany, J. Premereur, P. P. Miquel, B. Bijnens, P. V. d. Walle, N. Fattorelli, R. Mancuso, Microglia heterogeneity, modeling and cell-state annotation in development and neurodegeneration. Nat. Neurosci. 28, 1381–1392 (2025).

39. C. Escartin, E. Galea, A. Lakatos, J. P. O’Callaghan, G. C. Petzold, A. Serrano-Pozo, C. Steinhäuser, A. Volterra, G. Carmignoto, A. Agarwal, N. J. Allen, A. Araque, L. Barbeito, A. Barzilai, D. E. Bergles, G. Bonvento, A. M. Butt, W.-T. Chen, M. Cohen-Salmon, C. Cunningham, B. Deneen, B. D. Strooper, B. Díaz-Castro, C. Farina, M. Freeman, V. Gallo, J. E. Goldman, S. A. Goldman, M. Götz, A. Gutiérrez, P. G. Haydon, D. H. Heiland, E. M. Hol, M. G. Holt, M. Iino, K. V. Kastanenka, H. Kettenmann, B. S. Khakh, S. Koizumi, C. J. Lee, S. A. Liddelow, B. A. MacVicar, P. Magistretti, A. Messing, A. Mishra, A. V. Molofsky, K. K. Murai, C. M. Norris, S. Okada, S. H. R. Oliet, J. F. Oliveira, A. Panatier, V. Parpura, M. Pekna, M. Pekny, L. Pellerin, G. Perea, B. G. Pérez-Nievas, F. W. Pfrieger, K. E. Poskanzer, F. J. Quintana, R. M. Ransohoff, M. Riquelme-Perez, S. Robel, C. R. Rose, J. D. Rothstein, N. Rouach, D. H. Rowitch, A. Semyanov, S. Sirko, H. Sontheimer, R. A. Swanson, J. Vitorica, I.-B. Wanner, L. B. Wood, J. Wu, B. Zheng, E. R. Zimmer, R. Zorec, M. V. Sofroniew, A. Verkhratsky, Reactive astrocyte nomenclature, definitions, and future directions. Nat. Neurosci. 24, 312–325 (2021).

40. S. A. Liddelow, K. A. Guttenplan, L. E. Clarke, F. C. Bennett, C. J. Bohlen, L. Schirmer, M. L. Bennett, A. E. Münch, W.-S. Chung, T. C. Peterson, D. K. Wilton, A. Frouin, B. A. Napier, N. Panicker, M. Kumar, M. S. Buckwalter, D. H. Rowitch, V. L. Dawson, T. M. Dawson, B. Stevens, B. A. Barres, Neurotoxic reactive astrocytes are induced by activated microglia. Nature 541, 481–487 (2017).

41. P. Hasel, I. V. L. Rose, J. S. Sadick, R. D. Kim, S. A. Liddelow, Neuroinflammatory astrocyte subtypes in the mouse brain. Nat. Neurosci. 24, 1475–1487 (2021).

42. P. Hasel, W. H. Aisenberg, F. C. Bennett, S. A. Liddelow, Molecular and metabolic heterogeneity of astrocytes and microglia. Cell Metab. 35, 555–570 (2023).

43. V. Singhal, N. Chou, J. Lee, Y. Yue, J. Liu, W. K. Chock, L. Lin, Y.-C. Chang, E. M. L. Teo, J. Aow, H. K. Lee, K. H. Chen, S. Prabhakar, BANKSY unifies cell typing and tissue domain segmentation for scalable spatial omics data analysis. Nature Genetics 56, 431–441 (2024).

44. E. S. v. Etten, S. Mahinrad, J. D. Grill, S. Salloway, A. Atri, P. M. Cogswell, T. L. S. Benzinger, T. Iwatsubo, C. Iadecola, C. A. Lemere, J. A. R. Nicoll, S. M. Greenberg, M. C. Carrillo, C. R. Jack, R. A. Sperling, Amyloid-related imaging abnormalities (ARIA) in anti-amyloid therapies for Alzheimer’s disease: An update from the Alzheimer’s Association ARIA workgroup. Alzheimer’s Dement. 22, e71361 (2026).

45. U. K. Adhikari, R. Khan, M. Mikhael, R. Balez, M. A. David, D. Mahns, J. Hardy, M. Tayebi, Therapeutic anti-amyloid β antibodies cause neuronal disturbances. Alzheimer’s Dement. 19, 2479–2496 (2023).

46. E. Solopova, W. Romero-Fernandez, H. Harmsen, L. Ventura-Antunes, E. Wang, A. Shostak, J. Maldonado, M. Donahue, D. Schultz, T. M. Coyne, A. Charidimou, M. Schrag, Fatal Iatrogenic Cerebral Amyloid-Related Encephalitis in a patient treated with lecanemab for Alzheimer’s disease: neuroimaging and neuropathology. medRxiv, 2023.2004.2026.23289061 (2023).

47. G. A. Rodriguez, L. M. Tai, M. J. LaDu, G. W. Rebeck, Human APOE4 increases microglia reactivity at Aβ plaques in a mouse model of Aβ deposition. J. Neuroinflammation 11, 111 (2014).

48. P. Pikus, G. S. Healey, E. Xia, G. Shautidze, N. Siddapureddy, Y. Lee, C. Albanese, O. Rodriguez, R. S. Turner, G. W. Rebeck, Intravenous anti-Aβ immunotherapy acutely increases cerebral amyloid angiopathy and vascular damages in APOE4 mice. bioRxiv, 2026.2002.2009.704876 (2026).

49. H. Wang, E. S. M. Nery, P. Ardayfio, R. Khanna, D. O. Svaldi, I. Gueorguieva, S. Shcherbinin, S. W. Andersen, P. M. Hauck, S. E. Engle, D. A. Brooks, E. C. Collins, N. C. Fox, S. M. Greenberg, S. Salloway, M. A. Mintun, J. R. Sims, Modified titration of donanemab reduces ARIA risk and maintains amyloid reduction. Alzheimer’s Dement. 21, e70062 (2025).

50. P. T. Nelson, N. M. Pious, G. A. Jicha, D. M. Wilcock, D. W. Fardo, S. Estus, G. W. Rebeck, APOE-ε2 and APOE-ε4 Correlate With Increased Amyloid Accumulation in Cerebral Vasculature. J. Neuropathol. Exp. Neurol. 72, 708–715 (2013).

51. A. Biffi, A. Sonni, C. D. Anderson, B. Kissela, J. M. Jagiella, H. Schmidt, J. Jimenez-Conde, B. M. Hansen, I. Fernandez-Cadenas, L. Cortellini, A. Ayres, K. Schwab, K. Juchniewicz, A. Urbanik, N. S. Rost, A. Viswanathan, T. Seifert-Held, E.-M. Stoegerer, R. Lehner, B. Schenk, S. M. Greenberg, S. L. Silliman, R. Gill, A. Lindgren, A. Slowik, R. Schmidt, J. Montaner, B. B. Worrall, J. P. Broderick, B. M. Kissela, P. Sharma, J. Rosand, Variants at APOE influence risk of deep and lobar intracerebral hemorrhage. Ann. Neurol. 68, 934–943 (2010).

52. S. M. Greenberg, M. E. Briggs, B. T. Hyman, G. J. Kokoris, C. Takis, D. S. Kanter, C. S. Kase, M. S. Pessin, Apolipoprotein E epsilon 2 associated with vasculopathy in cerebral amyloid angiopathy. Neurology 50, 961–965 (1998).

53. S. M. Greenberg, G. W. Rebeck, J.-P. G. Vonsattel, T. Gomez-Isla, B. T. Hyman, Apolipoprotein E epsilon 2 and cerebral amyloid angiopathy dissociated from Alzheimer’s disease and associated with different hemorrhage complications. Ann. Neurol. 38, 254–259 (1995).

54. A. Charidimou, G. Boulouis, M. Pasi, E. Auriel, E. S. van Etten, K. Haley, A. Ayres, K. M. Schwab, S. Martinez-Ramirez, J. N. Goldstein, S. M. Greenberg, A. Viswanathan, APOE and cortical superficial siderosis in CAA. Neurology 93, e236–e247 (2019).

55. R. I. Mehta, T. Wang, S. E. Leurgans, D. A. Bennett, J. A. Schneider, Cerebral amyloid angiopathy and Alzheimer’s and related pathologies across APOE genotypes. Brain, (2026).

56. L. v. Olst, B. Simonton, A. J. Edwards, A. V. Forsyth, J. Boles, P. Jamshidi, T. Watson, N. Shepard, T. Krainc, B. M. R. Argue, Z. Zhang, J. Kuruvilla, L. Camp, M. Li, H. Xu, J. L. Norman, J. Cahan, R. Vassar, J. Chen, R. J. Castellani, J. A. R. Nicoll, D. Boche, D. Gate, Microglial mechanisms drive amyloid-β clearance in immunized patients with Alzheimer’s disease. Nat. Med., 1–13 (2025).

57. D. M. Wilcock, C. A. Colton, Anti-Amyloid-β Immunotherapy in Alzheimer’s Disease: Relevance of Transgenic Mouse Studies to Clinical Trials. J. Alzheimer’s Dis. 15, 555–569 (2008).

58. G. Albertini, M. Zielonka, M.-L. Cuypers, A. Snellinx, C. Xu, S. Poovathingal, M. Wojno, K. Davie, V. v. Lieshout, K. Craessaerts, L. Wolfs, E. Pasciuto, T. Jaspers, K. Horré, L. Serneels, M. Fiers, M. Dewilde, B. D. Strooper, The Alzheimer’s therapeutic Lecanemab attenuates Aβ pathology by inducing an amyloid-clearing program in microglia. Nat. Neurosci., 1–11 (2025).

59. S. Krasemann, C. Madore, R. Cialic, C. Baufeld, N. Calcagno, R. E. Fatimy, L. Beckers, E. O’Loughlin, Y. Xu, Z. Fanek, D. J. Greco, S. T. Smith, G. Tweet, Z. Humulock, T. Zrzavy, P. Conde-Sanroman, M. Gacias, Z. Weng, H. Chen, E. Tjon, F. Mazaheri, K. Hartmann, A. Madi, J. D. Ulrich, M. Glatzel, A. Worthmann, J. Heeren, B. Budnik, C. Lemere, T. Ikezu, F. L. Heppner, V. Litvak, D. M. Holtzman, H. Lassmann, H. L. Weiner, J. Ochando, C. Haass, O. Butovsky, The TREM2-APOE Pathway Drives the Transcriptional Phenotype of Dysfunctional Microglia in Neurodegenerative Diseases. Immunity 47, 566–581.e569 (2017).

60. H. Keren-Shaul, A. Spinrad, A. Weiner, O. Matcovitch-Natan, R. Dvir-Szternfeld, T. K. Ulland, E. David, K. Baruch, D. Lara-Astaiso, B. Toth, S. Itzkovitz, M. Colonna, M. Schwartz, I. Amit, A Unique Microglia Type Associated with Restricting Development of Alzheimer’s Disease. Cell 169, 1276–1290.e1217 (2017).

61. S. Lee, N. A. Devanney, L. R. Golden, C. T. Smith, J. L. Schwartz, A. E. Walsh, H. A. Clarke, D. S. Goulding, E. J. Allenger, G. Morillo-Segovia, C. M. Friday, A. A. Gorman, T. R. Hawkinson, S. M. MacLean, H. C. Williams, R. C. Sun, J. M. Morganti, L. A. Johnson, APOE modulates microglial immunometabolism in response to age, amyloid pathology, and inflammatory challenge. Cell Rep. 42, 112196 (2023).

62. M. J. LaDu, M. T. Falduto, A. M. Manelli, C. A. Reardon, G. S. Getz, D. E. Frail, Isoform-specific binding of apolipoprotein E to beta-amyloid. J. Biol. Chem. 269, 23403–23406 (1994).

63. Y. Namba, M. Tomonaga, H. Kawasaki, E. Otomo, K. Ikeda, Apolipoprotein E immunoreactivity in cerebral amyloid deposits and neurofibrillary tangles in Alzheimer’s disease and kuru plaque amyloid in Creutzfeldt-Jakob disease. Brain Research 541, 163–166 (1991).

64. G. W. Rebeck, J. S. Reiter, D. K. Strickland, B. T. Hyman, Apolipoprotein E in sporadic Alzheimer’s disease: allelic variation and receptor interactions. Neuron 11, 575–580 (1993).

65. D. A. Sanan, K. H. Weisgraber, S. J. Russell, R. W. Mahley, D. Huang, A. Saunders, D. Schmechel, T. Wisniewski, B. Frangione, A. D. Roses, W. J. Strittmatter, Apolipoprotein E associates with beta amyloid peptide of Alzheimer’s disease to form novel monofibrils: isoform apoE4 associates more efficiently than apoE3. J. Clin. Investig. 94, 860–869 (1994).

66. S. Kaji, S. A. Berghoff, L. Spieth, L. Schlaphoff, A. O. Sasmita, S. Vitale, L. Büschgens, S. Kedia, M. Zirngibl, T. Nazarenko, A. Damkou, L. Hosang, C. Depp, F. Kamp, P. Scholz, D. Ewers, M. Giera, T. Ischebeck, W. Wurst, B. Wefers, M. Schifferer, M. Willem, K.-A. Nave, C. Haass, T. Arzberger, S. Jäkel, O. Wirths, G. Saher, M. Simons, Apolipoprotein E aggregation in microglia initiates Alzheimer’s disease pathology by seeding β-amyloidosis. Immunity 57, 2651–2668.e2612 (2024).

67. Z. Xia, E. E. Prescott, A. Urbanek, H. E. Wareing, M. C. King, A. Olerinyova, H. Dakin, T. Leah, K. A. Barnes, M. M. Matuszyk, E. Dimou, E. Hidari, Y. P. Zhang, J. Y. L. Lam, J. S. H. Danial, M. R. Strickland, H. Jiang, P. Thornton, D. C. Crowther, S. Ohtonen, M. Gomez-Budia, S. M. Bell, L. Ferraiuolo, H. Mortiboys, A. Higginbottom, S. B. Wharton, D. M. Holtzman, T. Malm, R. T. Ranasinghe, D. Klenerman, S. De, Co-aggregation with apolipoprotein E modulates the function of amyloid-β in Alzheimer’s disease. Nat. Commun. 15, 4695 (2024).

68. J. D. Ulrich, T. K. Ulland, T. E. Mahan, S. Nyström, K. P. Nilsson, W. M. Song, Y. Zhou, M. Reinartz, S. Choi, H. Jiang, F. R. Stewart, E. Anderson, Y. Wang, M. Colonna, D. M. Holtzman, ApoE facilitates the microglial response to amyloid plaque pathology. J. Exp. Med. 215, 1047–1058 (2018).

69. A. Anfray, S. Schaeffer, Y. Hattori, M. M. Santisteban, N. Casey, G. Wang, M. Strickland, P. Zhou, D. M. Holtzman, J. Anrather, L. Park, C. Iadecola, A cell-autonomous role for border-associated macrophages in ApoE4 neurovascular dysfunction and susceptibility to white matter injury. Nat. Neurosci., 1–14 (2024).

70. K. Uekawa, Y. Hattori, S. J. Ahn, J. Seo, N. Casey, A. Anfray, P. Zhou, W. Luo, J. Anrather, L. Park, C. Iadecola, Border-associated macrophages promote cerebral amyloid angiopathy and cognitive impairment through vascular oxidative stress. Mol. Neurodegener. 18, 73 (2023).

71. A. Jullienne, J. I. Szu, R. Quan, M. V. Trinh, T. Norouzi, B. P. Noarbe, A. A. Bedwell, K. Eldridge, S. C. Persohn, P. R. Territo, A. Obenaus, Cortical cerebrovascular and metabolic perturbations in the 5xFAD mouse model of Alzheimer’s disease. Front. Aging Neurosci. 15, 1220036 (2023).

72. K. D. Onos, P. B. Lin, R. S. Pandey, S. A. Persohn, C. P. Burton, E. W. Miner, K. Eldridge, J. N. Kanyinda, K. E. Foley, G. W. Carter, G. R. Howell, P. R. Territo, Assessment of neurovascular uncoupling: APOE status is a key driver of early metabolic and vascular dysfunction. Alzheimer’s Dement. 20, 4951–4969 (2024).

73. J. A. K. C. Chie, S. A. Persohn, R. S. Pandey, G. W. Carter, O. R. Simcox, P. Salama, P. R. Territo, f. t. A. s. D. N. Initiative, Neurometabolic and vascular dysfunction as an early diagnostic for Alzheimer’s disease and related dementias. Alzheimer’s Dement. 21, e70790 (2025).

74. J. A. K. C. Chie, S. A. Persohn, O. R. Simcox, A. Collins, P. Salama, P. R. Territo, f. t. A. s. D. N. I. consortium, Model-Ad, Neurovascular–metabolic dysregulation, metabolic connectomics, and metabolic functional changes in Alzheimer’s disease: A preclinical and clinical comparison. Alzheimer’s Dement. 22, e71496 (2026).

75. P. Bathini, S. Schilling, J.-U. Rahfeld, D. M. Holtzman, T. C. Saido, C. A. Lemere, Early Binding of Anti-Amyloid Antibodies to CAA Drives Complement Activation, Inflammation and ARIA in Mice. bioRxiv, 2026.2003.2004.709591 (2026).

76. M. Jorfi, J. Park, C. K. Hall, C.-C. J. Lin, M. Chen, D. v. Maydell, J. M. Kruskop, B. Kang, Y. Choi, D. Prokopenko, D. Irimia, D. Y. Kim, R. E. Tanzi, Infiltrating CD8+ T cells exacerbate Alzheimer’s disease pathology in a 3D human neuroimmune axis model. Nat. Neurosci. 26, 1489–1504 (2023).

77. L. v. Olst, A. Kamermans, S. Halters, S. M. A. v. d. Pol, E. Rodriguez, I. M. W. Verberk, S. G. S. Verberk, D. W. R. Wessels, C. Rodriguez-Mogeda, J. Verhoeff, D. Wouters, J. V. d. Bossche, J. J. Garcia-Vallejo, A. W. Lemstra, M. E. Witte, W. M. v. d. Flier, C. E. Teunissen, H. E. d. Vries, Adaptive immune changes associate with clinical progression of Alzheimer’s disease. Mol. Neurodegener. 19, 38 (2024).

78. A. Ramakrishnan, N. Piehl, B. Simonton, M. Parikh, Z. Zhang, V. Teregulova, L. v. Olst, D. Gate, Epigenetic dysregulation in Alzheimer’s disease peripheral immunity. Neuron, (2024).

79. X. Chen, M. Firulyova, M. Manis, J. Herz, I. Smirnov, E. Aladyeva, C. Wang, X. Bao, M. B. Finn, H. Hu, I. Shchukina, M. W. Kim, C. M. Yuede, J. Kipnis, M. N. Artyomov, J. D. Ulrich, D. M. Holtzman, Microglia-mediated T cell infiltration drives neurodegeneration in tauopathy. Nature 615, 668–677 (2023).

80. J. Groh, R. Feng, X. Yuan, L. Liu, D. Klein, G. Hutahaean, E. Butz, Z. Wang, L. Steinbrecher, J. Neher, R. Martini, M. Simons, Microglia activation orchestrates CXCL10-mediated CD8+ T cell recruitment to promote aging-related white matter degeneration. Nat. Neurosci. 28, 1160–1173 (2025).

81. L. A. Johnson, K. Saito, A. V. Pallerla, J. L. Funnell, A. R. Ezzo, C. M. Song, D. A. Harrison, N. J. Norton, L. C. Moore, L. J. V. Eldik, D. W. Fardo, G. E. Cooper, J. M. Morganti, Clonal expansion of cytotoxic CD8⁺ T cells in lecanemab-associated ARIA. Nat. Commun., (2026).

82. M. E. Pizzo, E. D. Plowey, N. Khoury, W. Kwan, J. Abettan, S. L. DeVos, C. B. Discenza, T. Earr, D. Joy, M. Lye-Barthel, E. Roche, D. Chan, J. C. Dugas, K. Gadkar, S. Hamann, R. Meisner, J. Sebalusky, A. C. S. Amaral, I. Becerra, R. Chau, J. Chow, A. J. Clemens, M. S. Dennis, J. Duque, L. Fusaro, J. A. Getz, M. S. Kariolis, D. J. Kim, K. J. Lechtenberg, A. W.-S. Leung, A. Moshkforoush, H. N. Nguyen, E. S. Ojo, E. R. Thomsen, V. O. Torres, P. E. Sanchez, L. Shan, A. P. Silverman, Z. K. Sweeney, H. Solanoy, R. Tong, M. E. Calvert, R. J. Watts, R. G. Thorne, P. H. Weinreb, D. M. Walsh, J. W. Lewcock, T. Bussiere, Y. J. Y. Zuchero, Transferrin receptor–targeted anti-amyloid antibody enhances brain delivery and mitigates ARIA. Science 389, eads3204 (2025).

83. J. Smith, C. J. Mummery, J. L. Cummings, G. D. Rabinovici, S. Salloway, R. A. Sperling, H. Zetterberg, A. Thanasopoulou, C. Lane, P. Delmar, G. Klein, R. Croney, J. Wojtowicz, C. Hofmann, L. Kulic, H. Garren, TRONTIER 1 and TRONTIER 2: Pivotal trials of trontinemab in early symptomatic Alzheimer’s disease. Alzheimer’s Dement. 21, e104294 (2025).

84. E. H. Doud, K. Hansen, K. A. Haynes, K. Eldridge, J. Arrivalagan, N. Charbe, L. D. Silva, S. K. Quinney, A. L. Mosley, S. J. S. Rizzo, P. R. Territo, Analytical development and application of a targeted liquid chromatography-tandem mass spectrometry assay for chimeric aducanumab. mAbs 17, 2537118 (2025).

85. Q. Wang, S.-L. Ding, Y. Li, J. Royall, D. Feng, P. Lesnar, N. Graddis, M. Naeemi, B. Facer, A. Ho, T. Dolbeare, B. Blanchard, N. Dee, W. Wakeman, K. E. Hirokawa, A. Szafer, S. M. Sunkin, S. W. Oh, A. Bernard, J. W. Phillips, M. Hawrylycz, C. Koch, H. Zeng, J. A. Harris, L. Ng, The Allen Mouse Brain Common Coordinate Framework: A 3D Reference Atlas. Cell 181, 936–953.e920 (2020).

86. K. D. Onos, O. J. Marola, A. Uyar, K. J. Keezer, J. A. K. C. Chie, K. Elk, P. Bohn, A. E. Cullen, K. Eldridge, S. Persohn, J. N. Kanyinda, H. K. Kocalis, J. D. Whitesell, J. A. Harris, P. Salama, A. E. Walker, G. Carter, M. Sasner, P. R. Territo, G. R. Howell, WSB.APP/PS1 mice develop age-dependent cerebral amyloid angiopathy, cerebrovascular dysfunction, and white matter deficits. bioRxiv, 2025.2010.2008.681261 (2025).

87. R. Satija, J. A. Farrell, D. Gennert, A. F. Schier, A. Regev, Spatial reconstruction of single-cell gene expression data. Nature Biotechnology 33, 495–502 (2015).

88. A. Butler, P. Hoffman, P. Smibert, E. Papalexi, R. Satija, Integrating single-cell transcriptomic data across different conditions, technologies, and species. Nature Biotechnology 36, 411–420 (2018).

89. Y. Hao, S. Hao, E. Andersen-Nissen, W. M. Mauck, S. Zheng, A. Butler, M. J. Lee, A. J. Wilk, C. Darby, M. Zagar, P. Hoffman, M. Stoeckius, E. Papalexi, E. P. Mimitou, J. Jain, A. Srivastava, T. Stuart, L. B. Fleming, B. Yeung, A. J. Rogers, J. M. McElrath, C. A. Blish, R. Gottardo, P. Smibert, R. Satija, Integrated analysis of multimodal single-cell data. Cell 184, 3573–3587 (2021).

90. Y. Hao, T. Stuart, M. H. Kowalski, S. Choudhary, P. Hoffman, A. Hartman, A. Srivastava, G. Molla, S. Madad, C. Fernandez-Granda, R. Satija, Dictionary learning for integrative, multimodal and scalable single-cell analysis. Nature Biotechnology 42, 293–304 (2023).

91. T. Stuart, A. Butler, P. Hoffman, C. Hafemeister, E. Papalexi, W. M. Mauck, Y. Hao, M. Stoeckius, P. Smibert, R. Satija, Comprehensive Integration of Single-Cell Data. Cell 177, 1888–1902 (2019).

92. L. McInnes, J. Healy, J. Melville, UMAP: Uniform Manifold Approximation and Projection for Dimension Reduction. arXiv abs/1802.03426, (2018).

93. S. Jin, C. F. Guerrero-Juarez, L. Zhang, I. Chang, R. Ramos, C.-H. Kuan, P. Myung, M. V. Plikus, Q. Nie, Inference and analysis of cell-cell communication using CellChat. Nat. Commun. 12, 1088 (2021).

94. S. Jin, M. V. Plikus, Q. Nie, CellChat for systematic analysis of cell-cell communication from single-cell transcriptomics. Nature Protocols 20, 180–219 (2025).

95. P. L. Germain, A. Lun, C. Garcia Meixide, W. Macnair, M. D. Robinson, Doublet identification in single-cell sequencing data using scDblFinder. F1000Res 10, 979 (2021).

96. J. A. Ramilowski, T. Goldberg, J. Harshbarger, E. Kloppmann, M. Lizio, V. P. Satagopam, M. Itoh, H. Kawaji, P. Carninci, B. Rost, A. R. R. Forrest, A draft network of ligand-receptor-mediated multicellular signalling in human. Nat. Commun. 6, 7866 (2015).

97. N. Borcherding, A. Vishwakarma, A. P. Voigt, A. Bellizzi, J. Kaplan, K. Nepple, A. K. Salem, R. W. Jenkins, Y. Zakharia, W. Zhang, Mapping the immune environment in clear cell renal carcinoma by single-cell genomics. Communications Biology 4, 122 (2021).

98. M. Ashburner, C. A. Ball, J. A. Blake, D. Botstein, H. Butler, J. M. Cherry, A. P. Davis, K. Dolinski, S. S. Dwight, J. T. Eppig, M. A. Harris, D. P. Hill, L. Issel-Tarver, A. Kasarskis, S. E. Lewis, J. C. Matese, J. E. Richardson, M. Ringwald, G. M. Rubin, G. Sherlock, Gene ontology: tool for the unification of biology. Nature Genetics 25, 25–29 (2000).

99. I. Dolgalev. (CRAN, 2025).

100. C. Gene Ontology, S. A. Aleksander, J. Balhoff, S. Carbon, J. M. Cherry, H. J. Drabkin, D. Ebert, M. Feuermann, P. Gaudet, N. L. Harris, D. P. Hill, The Gene Ontology knowledgebase in 2023. Genetics 224, iyad031 (2023).

101. B. Jassal, L. Matthews, G. Viteri, C. Gong, P. Lorente, A. Fabregat, K. Sidiropoulos, J. Cook, M. Gillespie, R. Haw, F. Loney, B. May, M. Milacic, K. Rothfels, C. Sevilla, V. Shamovsky, S. Shorser, T. Varusai, J. Weiser, G. Wu, L. Stein, H. Hermjakob, P. D’Eustachio, The reactome pathway knowledgebase. Nucleic Acids Research 48, D498–D503 (2020).

102. G. Korotkevich, V. Sukhov, N. Budin, B. Shpak, M. N. Artyomov, A. Sergushichev, Fast gene set enrichment analysis. bioRxiv, (2021).

103. A. Liberzon, C. Birger, H. Thorvaldsdottir, M. Ghandi, J. P. Mesirov, P. Tamayo, The Molecular Signatures Database (MSigDB) hallmark gene set collection. Cell Systems 1, 417–425 (2015).

104. A. Subramanian, P. Tamayo, V. K. Mootha, S. Mukherjee, B. L. Ebert, M. A. Gillette, A. Paulovich, S. L. Pomeroy, T. R. Golub, E. S. Lander, J. P. Mesirov, Gene set enrichment analysis: a knowledge-based approach for interpreting genome-wide expression profiles. Proceedings of the National Academy of Sciences of the United States of America 102, 15545–15550 (2005).

105. M. Bilous, D. Buszta, J. Bac, S. Kang, Y. Dong, S. Tissot, S. Andre, M. Alexandre-Gaveta, C. Voize, S. Peters, K. Homicsko, R. Gottardo, Resolving sensitivity, specificity and signal contamination in Xenium spatial transcriptomics. Nature Methods, (2026).

106. D. M. Cable, E. Murray, L. S. Zou, A. Goeva, E. Z. Macosko, F. Chen, R. A. Irizarry, Robust decomposition of cell type mixtures in spatial transcriptomics. Nature Biotechnology 40, 517–526 (2022).

107. K. Saito, D. S. Goulding, G. L. Nolt, S. H. Dimas, L. C. Moore, I. O. Stevens, S. Anderson, A. Snipes, S. L. Macauley, P. T. Nelson, L. A. Johnson, J. M. Morganti, High-Resolution Spatial Profiling of Microglia Reveals Proximity Associated Immunometabolic Reprogramming in Alzheimer’s Disease. bioRxiv, 2025.2005.2016.654329 (2025).

108. B. Phipson, C. B. Sim, E. R. Porrello, A. W. Hewitt, J. Powell, A. Oshlack, propeller: testing for differences in cell type proportions in single cell data. Bioinformatics 38, 4720–4726 (2022).

