## Supplemental Figures for "Anti-amyloid immunotherapy drives APOE4 specific increases in glial reactivity, perivascular immune activation, and ARIA-like events"

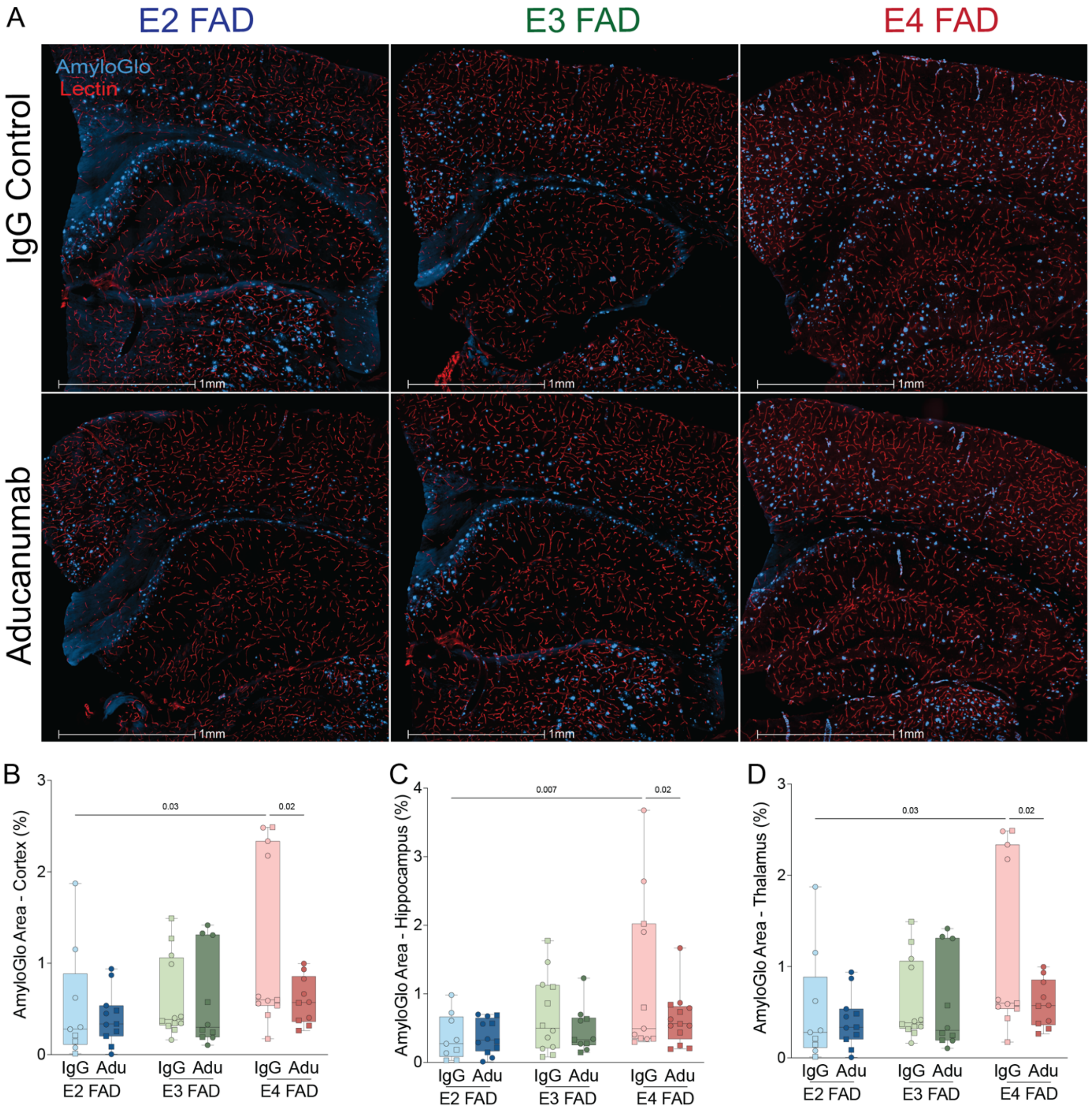


**Supplementary Figure 1:** Regional quantification of amyloid burden. A) Representative image of amyloid plaque staining (AmyloGlo, blue) and vessels (Isolectin, red). B-D) Quantification of amyloid burden in cortex, hippocampus, and thalamus, respectively as percent area coverage.

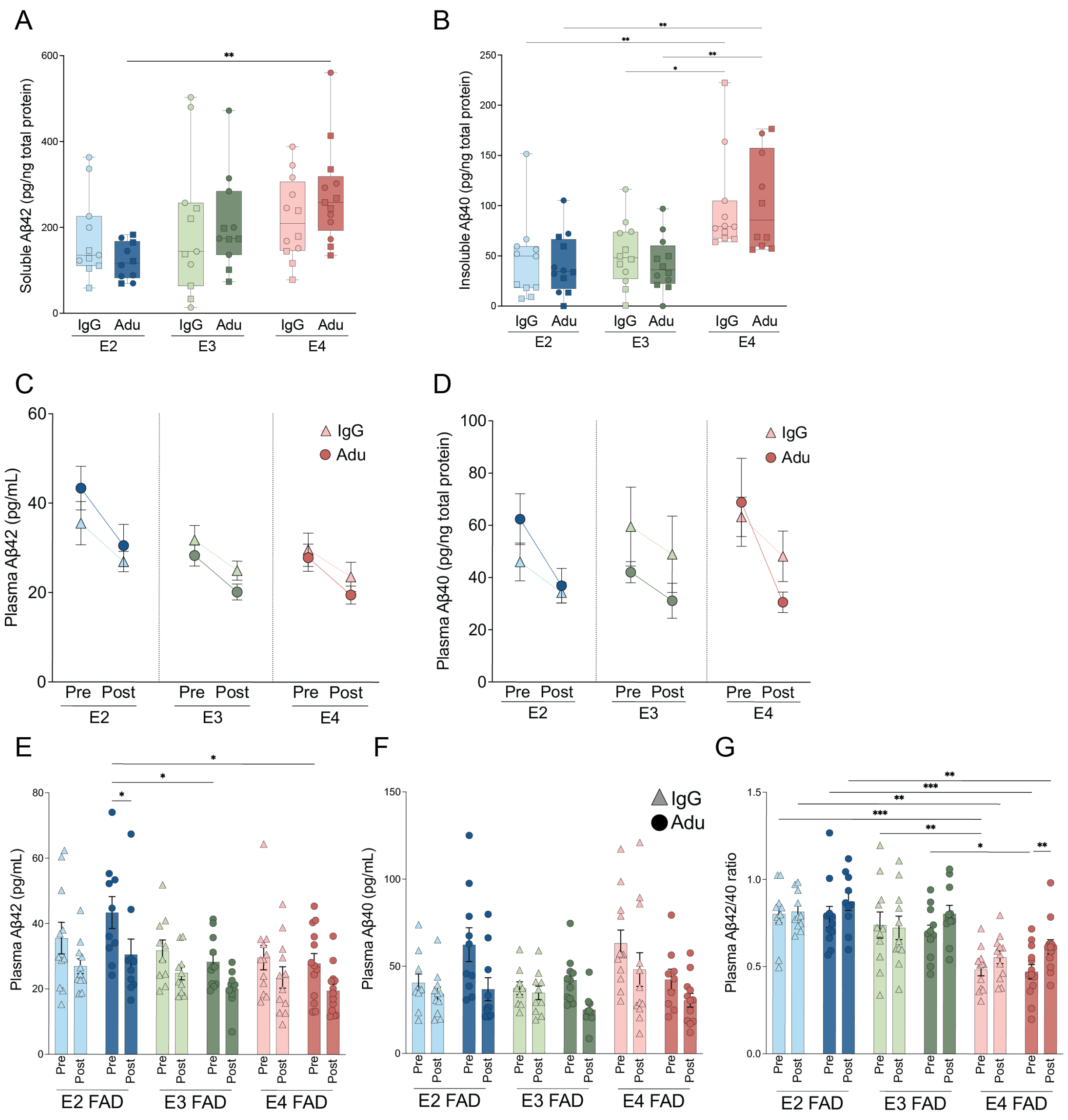


**Supplementary Figure 2:** ELISA quantification of Aβ40 and 42 in plasma and brain tissue. A-B) Quantification of Aβ42 and Aβ40 in insoluble brain fraction. C-D) Quantification of plasma Aβ42 and plasma Aβ40 over treatment course. E-G) Analysis breakdown of pre-post treatment plasma Aβ42, Aβ40, Aβ42/40 ratio quantification, respectively.

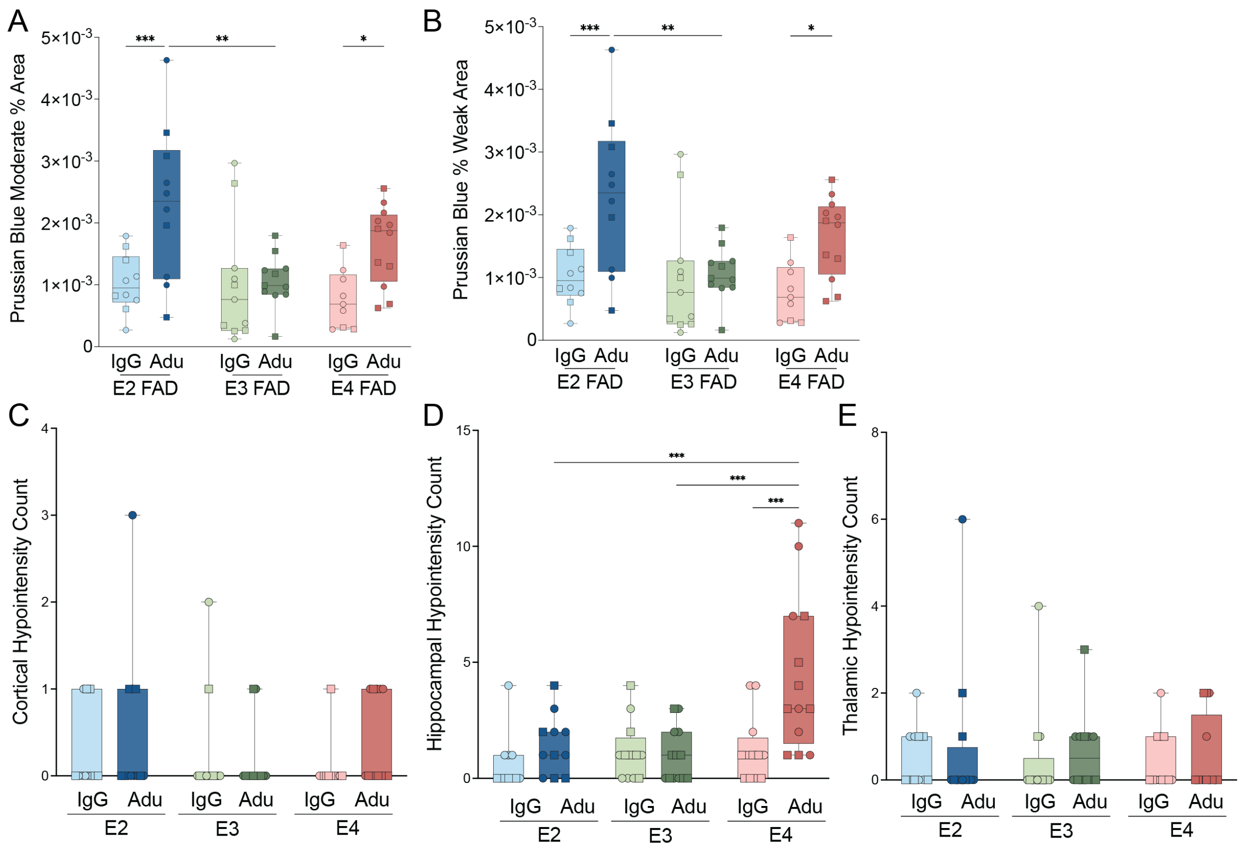


**Supplementary Figure 3:** Additional quantifications of Prussian Blue and regional hypointensity occurrence on MRI. A-B) Quantification of Prussian blue moderate area (A) and weak area (B). C-E) Hypointensity occurrence in cortex, hippocampus, and thalamus, respectively.

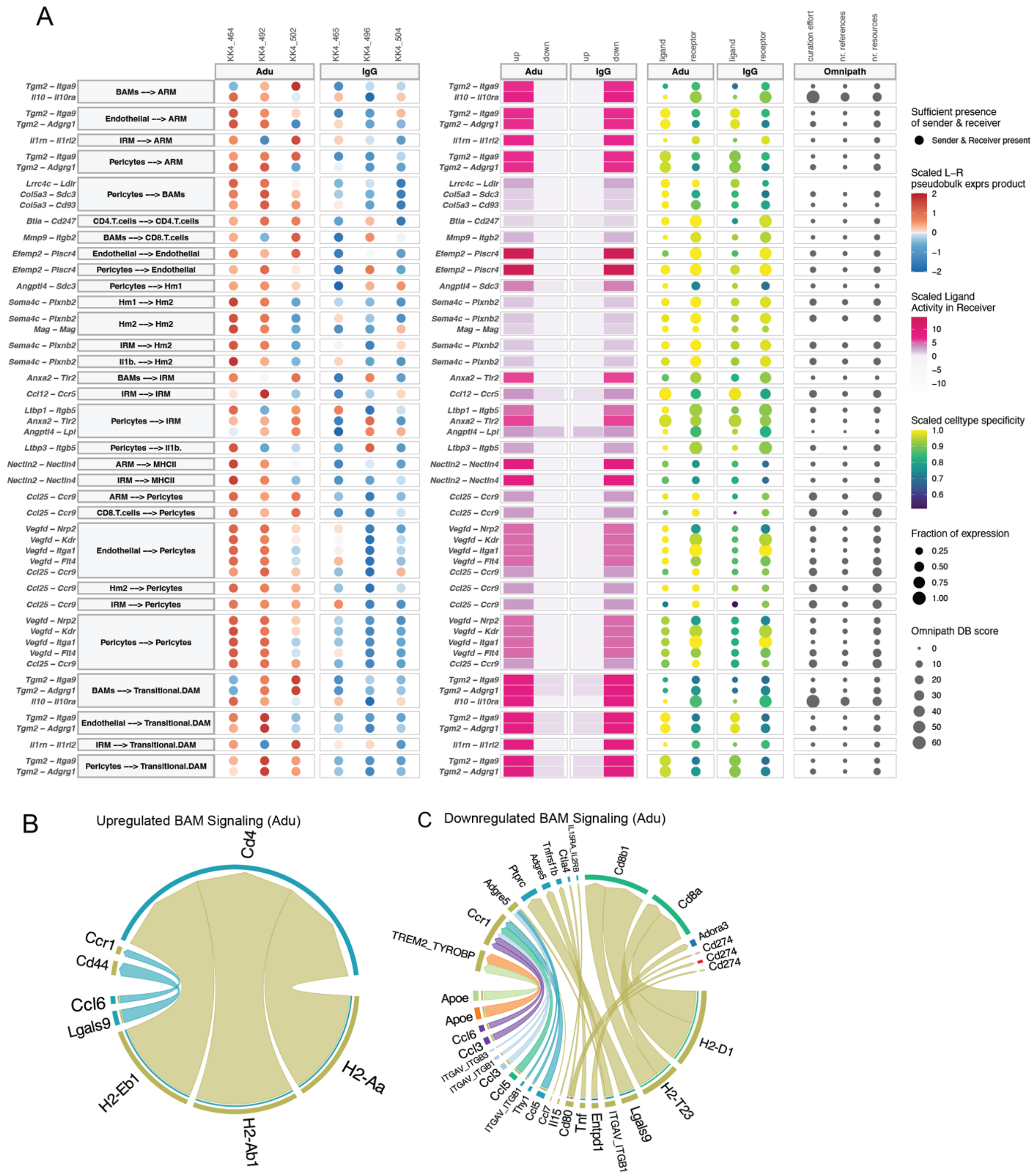


**Supplementary Figure 4:** MultiNicheNet and CellChat signaling between vascular and immune cells. A) MultiNicheNet diagram depicting signaling pathways up/downregulated in single cell RNA sequencing samples with pathway and celltypes on left, regulation status in the middle and cellular specificity on right. B-C) Up and downregulated signaling after chAdu treatment in border associated macrophages.

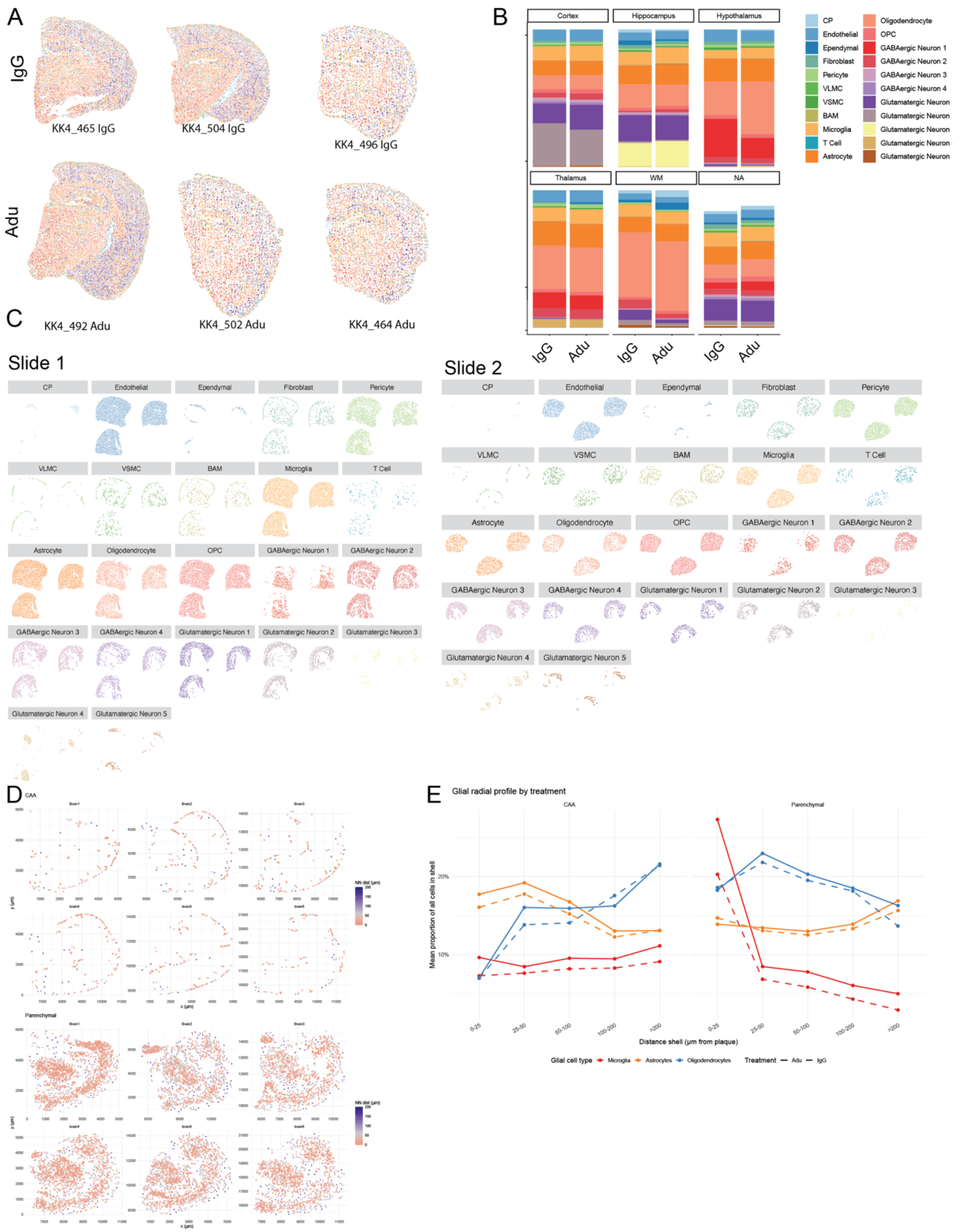


**Supplementary Figure 5:** Xenium data characterization. A) Representative images of 6 brains used for Xenium ST. B) Cell type proportions split by treatment and region. C) Cell type spatial distribution breakdown. D) Spatial distribution of plaques in each brain sample. E) Composition of peri-plaque and peri-caa niche in terms of microglia, astrocytes, and oligodendrocytes by distance.

Supplementary Figure 6

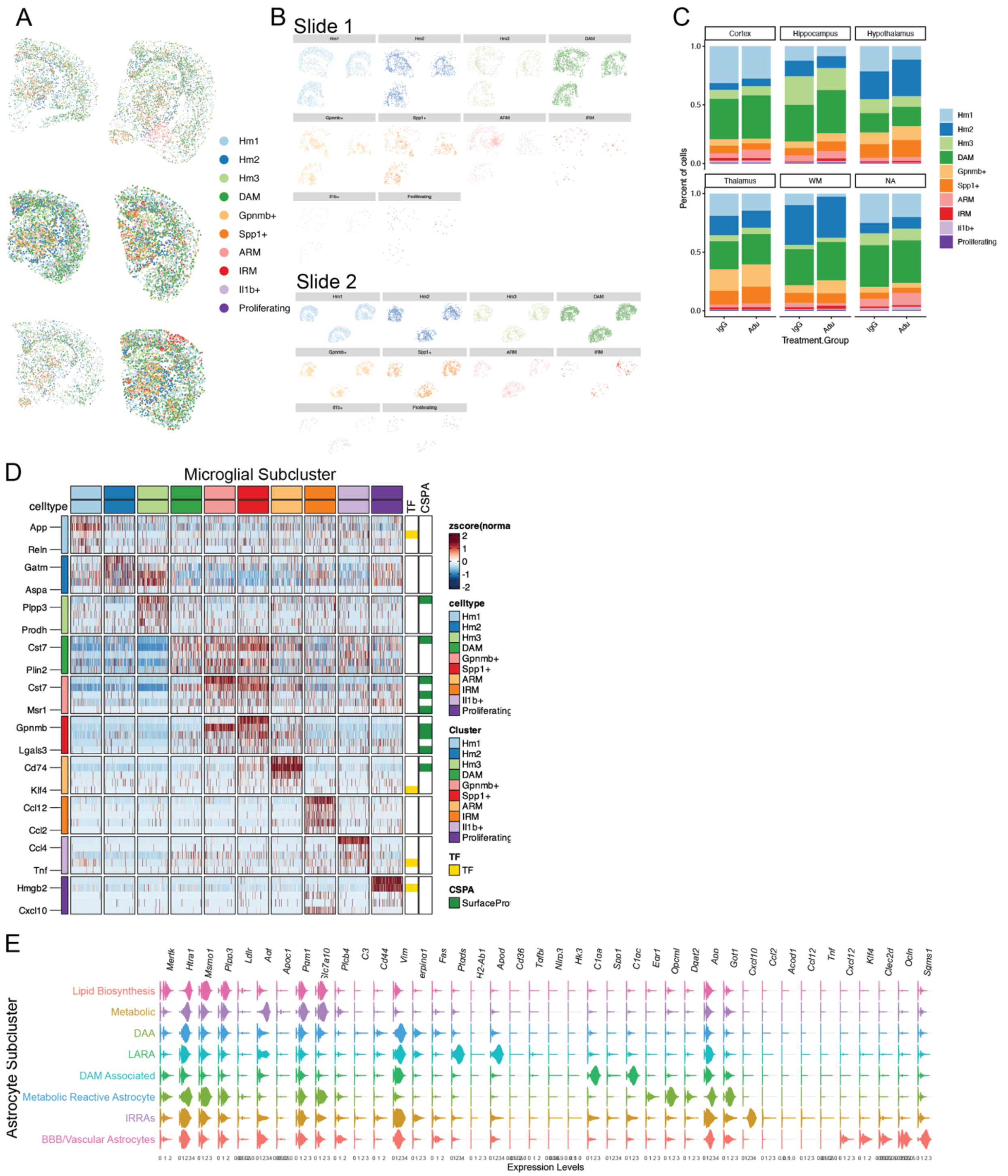


**Supplementary Figure 6:** Breakdown of Astrocytes and microglia in Xenium spatial transcriptomics and cellular annotations. A) Representative image of Microglial breakdown. B) Microglial subcluster spatial breakdown. C) Relative proportions of microglial subclusters by region and treatment group. D) expression of canonical microglial state markers by subcluster. E) Expression of canonical astrocyte state markers by subcluster.

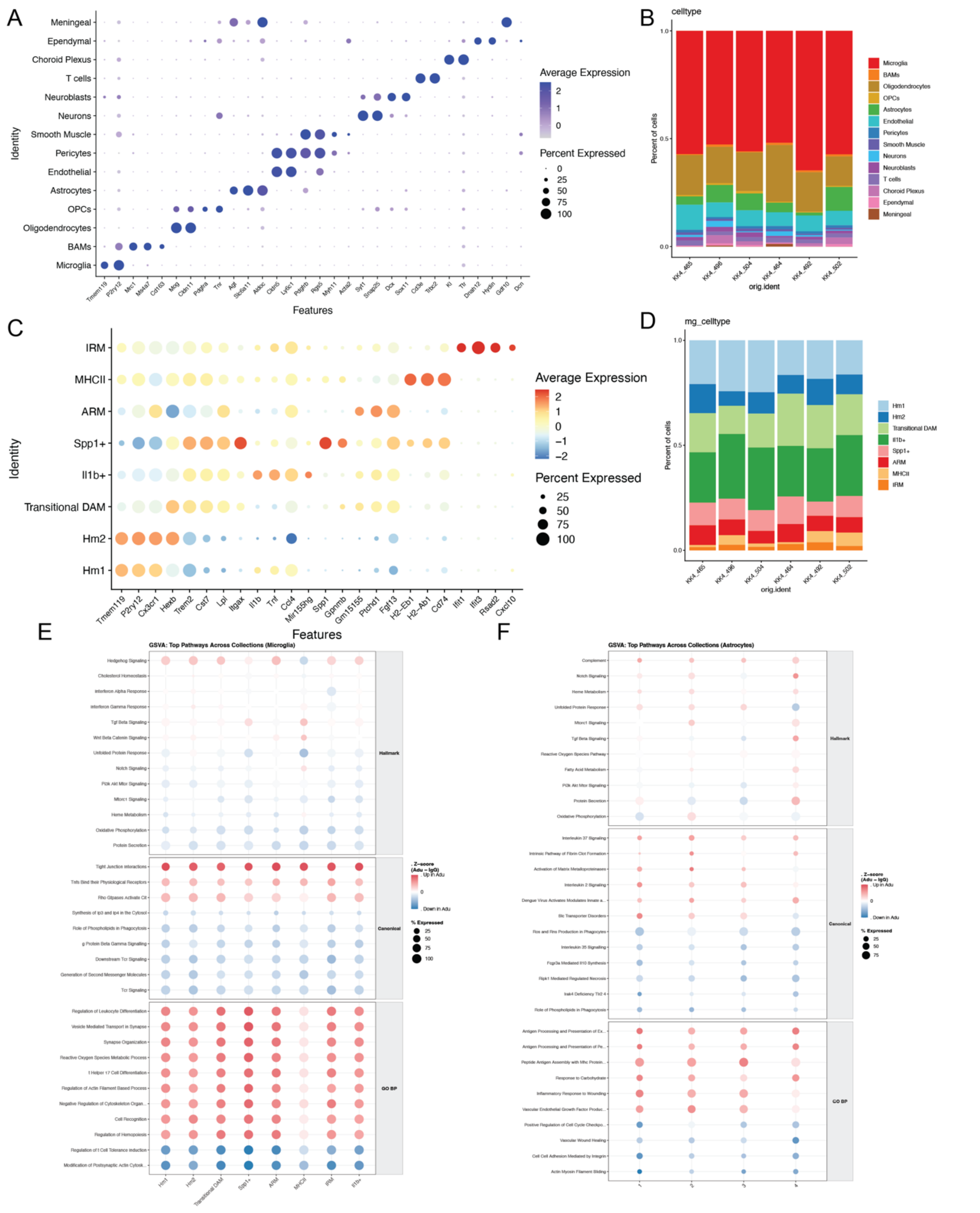

**Supplementary Figure 7:** Cell type annotations in scRNAseq data and Gene set variation analysis. A) Cell type x canonical marker matrix dotplot. B) Distribution of celltypes in scRNAseq data. C) Microglial subcluster x canonical marker matrix dotplot. D) Proportions of microglial subclusters in scRNAseq data. E) Gene set variation analysis across 3 pathway sets in microglia (E) and astrocytes (F).

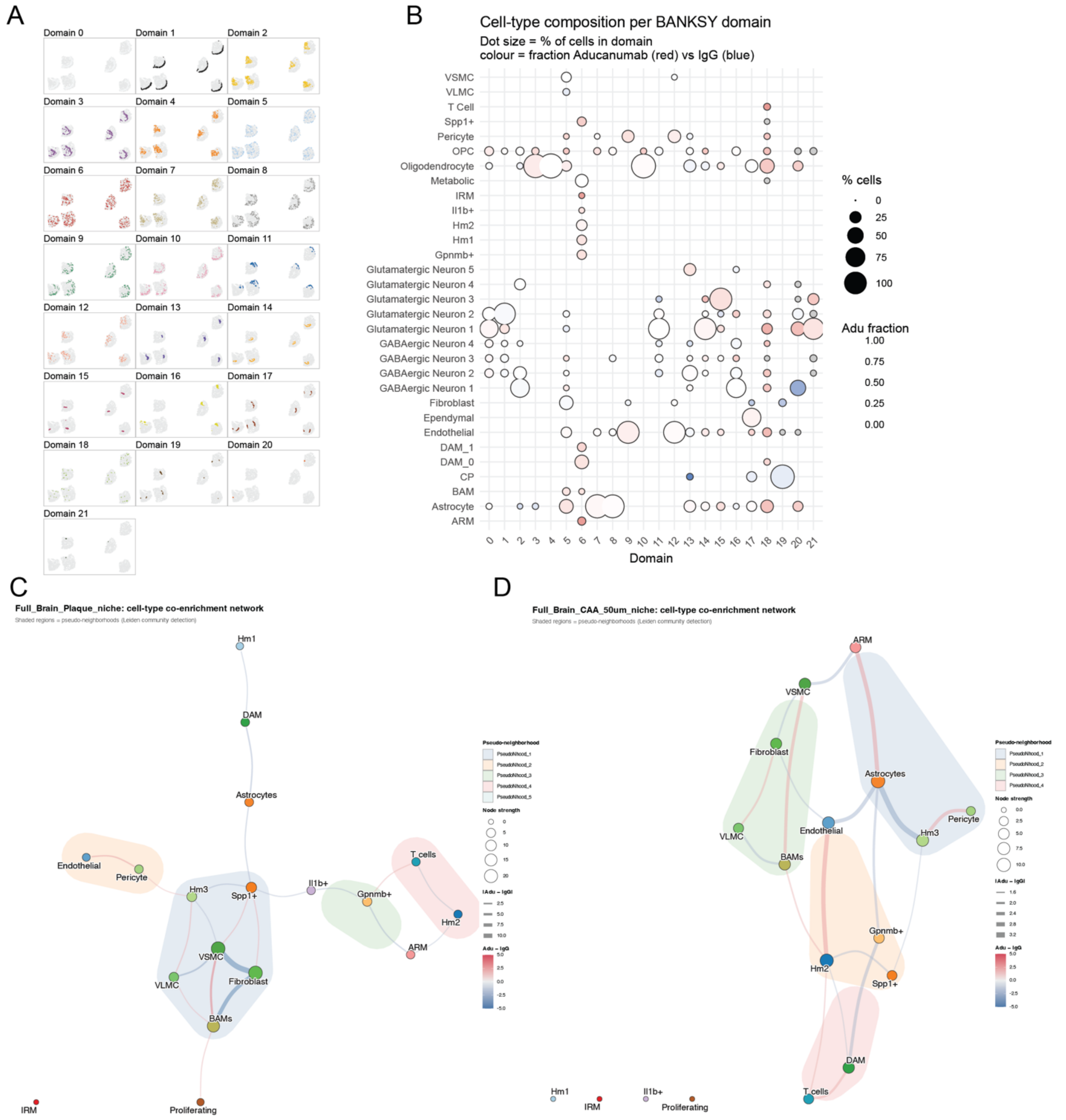


**Supplementary Figure 8:** BANKSY Domain compositions and Networks. A) Spatial distribution of BANKSY domains. B) Cell type composition of BANKSY domains. C-D) Spatial neighborhood network plot of parenchymal plaque spatial neighborhood (C) and CAA spatial neighborhood (D).
